# Iterative Bump-and-hole engineering creates a bioorthogonal reporter for *N*-acetylglucosaminyltransferase I

**DOI:** 10.64898/2026.01.16.699845

**Authors:** Yu Liu, Saskia Pieters, Ganka Bineva-Todd, Mert Sagiroglugil, Sean A. Burnap, Freya Hoddle, Anna Cioce, Andre Ohara, Kevin Bruemmer, Carolyn R. Bertozzi, Karen Polizzi, Weston B. Struwe, Carme Rovira, Benjamin Schumann

**Author notes:** These authors contributed equally (Yu Liu, Saskia Pieters).

## Abstract

Asparagine-linked protein glycosylation is among the most frequent modifications of proteins trafficking through the secretory pathway. These glycans are manufactured in an assembly line process to a common precursor that is then subject to individual modifications with different levels of complexity. An important biosynthetic modulator is the incorporation of *N*-acetylglucosamine (GlcNAc) at distinct positions in N-linked glycan biosynthesis, commencing with the activity of the glycosyltransferase MGAT1. While mapping of N-glycans to their corresponding protein attachment sites is generally possible, not much is known about the glycoprotein substrate choice for MGAT1 and related transferases. Analogs of GlcNAc with small bioorthogonal tags can be incorporated into N-glycans. However, due to the promiscuity of some GlcNAc transferases, incorporation is of little specificity towards individual positions. Here, we report an iterative bump-and-hole approach in the design of a bioorthogonal precision tool for the activity of MGAT1 in mammalian cells. Structure-informed protein engineering abrogated the activity of MGAT1 towards the nucleotide-sugar UDP-GlcNAc while retaining activity towards bumped, azide-modified analogs. Kinetic and computational analyses using a neural network approach informed the synthesis of a tailored UDP-GlcNAc analog with preferential acceptance by the engineered enzyme. Following substrate biosynthesis, the strategy allowed selective incorporation of a chemical tag on MGAT1 substrate proteins in living mammalian cells with little background incorporation by other GlcNAc transferases. Our work expands the toolbox for glycan-based reporter compounds.

## Introduction

Asparagine (N)-linked glycosylation is one of the most abundant posttranslational modifications with profound disease relevance. More than 100 congenital disorders of glycosylation are associated with defects in N-glycan biosynthesis, commensurate with their roles in protein maturation, stability, and function.^1, 2^ Early biosynthesis features transfer of a pre-assembled oligosaccharide to protein substrates before subsequent trimming and glycosylation events take effect.^3^ These elaboration events bear resemblance to an assembly line, with bifurcation points leading to distinct biosynthetic fates (Fig. 1A). The sequential addition of the sugar *N*-acetylglucosamine (GlcNAc) distinguishes between N-glycan subtypes, introduced from the uridine diphospho (UDP)-GlcNAc donor by a set of structurally diverse GlcNAc transferase enzymes.^4–6^ The determinants of differential N-glycan elaboration have been a longstanding subject of investigation.^4,7,8^ Unraveling the mechanistic details necessitates tools for detection of transferase activity, especially for the GlcNAc-inserting bifurcation points in N-glycan assembly.

**Fig. 1:**
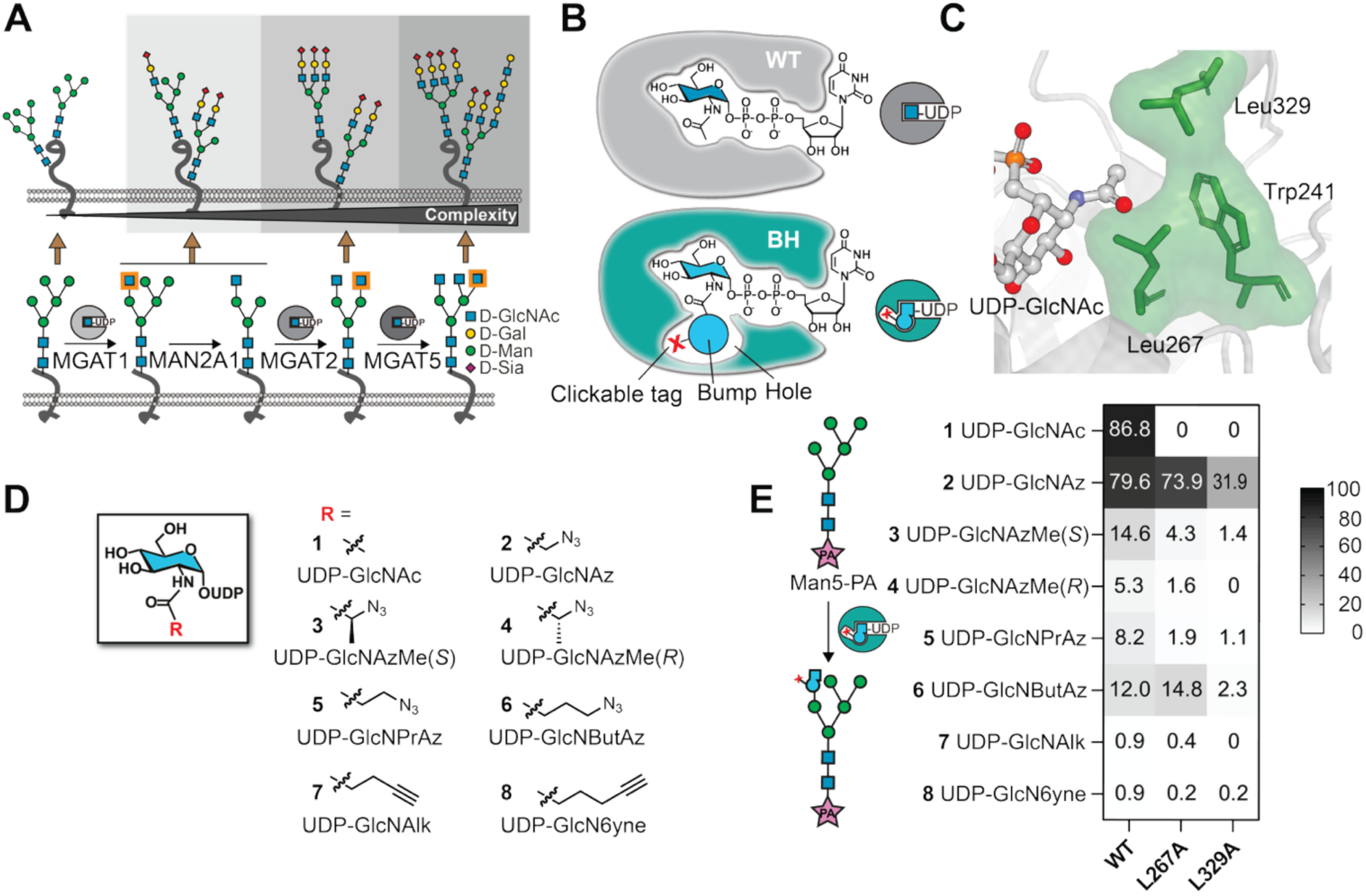
Protein engineering to restrict the activity of MGAT1 towards UDP-GlcNAc. *A*, successive elaboration of a pentamannosylated N-glycan precursor with GlcNAc by the enzymes MGAT1, MAN2A1, MGAT2 and MGAT5. GlcNAc moieties newly added in each reaction are highlighted. *B*, engineering of WT-MGAT1 to yield a BH variant that binds a bumped UDP-GlcNAc analog. *C*, gatekeeper residues of MGAT1 within close proximity to the acetamide moiety of UDP-GlcNAc (PDB: 1FOA). Annotation of gatekeeper residues is made based on the human enzyme. Gatekeepers in the original rabbit MGAT1 structure are Trp243, Leu269 and Leu331. *D*, collection of bumped, bioorthogonal UDP-GlcNAc analogs.^40^ *E*, enzymatic turnover using recombinant MGAT1 and a procainamide (PA)-labelled pentamannosylated glycan as acceptor (30-min reaction) as assessed by UHPLC with detection by absorption at 302 nm (**1** and analogs **3-8**) or mass spectrometry (analog **2**) (see Supporting Fig. 2 for representative traces). Data are means from two independent replicates. See Supporting Fig. 3 for individual data points.

GlcNAc transferase I, termed GnT-I or MGAT1, is necessary for the elaboration of all hybrid and complex N-glycans.^9^ The enzyme forms a β-1,2-glycosidic bond between GlcNAc and a mannose on the α-1,3-arm of a pentamannosylated N-glycan precursor, Man_5_GlcNAc_2_-Asn (Fig. 1A).^4,10^ Subsequent biosynthetic steps feature glycan trimming by the mannosidase MAN2A1 and GlcNAc addition by other transferases such as MGAT2 and MGAT5 to furnish more complex glycan structures.^9^ MGAT1 is a type II transmembrane protein with catalytic activity in the Golgi lumen.^11,12^ Expression of *Mgat1* is essential in mice, and inactivating mutations are associated with smaller embryo size, inverted left-right asymmetry, impaired vascularization, and death at embryonic day 9.5.^13,14,15^ While structurally related to other GlcNAc transferases, MGAT1 bears sequence homology to only a small number of other human enzymes within the glycosyltransferase (GT) family 13.^16^ Investigating the decisive role of MGAT1 on the biosynthetic fate of N-glycans is of fundamental importance. Genetic engineering has underpinned existing efforts, for instance through the Lec1 Chinese hamster ovary (CHO) cell line that lacks endogenous MGAT1 activity.^17,18^ Yet, we lack methods to directly visualize and profile the substrates of MGAT1 as a crucial bifurcation point in N-glycan biosynthesis.

Bioorthogonal tools have revolutionized the field of glycobiology. Monosaccharides with azide, alkyne or other bioorthogonal groups can be used by cellular or artificial biosynthetic machineries to be turned into nucleotide-sugars and accepted as substrates by GT enzymes.^19–21^ Bioorthogonal derivatives of GlcNAc have been extensively studied, starting with the azide-displaying monosaccharide *N*-azidoacetyl-D-glucosamine GlcNAz and applied to a myriad of related compounds.^22–27^ These analogs can be incorporated into many cellular glycoconjugates including N-glycosylated proteins.^28–33^ A function of their incorporation is the biosynthesis of the corresponding nucleotide-sugars that can be boosted by expression of heterologous or engineered metabolic enzymes. For instance, the monosaccharide kinase NahK from *Bifidobacterium longum* and engineered versions of the human pyrophosphorylases AGX1/2 enhance biosynthesis of chemically tagged UDP-sugar analogs.^30,31,34–36^

We reasoned that a reporter tool selective for MGAT1 would allow visualization of N-glycosylated proteins, with eventual applications in dissecting glycan biosynthesis in disease-relevant systems. A strategy termed bump-and-hole (BH) engineering can be employed to this end, by creating a bioorthogonal, “bumped” UDP-sugar that is selectively used by an engineered GT of interest.^28,37–39^ Engineering introduces a “hole” in the active site that accommodates the “bump” (Fig. 1B). Ideally, the BH enzyme-substrate pair selectively modifies glycoproteins without substantial background incorporation by any wild-type (WT) GT. We recently reported a BH approach for the activity of the GlcNAc transferase MGAT5, employing UDP-*N*-4-azidobutanoyl-D-glucosamine (UDP-GlcNButAz) out of a collection of nine bioorthogonal, bumped UDP-GlcNAc analogs.^40^ This work highlighted the substrate promiscuity of GlcNAc transferases towards chemically modified UDP-GlcNAc analogs, with BH engineering primarily serving to restrict activity towards the native substrate UDP-GlcNAc. Mukherjee, Hanover and colleagues recently reported a bioorthogonal UDP-GlcNAc derivative with a 6-toluenesulfonyl (OTs) group that exerted selectivity for MGAT1 and tagged glycoproteins in mammalian cells.^41^ The corresponding analog was fed to cells at concentrations above 100 µM and a propionyl ester caging group was found to remain on the GlcNAc analog after incorporation. We reasoned that a structurally-inspired, bottom-up probe discovery approach would deliver a complementary, selective reporter for MGAT1 activity while allowing for tuneable biosynthesis by UDP-GlcNAc salvage pathways. The availability of a bioorthogonal, bumped UDP-GlcNAc analog collection and available crystal structures fuelled a BH approach for MGAT1.^12,42^

## Results and Discussion

### Structurally informed development of an MGAT1 bump-and-hole enzyme-substrate pair

We used a published crystal structure of rabbit MGAT1 to identify the so-called gatekeeper residues that are large and hydrophobic, and within 8-9 Å of the UDP-GlcNAc acetamide (Fig. 1C).^12, 42^ Trp241, Leu267, and Leu329 in human MGAT1 fulfilled these criteria. Employing an MGAT1 secretion construct as a template, site-directed mutagenesis was used to generate seven constructs in which the gatekeeper residues Leu267 and Leu329 were primarily substituted with smaller Ala residues, with additional substitutions of Trp241 with His. Recombinant enzymes were expressed in Expi293 cells and purified by Ni-NTA affinity chromatography (Supporting Fig. 1).

*In vitro* enzymatic turnover was performed to assess the ability of MGAT1 constructs to accept the modified UDP-GlcNAc analogs. We subjected all seven enzymes to glycosylation reactions with either UDP-GlcNAc **1** or each of our seven previously synthesized, bumped UDP-GlcNAc analogs **2**-**8**.^40^ The analogs contained hydrophobic modifications as well as bioorthogonal azides or alkynes for downstream application (Fig. 1D). A procainamide (PA)-labelled, pentamannosylated N-glycan derivative termed Man5-PA was used as an acceptor substrate in all enzymatic assays (Fig. 1E). Incorporation of a GlcNAc analog by MGAT1 into Man5-PA allowed separation by ultra-high-performance liquid chromatography (UHPLC) with either UV or mass spectrometric detection to calculate turnover (Supporting Fig. 2). WT-MGAT1 exhibited enzymatic activity against UDP-GlcNAc **1** and azide-containing analogs **2**-**6**, but not alkyne-containing analogs **7** or **8**. Out of the six MGAT1 variants tested, only the Leu267Ala and Leu329Ala variants exhibited enzymatic activity in a 30*-*min reaction (Fig. 1E, Supporting Fig. 3). Both accepted UDP-GlcNAz **2** with the smallest bioorthogonal modification as the best substrate, while crucially, acceptance of UDP-GlcNAc **1** was lost in both variants. The Leu267Ala variant displayed similar activity with the substrate UDP-GlcNButAz **6** as WT-MGAT1 at 14.8% and 12% incorporation, respectively. These data suggest that the Leu267Ala variant, called BH-MGAT1, is a suitable enzyme for tracing the activity of MGAT1 due to its loss of acceptance of the native substrate UDP-GlcNAc **1**. We note that this behaviour of MGAT1 bears resemblance to MGAT5, where BH engineering primarily led to loss of UDP-GlcNAc acceptance.^40^

**Fig. 2:**
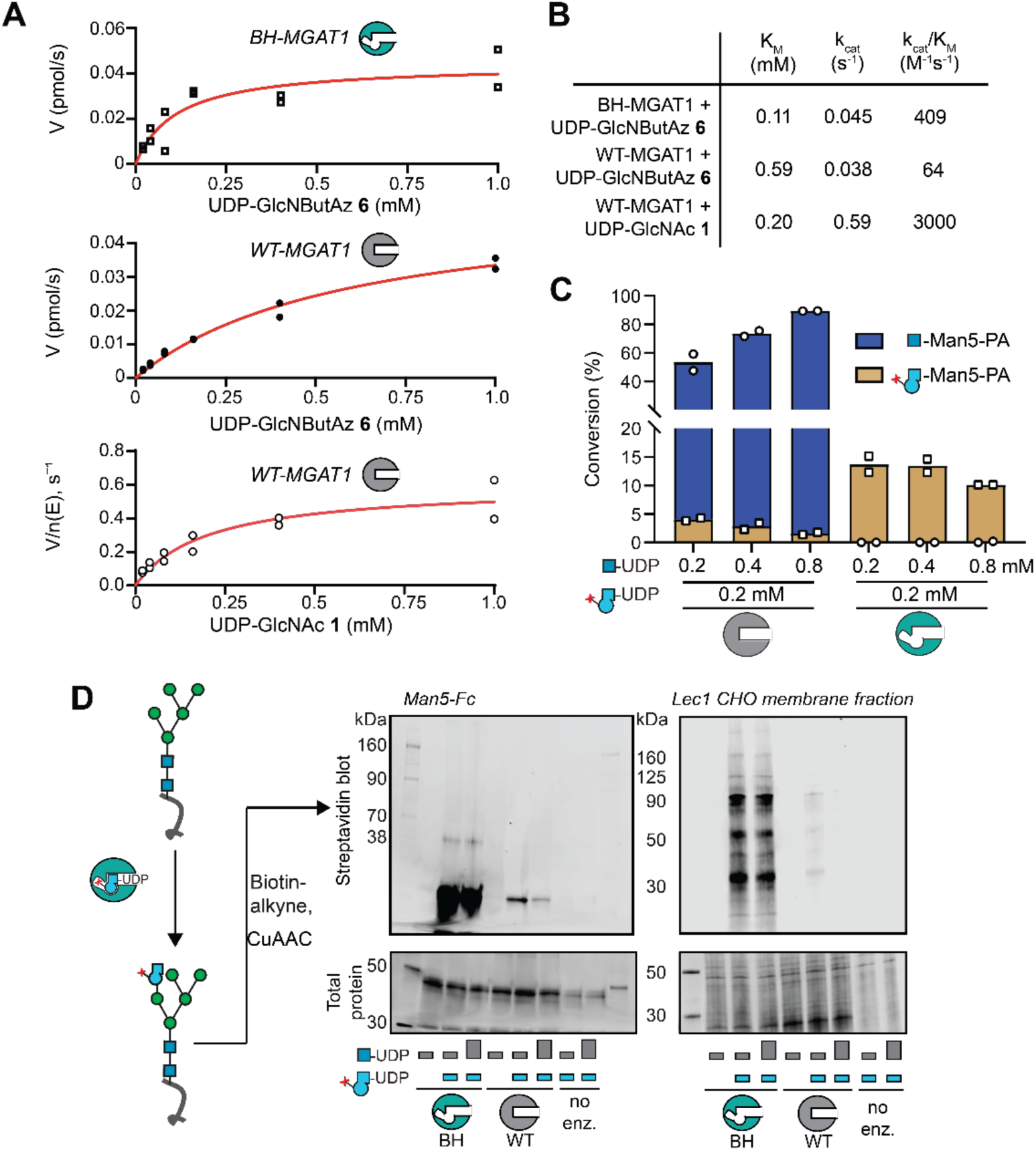
Evaluation of the BH-MGAT1/**6** enzyme-substrate pair. *A*, Michaelis-Menten kinetic experiment of either BH-MGAT1 or WT-MGAT1 with UDP-GlcNButAz **6**. Data are individual values of two independent experiments. Data were fitted with a Michaelis-Menten model (red line). *B*, calculated kinetic parameters for enzyme-substrate pairs measured through curve fitting in *A*. *C*, competition experiment between the use of UDP-GlcNAc and UDP-GlcNButAz **6**. Both UDP-sugars were supplied in 30-min *in vitro* glycosylation experiments in depicted concentrations and individual reaction products were quantified by UHPLC. Data are individual datapoints and means from two independent replicates. *D*, *In vitro* glycosylation of Man5-containing glycoprotein substrates by BH-engineered MGAT1. Glycoproteins were treated with WT- or BH-MGAT1 and the substrates UDP-GlcNAc (grey bars) and UDP-GlcNButAz **6** (blue bars) in either 0.2 mM (small bars) or 0.8 mM (big bars) concentration. *In vitro* glycosylation was applied to Man5-Fc (left) or a Lec1 CHO cell membrane fraction (right). Reaction mixtures were treated with biotin-alkyne under CuAAC conditions and bioorthogonal tagging analyzed by streptavidin blot. Data are from one out of two independent replicates.

**Fig. 3:**
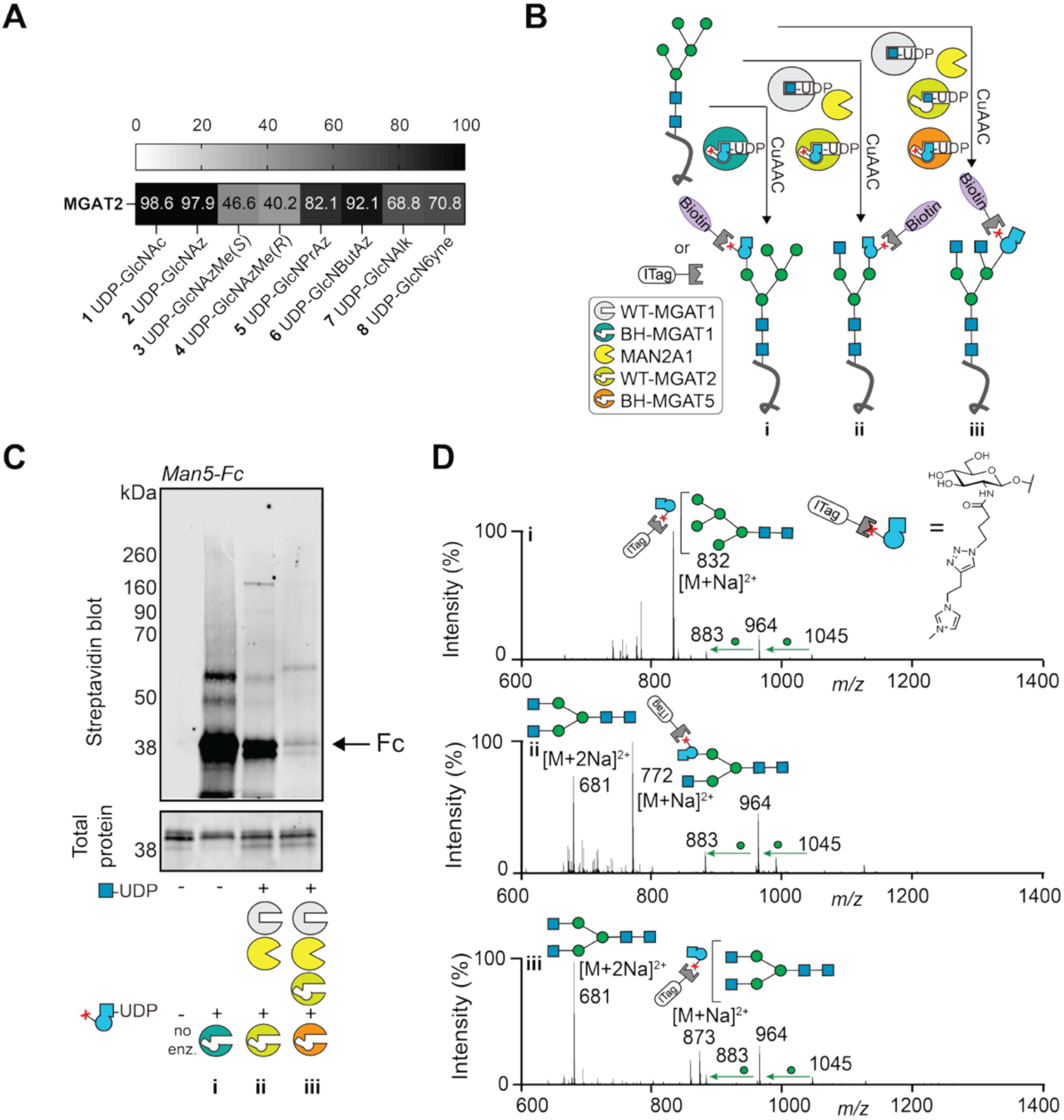
Differential bioorthogonal tagging of N-glycan intermediates with UDP-GlcNButAz **6**. *A*, enzymatic turnover using recombinant MGAT2 and a procainamide (PA)-labelled A1 acceptor (16-h reaction) as assessed by UHPLC with detection by absorption (**1** and analogs **3-8**) or mass spectrometry (analog **2**) (see Supporting Fig. 7 for representative traces for incorporation of individual compounds). Data are means from two independent replicates. See Supporting Fig. 6 for individual data points. *B*, schematic of differential N-glycoprotein tagging with UDP-GlcNButAz **6** by BH-MGAT1 (i), WT-MGAT2 (ii) or BH-MGAT5 (iii), with subsequent treatment with biotin-alkyne or ITag-alkyne under CuAAC conditions. *C*, differential bioorthogonal tagging of Man5-Fc, as assessed by streptavidin blot after CuAAC with biotin-alkyne. Reactions contained 1 mM UDP-GlcNAc **1** and 0.2 mM UDP-GlcNButAz **6** where indicated. Data are from one experiment. *D*, Released N-glycan ion mobility (IM)-MS spectra of Man5-Fc after differential N-glycoprotein tagging with UDP-GlcNButAz **6** and BH-MGAT1 (i), WT-MGAT2 (ii) or BH-MGAT5 (iii) followed by click reaction with ITag-alkyne. Data shown are IM extracted doubly charged ions and are from one experiment.

### Evaluation of the BH-MGAT1/UDP-GlcNButAz **6** enzyme-substrate pair

We next compared the ability of both BH- and WT-MGAT1 to incorporate UDP-GlcNButAz **6** or the native substrate UDP-GlcNAc (Fig. 2A). A Michaelis-Menten kinetic experiment revealed that the K_M_ of WT-MGAT1 toward UDP-GlcNButAz **6** is more than 5-fold higher than of BH-MGAT1, while the k_cat_ is approx. 1.2-fold higher for the BH-MGAT1/UDP-GlcButAz **6** enzyme-substrate pair (Fig. 2B). The K_M_ of BH-MGAT1/UDP-GlcNButAz **6** (0.11 mM) is approx. two-fold lower than the K_M_ of the native enzyme-substrate pair WT-MGAT1/UDP-GlcNAc (0.20 mM).^10,43^ The catalytic efficiency k_cat_/K_M_ of BH-MGAT1/UDP-GlcNButAz **6** (409 M^-1^s^-1^) is approx. 7.3-fold lower than of the native enzyme-substrate pair (3000 M^-1^s^-1^).^10,43,44^ To further assess the acceptance of both UDP-sugars, we performed a competition experiment in which recombinant BH- and WT-MGAT1were treated with both UDP-GlcNAc **1** and UDP-GlcNButAz **6** in the presence of Man5-PA. Turnover with each UDP-sugar was assessed individually through the difference in retention time between both products by UPLC (Fig. 2C). While BH-MGAT1 did not accept UDP-GlcNAc even if used in 4-fold excess over UDP-GlcNButAz **6**, WT-MGAT1 preferred UDP-GlcNAc in all assays, with up to 55-fold higher incorporation of GlcNAc than GlcNButAz into substrate Man5-PA.

### Bioorthogonal tagging of MGAT1 substrate glycoproteins

We turned to application of BH-MGAT1/UDP-GlcNButAz **6** to chemical glycoprotein tagging *in vitro*. As a model glycoprotein substrate, we used an antibody Fc dimer expressed in the superMan5 *Pichia pastoris* yeast strain to contain a single Man5 N-glycan per polypeptide.^45,46^ We subjected this Man5-Fc construct to *in vitro* glycosylation using WT- or BH-MGAT1 and both UDP-GlcNAc **1** and UDP-GlcNButAz **6** in different ratios. Glycoprotein preparations were treated with alkyne-biotin under copper-catalyzed azide–alkyne cycloaddition (CuAAC) conditions and azidosugar modification was detected by streptavidin blot. Consistent and intense streptavidin signal was observed when BH-MGAT1 was used together with UDP-GlcNButAz **6** (Fig. 2D; Supporting Fig. 4). The signal was not diminished by the presence of UDP-GlcNAc, either in equimolar quantity or in 4-fold excess. In contrast, WT-MGAT1 led to minor streptavidin signal that was outcompeted by a 4-fold excess of UDP-GlcNAc. As a more complex source of substrate glycoproteins, we next chose a membrane protein fraction of the Lec1 CHO cell line that lacks MGAT1 and displays Man5 as the major N-glycan type.^17,18^ By employing CuAAC and streptavidin blot, we found that the BH-MGAT1/UDP-GlcNButAz **6** enzyme-substrate combination leads to strong signal irrespective of the concentration of UDP-GlcNAc used (Fig. 2D). WT-MGAT1 leads to a slight increase of streptavidin signal over background that is outcompeted by an excess of UDP-GlcNAc. These data suggest that BH-MGAT1/UDP-GlcNButAz **6** is a bump-and-hole enzyme-substrate pair to trace the activity of MGAT1 *in vitro*, with selectivity conferred over WT-MGAT1 primarily as an effect of a favourable K_M_.

**Fig. 4:**
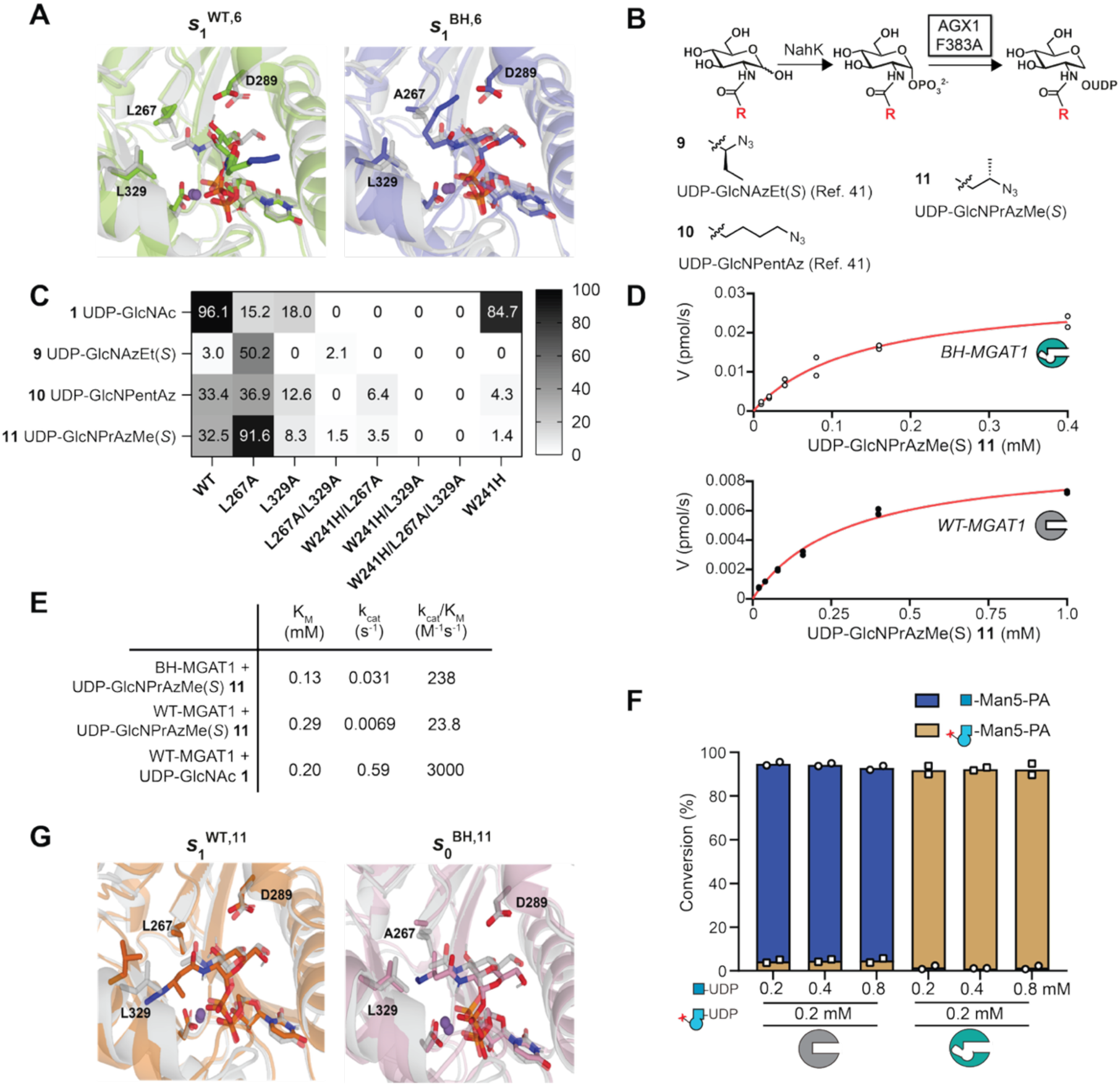
Structural refinement of MGAT1 BH engineering. *A*, VAMPnets binding poses comparing the binding of UDP-GlcNButAz **6** to WT- and BH-MGAT1 (PDB 2APC)^43^. The two major states, 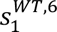 and 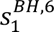, are shown. The binding pose of UDP-GlcNAc in WT-MGAT1 in the crystal structure, 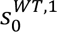, is shown in white. *B,* Collection of three additional bumped, bioorthogonal UDP-GlcNAc analogs including newly synthesized compound **11**. *C,* enzymatic turnover using recombinant MGAT1 and Man5-PA as acceptor (16-h reaction) as assessed by UHPLC with detection by absorption at 302 nm (**1** and analogs **9-11**). Data are means from two independent replicates. See Supporting Fig. 13 for individual data points. *D,* Michaelis-Menten kinetic experiment of either BH-MGAT1 or WT-MGAT1 with UDP-GlcNPrAzMe(*S*) **11**. Data are individual values of two independent experiments. Data were fitted with a Michaelis-Menten model (red line) *E,* calculated kinetic parameters for enzyme-substrate pairs measured through curve fitting. *F*, competition experiment between use of UDP-GlcNAc and UDP-GlcNPrAzMe(*S*) **11**. Both UDP-sugars were supplied in 16-h *in vitro* glycosylation experiments in depicted concentrations and individual reaction products were quantified. Data are individual datapoints and means from two independent replicates. *G*, VAMPnets binding states comparing the binding of UDP-GlcNPrAzMe(*S*) **11** to WT- and BH-MGAT1 based on (RSCB PDB: 2APC).^42^ The two major states, 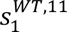 and 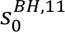, are shown.

Our previous BH campaign for the GlcNAc transferase MGAT5 employed UDP-GlcNButAz **6** for selective tagging of N-glycoproteins *in vitro*. We assessed whether the same compound **6** could be used for differential chemoenzymatic tagging of N-glycan intermediates, employing suitable GlcNAc transferases. Data by Kohler and colleagues suggested that the transferase MGAT2 is highly promiscuous towards chemical acylamide modifications in UDP-GlcNAc.^33,47^ We therefore expressed recombinant His-tagged MGAT2 in Expi293F cells (Supporting Fig. 5) and tested whether WT-MGAT2 incorporates UDP-GlcNAc **1** and analogs **2**-**8** into a procainamide-labelled acceptor glycan termed A1-PA *in vitro*. MGAT2 exhibited profound substrate promiscuity, incorporating all tagged GlcNAc analogs to more than 40% in 16-h reactions (Fig. 3A; Supporting Fig. 6). UDP-GlcNButAz **6** led to almost quantitative incorporation by WT-MGAT2. This finding suggested that differential tagging with GlcNButAz was within reach *in vitro*, employing chemoenzymatic assembly of an N-glycan core by WT-GlcNAc transferases and either BH-MGAT1, WT-MGAT2 or MGAT5^F458V/F517L^ (termed BH-MGAT5) for GlcNButAz incorporation (Fig. 3B). Incubation of Man5-Fc with BH-MGAT1/UDP-GlcNButAz **6** led to efficient chemical tagging that was detectable by streptavidin blot after treatment with biotin-alkyne under CuAAC conditions (Fig. 3B, C, sample i).

**Fig. 5:**
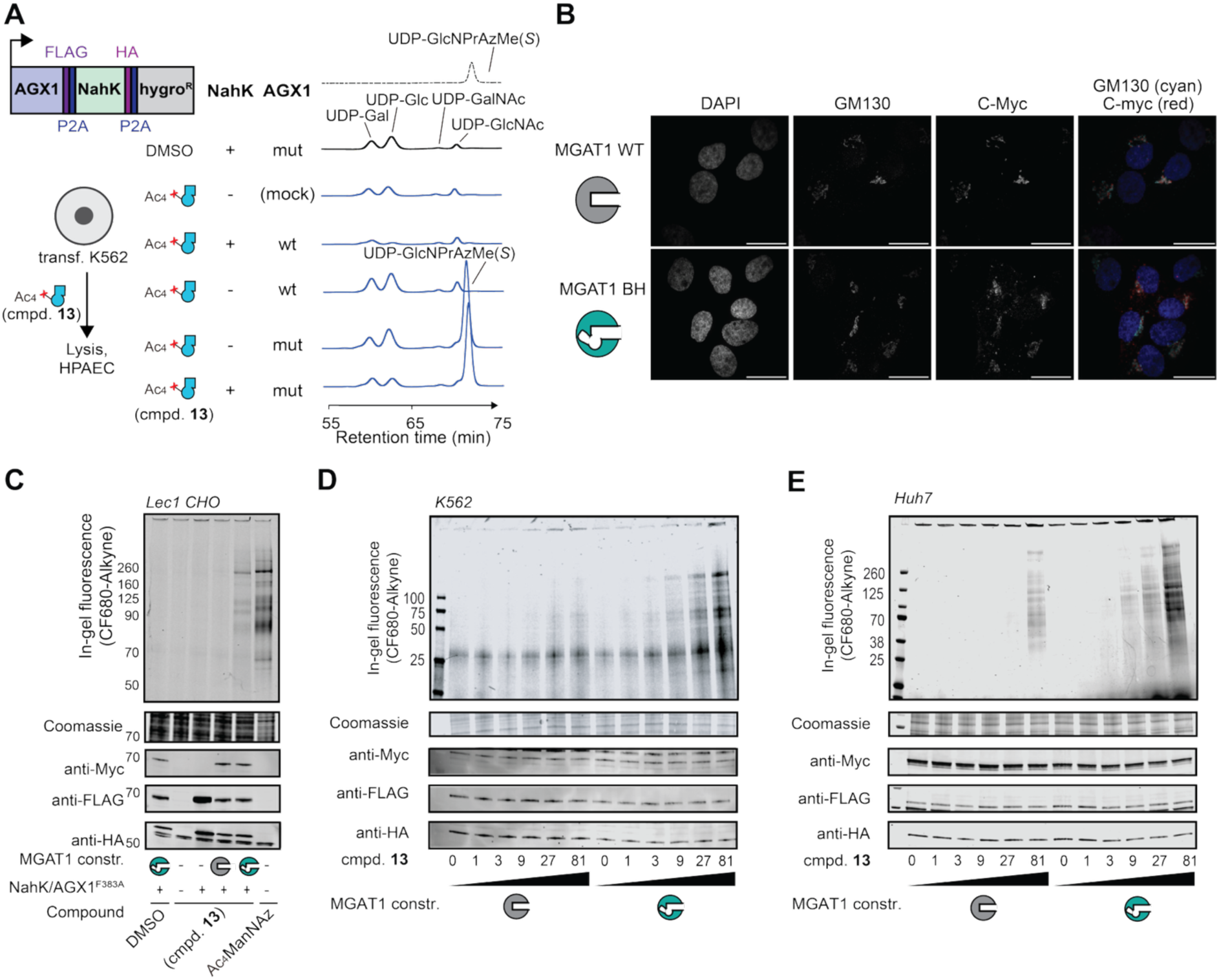
In-cell application of the GlcNPrAzMe(*S*)/BH-MGAT1 bump-and-hole system. *A,* biosynthesis of UDP-GlcNPrAzMe(*S*) **11** was assessed by anion exchange chromatography. Cells were fed with 250 μM Ac_4_GlcNPrAzMe (*S*) **13** or DMSO and extracts assessed by anion exchange chromatography. Data are from one representative out of two replicates. *B*, Fluorescence microscopy of Huh7 cells stably transfected with MGAT1 constructs, and subsequently stained. Scale bar, 20 nm. Data are from one experiment. *C*, incorporation of GlcNPrAzMe(*S*) into the cell-surface glycoproteome of Lec1 CHO cells. Cells were stably transfected with plasmids as indicated, fed with 9 μM Ac_4_GlcNPrAzMe(*S*) or 2.5 μM Ac_4_ManNAz, subjected to on-cell CuAAC with CF680-alkyne, and fluorescent tagging was assessed by in-gel fluorescence. Data are from one representative out of two independent replicates. *D*, *E,* incorporation of GlcNPrAzMe(*S*) into the cell-surface glycoproteome of K562 (*D*) cells and Huh7 (*E*) cells. Cells were fed with 0 or 1-81 µM Ac_4_GlcNPrAzMe(*S*) **13** and treated as in *C*. Data are from one representative out of 2 independent replicates for both *D* and *E*. See Supporting Figs. 23 and 24 for the other replicates.

When WT-MGAT1/UDP-GlcNAc **1** and MAN2A1 were employed to generate the A1 intermediate, WT-MGAT2/UDP-GlcNButAz **6** chemically tagged these intermediates (Fig. 3B, C, sample ii). Treating the A1 intermediate with WT-MGAT2 in the presence of only UDP-GlcNAc **1** yielded the so-called A2 precursor as a substrate for MGAT5.^48^ Treatment with BH-MGAT5/UDP-GlcNButAz **6** led to incorporation of GlcNButAz that was detectable by streptavidin blot (Fig. 3B, C, sample iii). We noted that incorporation efficiency decreases between earlier (Fig. 3C, sample i) and later (Fig. 3C, sample iii) stages of N-glycan maturation, potentially reflecting differences in enzymatic activities between ensuing GlcNAc transferases. We then traced differential bioorthogonal N-glycan tagging by mass spectrometry (Fig 3D, Supporting Fig. 8, 9). The approach integrates chemoenzymatic installation of GlcNButAz at defined N-glycan bifurcation sites of Man5-Fc with a clickable, positively charged imidazolium tag (ITag), enabling collision-induced dissociation tandem MS detection via a diagnostic 407.2 *m/z* ITag–GlcNButAz ion and glycan fragmentation patterns that distinguish tagged from native GlcNAc residues (Supporting Fig. 8, 9).^40,49–51^ Treatment with BH-MGAT1 and UDP-GlcNButAz **6** followed by CuAAC with ITag-alkyne allowed detection of the newly formed GlcNButAz-Man5 structure (832 *m/z*, Fig. 3D, top). When the A1 intermediate was generated by WT-MGAT1/UDP-GlcNAc **1**, it was chemically tagged by WT-MGAT2/UDP-GlcNButAz **6**. After CuAAC with ITag-alkyne, this allowed detection of the GlcNButAz-A1 structure (772 *m/z,* Fig. 3D, middle). The A2 intermediate was generated by treatment of the A1 intermediate with WT-MGAT2/UDP-GlcNAc **1**. The A2 intermediate could be chemically tagged by BH-MGAT5/UDP-GlcNButAz **6**.

Following CuAAC with ITag-alkyne, this allowed detection of the GlcNButAz-A2 structure (873 *m/z*, Fig. 3D, bottom). Additional low-abundance peaks at 883, 964 and 1045, corresponding to higher high-mannose glycans (Man_8_–Man_10_) were detected and are attributable to the intrinsic glycan heterogeneity of the yeast-produced Man5-Fc, which is known to exhibit a shift toward larger high-mannose structures under methanol-induced expression conditions, as previously reported.^52^

### Structural refinement of BH engineering leads to a cellular probe for MGAT1 activity

Our data indicated that UDP-GlcNButAz **6** can be used by WT GlcNAc transferases such as WT-MGAT1 (Fig. 1E) and WT-MGAT2 (Fig. 3A), thereby impacting selectivity towards BH-MGAT1 when employed in cellular models. To test this notion, we devised a strategy to deliver **6** to the cytosol of mammalian cells, employing per-acetylated GlcNButAz, Ac_4_GlcNButAz **12**, as a precursor. We tested an engineered metabolic pathway using *B. longum* NahK and the engineered human enzyme AGX1^F383A^ to assess UDP-GlcNButAz **6** biosynthesis. When human K562 cells expressing either AGX1^F383A^ alone or in combination with NahK were fed Ac_4_GlcNButAz **12**, UDP-GlcNButAz **6** formation was detectable in a nucleotide-sugar extract by anion exchange chromatography (Supporting Fig. 10). With a method for biosynthesis of UDP-GlcNButAz **6**, we assessed incorporation by cellular BH-MGAT1. Lec1 CHO cells expressing WT- or BH-MGAT1 as well as NahK/AGX1^F383A^ were fed with Ac_4_GlcNButAz **12**. Subsequent treatment with the clickable fluorophore CF680-alkyne allowed visualization of cell surface glycoproteins by in-gel fluorescence.^31,32,38,53^ Visible and comparable incorporation into cell-surface glycoproteins was seen in cells expressing either WT- or BH-MGAT1 constructs with Ac_4_GlcNButAz **12** concentrations above 3 μM. These data indicated that UDP-GlcNButAz **6** does not allow for selective incorporation by BH-MGAT1 in living cells (Supporting Fig. 11). Molecular dynamics (MD) simulations were performed to explore the structural basis for this lack of selectivity. We examined the binding of UDP-GlcNButAz **6** to WT- and BH-MGAT1 using a neural network approach (VAMPnets) on data generated by MD simulations. A two-state Markov model was used to estimate the relative populations of the binding states. Here, 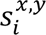 denotes the *i*th state of *y* in *x*, and 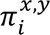 denotes the corresponding state occupancy. In the WT enzyme, UDP-GlcNButAz **6** adopts two nearly equal binding states (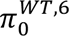 = 0.44, 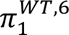 = 0.56; Fig. 4A *left*). In the minor state 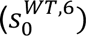, the azide tail extends between Leu267 and Leu329, similar to the orientation of UDP-GlcNAc in the published crystal structure (Supporting Fig. 12). In the major state 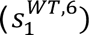, the pyranose ring rotates outward around its attachment point, reducing anomeric carbon accessibility and redirecting the C3/C4 hydroxyl groups toward Asp289. In the BH enzyme, the same two states appear, but with a much larger population (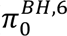 = 0.12, 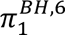 = 0.88; Fig. 4A *middle left*). The minor state 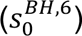 has the pyranose rotated outward and C3/C4 directed toward Asp289. The major state 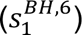 places the tail into the cavity created by the Leu267Ala mutation and largely maintains the pyranose orientation similar to 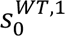 (Fig. 4A). Although the WT enzyme slightly favours 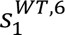 that is likely not catalytically active, the two states interconvert far more rapidly than catalysis itself (in ns). As a result, the WT enzyme effectively equilibrates immediately on experimental timescales and, under forward reaction conditions, will continue to turnover until it reaches (near) complete conversion.

With the aim of identifying a more selective MGAT1 bump-and-hole enzyme-substrate pair, we explored a series of UDP-GlcNAc analogs **9**, **10** and newly-synthesized **11** with more sterically encumbered acylamides (Fig. 4B). In comparison to the 4-azidobutyramide in compound **6**, UDP-GlcNPentAz **10** exhibited a longer linear 5-azidopentamide chain, whereas UDP-GlcNAzEt(*S*) **9** and UDP-GlcNPrAzMe(*S*) **11** are branched isomers of **6**. Again, *in vitro* enzymatic turnover was performed to assess the ability of MGAT1 mutants to accept the modified UDP-GlcNAc analogs, using Man5-PA as the acceptor substrate (Fig. 4C; Supporting Fig. 13). Out of the MGAT1 variants tested, BH-MGAT1 variant Leu267Ala showed the highest enzymatic activity. The Leu267Ala variant accepted UDP-GlcNPrAzMe(*S*) **11** as the best substrate with 91% conversion, while acceptance of the endogenous UDP-GlcNAc **1** was 15% in a 16 hour reaction. WT-MGAT1 converted UDP-GlcNPrAzMe(*S*) **11** with 2.8-fold lower turnover compared to the Leu267Ala variant, indicating increased selectivity of this BH pair. In 30 min reactions, the BH-MGAT1 Leu267Ala variant accepted **11** in 60% conversoin, compared to 78% with previously identified UDP-GlcNButAz **6**. In contrast, WT-MGAT1 accepted **11** in 9% conversion, compared to 56% for UDP-GlcNButAz **6** (Supporting Fig. 14). These data suggest that UDP-GlcNPrAzMe(*S*) **11** exhibits superior selectivity for BH-MGAT1 compared to UDP-GlcNButAz **6** *in vitro*.

We then compared the ability of both BH- and WT-MGAT1 to incorporate UDP-GlcNPrAzMe(*S*) **11** (Fig. 4D). Michaelis-Menten kinetics revealed that the K_M_ of WT-MGAT1 toward UDP-GlcNPrAzMe(*S*) **11** is approx. 2.2-fold higher than that of BH-MGAT1, while the k_cat_ is approx. 4.5-fold higher for the BH-MGAT1/UDP-GlcNPrAzMe(*S*) **11** enzyme-substrate pair (Fig. 4E). As a result, the catalytic efficiency k_cat_/K_M_ of BH-MGAT1/UDP-GlcNPrAzMe(*S*) **11** is approx. 10-fold higher than that of WT-MGAT1/UDP-GlcNPrAzMe(*S*) **11**. The acceptance of **11** was further assessed in a competition experiment (Fig. 4F). BH-MGAT1 did not accept UDP-GlcNAc **1** even when used in a 4-fold excess over UDP-GlcNPrAzMe(*S*) **11**. WT-MGAT1 accepted UDP-GlcNPrAzMe(*S*) **11** to a negligible extent and showed strong preference (>95%) for the endogenous substrate UDP-GlcNAc **1** in all assays.

Although both compounds **6** and **11** display enhanced activity toward BH-MGAT1, their selectivity arises from different structural origins. To understand what features of the BH-MGAT1/UDP-GlcNPrAzMe(*S*) **11** pair drive its kinetic preference, we turned again to MD simulations of either **6** or **11** docked into the active site of WT- or BH-MGAT1. For WT-MGAT1, 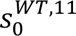 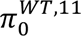 = 0.07) is a minor state in which the pyranose ring retains the orientation seen in the WT-MGAT1/UDP-GlcNAc crystal structure (Supporting Fig. 15), while 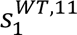 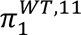 = 0.93) displaces the sugar ring further from Asp289 such as seen with compound **6** in 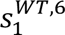 (Fig. 4G). For the BH enzyme, the major state, 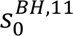 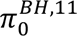 = 0.57), adopts a likely catalytically active pyranose orientation (Fig. 4G), while the minor state 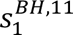 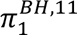 = 0.43) features the pyranose rotating toward Leu329 (Supporting Fig, 15). Computational kinetic analysis revealed that WT-MGAT1 enters the likely inactive state 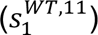 much faster than it escapes from it, leading to a non-productive enzyme-substrate pair. By contrast, in BH-MGAT1, the equilibrium state distribution is shifted toward 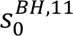 that resembles the structure of the native enzyme-substrate pair. As a result, the selectivity manifests as a higher conversion rate in BH-MGAT1. WT-MGAT1 can still reach complete conversion under forward-driven conditions but does so significantly more slowly. Catalytic efficiencies show a ∼10-fold advantage for BH-MGAT1 toward UDP-GlcNPrAzMe(*S*) **11** (Fig. 4E), consistent with the ∼8.3-fold difference in productive-state occupancy (BH ≈ 0.57 vs WT ≈ 0.07). Overall, compound **11** is thermodynamically and kinetically favoured to occupy a productive orientiation in BH-MGAT1 over WT-MGAT1.

We also assessed whether UDP-GlcNPrAzMe(*S*) **11** is an effective substrate for WT-MGAT2 in the presence of UDP-GlcNAc **1** (Supporting Fig. 16). While compound **11** is accepted by WT-MGAT2, turnover of UDP-GlcNButAz **6** was 1.5- to 2-fold higher than that of **11** at equal concentration and reaction times. Thus, BH-MGAT1/UDP-GlcNPrAzMe(*S*) **11** is a more specific reporter pair for the activity of MGAT1 compared to BH-MGAT1/UDP-GlcNButAz **6**, due to its reduced acceptance by WT-MGAT1 and WT-MGAT2.

Finally, we attempted to take inspiration from Mukerjee et al. who introduced an OTs group at 6-position of a GlcNAc analog for acceptance by WT-MGAT1.^41^ To assess whether 6-OTs could be integrated into our scaffold, we furnished the corresponding analogs of GlcNButAz **14**, GlcNPrAzMe(*S*) **15** and GlcNAlk **16** (Supporting Fig. 17). Compound **16** was the design used by Mukherjee et al. and made as a control.^41^ We first assessed whether the tosylated GlcNAc analogs are substrates for the biosynthetic enzymes NahK, WT-AGX1 and AGX1^F383A^ through *in vitro* enzymatic assays. None of the GlcNAc analogs **14**, **15** or **16** were converted into their corresponding UDP-GlcNAc analogs under these conditions (Supporting Fig. 18), indicating that the 6-toluenesulfonyl group may be associated with a metabolic roadblock towards UDP-GlcNAc synthesis by NahK/AGX1^F383A^. Since NahK is a bacterial enzyme, we note that human biosynthetic pathways may still be able to convert tosylated GlcNAc analogs to their corresponding UDP-sugars. However, we also note that derivatizing the 6-position of GlcNAc analogs with a OTs group would block the first biosynthetic activation step to phosphorylate the 6-position.

### Selective incorporation of GlcNPrAzMe by BH-MGAT1 into cell-surface glycoproteins

To establish a cellular BH-MGAT1 system, we tested biosynthesis of UDP-GlcNPrAzMe*(S*) **11** by NahK/AGX1^F383A^ (Fig. 5A and Supporting Fig. 19). K562 cells fed with Ac_4_GlcNPrAzMe(*S*) **13** biosynthesized the UDP-sugar **11** in the presence of AGX1^F383A^ irrespective of NahK expression, as assessed by anion exchange chromatography in comparison with synthetic **11** as a standard. Cells overexpressing WT-AGX1 or transfected with an empty plasmid did not exhibit biosynthesis of **11**. We next confirmed that both WT- and BH-MGAT1 localize to the Golgi compartment. Immunofluorescence microscopy of cells overexpressing either MGAT1 construct with a myc epitope tag showed colocalization with the Golgi marker giantin, confirming native localization of the engineered MGAT1 (Fig. 5B).

With all components of a cellular bump-and-hole system confirmed individually, we assessed incorporation of GlcNPrAzMe(*S*) by BH-MGAT1 into cell-surface glycans. Lec1 CHO cells expressing WT- or BH-MGAT1 as well as NahK/AGX1^F383A^ were fed with Ac_4_GlcNPrAzMe(*S*) **13** (Fig. 5C; Supporting Fig. 20). Subsequent treatment with CF680-alkyne under CuAAC conditions allowed visualization of cell surface glycoproteins by in-gel fluorescence. In *Mgat1*-deficient Lec1 CHO cells, no fluorescent signal was observed in cells fed with DMSO vehicle control. Cells fed with Ac_4_GlcNPrAzMe(*S*) **13** exhibited fluorescent signal only when containing the NahK/AGX1^F383A^ biosynthetic pathway and BH-MGAT1. No fluorescent signal was observed in cells expressing WT-MGAT1, only NahK/AGX1^F383A^ or transfected with empty plasmid. A specific band pattern of fluorescently tagged glycoproteins was observed in cells expressing the MGAT1 BH system. Ac_4_ManNAz, a well-characterized azide-containing sialic acid precursor, was used as a control and led to a larger number of fluorescently tagged cell surface glycoprotein bands. These data confirm that BH-MGAT1 specifically tags a subset of cell surface glycoproteins.

Chemical tagging of cell-surface glycans by the BH-MGAT1 system was further confirmed in human lymphoblast K562 (Fig. 5D, Supporting Fig. 21) and hepatocarcinoma Huh7 cells (Fig. 5E, Supporting Fig. 22). In both cell lines, expressing the BH-MGAT1 system together with NahK/AGX1^F383A^ led to increased fluorescent intensity at lower concentrations of Ac_4_GlcNPrAzMe(*S*) **13** than in cells expressing WT-MGAT1 with NahK/AGX1^F383A^ (Fig. 5D, E, and Supporting Figs. 23 and 24). BH-MGAT1 led a dose-dependent increase in signal intensity over a concentration range from 1 μM to 81 μM. Together, these data suggest that GlcNPrAzMe(*S*) is specifically incorporated into certain cell-surface glycoproteins by BH-MGAT1 in a range of cell lines.

We also investigated incorporation of tosylated GlcNAc analogs into cell surface glycans of K562 cells. Compounds **17**, **18**, and **19** were made as peracetylated versions of GlcNAc analogs **14**, **15**, and **16**, respectively. None of the tosylated compounds achieved a fluorescent tagging intensity comparable to the BH-MGAT1/UDP-GlcNPrAzMe(*S*) **11** pair in cells (Supporting Figs. 25 and 26), with fluorescence tagging visible at concentrations of 81 μM. These results demonstrate that rational bump-and-hole engineering led to a reporter system for MGAT1 activity using Ac_4_GlcNPrAzMe(*S*) **13**, with intense, consistent and selective chemical tagging of cell surface glycoproteins.

## Conclusion

Tools to probe the glycoproteome are essential to shed light on the intricacies of glycans in physiology. Deficiencies in N-glycan biosynthesis are associated with a number of congenital disorders,^1,2^ highlighting the importance of these modifications in development. Furthermore, glycosylation plays a key role in cancer, and defined N-glycan substructures are essential for functional maturation of the SARS-CoV2 spike protein.^54^ As part of our efforts to probe the glycoproteome with chemical tools, investigating MGAT1 was an important target as a bifurcation point between oligo-mannose and hybrid or complex glycans. We note that the pentamannosyl glycan substrate of MGAT1 needs to be provided by the activity of a set of mannosidases that have been proposed as the main differentiator between high-mannose and other N-glycans, based on steric factors.^55,56^ Since most pentamannosyl N-glycans will be processed by MGAT1, a bioorthogonal reporter for MGAT1 activity should provide the basis for an accurate view on glycan accessibility to mannosidases, and for investigation of the switch between oligo-mannose and all other N-glycans.

Our MGAT1 bump-and-hole system benefited from an iterative optimization approach to achieve selective chemical tagging of MGAT1 substrates in living cells. Although our initial screen of seven UDP-GlcNAc analogs identified UDP-GlcNButAz **6** as the most selective “bumped” analog, cellular studies suggested no selectivity for BH-MGAT1 over background incorporation. Inspired by experimental and MD studies, we iterated the BH system towards UDP-GlcNPrAzMe(*S*) **11** as a superior “bumped” analog with selective incorporation by BH-MGAT1 in cells. Further optimization efforts through tosyl modification at the 6-position were unsuccessful in our hands. Mukherjee, Hanover and colleagues detected the tosylated UDP-GlcNAc analog in cells and on glycoproteins after treatment of 100 μM precursor concentrations.^42^ Thus, incorporation of such analogs is subject to further experimental investigation.

Taken together, our studies strongly support the combination of synthetic nucleotide-sugars, *in silico* calculations and enzymology for successful BH engineering, bolstering such approaches in the future.

## Acknowledgements

We thank Prof. Pamela Stanley (Albert Einstein College of Medicine, New York, USA) for kindly providing Lec1 CHO cell lines. We thank Dr. Mia Zol-Hanlon (Francis Crick Institute) for help growing Expi293F cells. We thank Dr. Zach Armstrong and Prof. Gideon J. Davies for providing recombinant MAN2A1. We thank Dr. Sharon Tooze for kindly providing anti-GM130 antibody. We thank the Francis Crick Institute Cell Sciences, Structural Biology and Chemical Biology Science Technology Platforms for valuable help. This work was supported by the Francis Crick Institute (to B. S.) which receives its core funding from Cancer Research UK (CC2127), the UK Medical Research Council (CC2127) and Wellcome Trust (CC2127). This work was supported by UK Research and Innovation (UKRI) under the UK government’s Horizon Europe funding guarantee (grant numbers EP/X042383/1 and EP/Y032527/1 to B. S.) underwriting association to a H2020 Marie Skłodowska-Curie Actions Doctoral Network (project GLYCO-N, grant 101119499), and a UKRI Leaders Fellowship (MR/V02213X/1 and UKRI2014) to W. B. S. We thank the BBSRC (BB/V014862/11 and BB/V008439/1 to B. S.) This work was supported by Cancer Research UK (DRCMDP-Nov22/100011). S. A. B. is funded by Wellcome Trust Early-Career Award 312750/Z/24/Z. We gratefully acknowledge the Howard Hughes Medical Institute for funding C.R.B. (project number: CC31920). K.B. acknowledges a Children’s Tumor Foundation Young Investigator Award for postdoctoral funding. Plasmids used herein were generated by Moremen et al.^57^ for the PSI Materials Repository related to human and bacterial glycosylation enzymes (Glyco-enzyme clone collection) and generated through the support of NIH grant RR005351. We are grateful for the technical assistance provided by the support of the CTE-Power supercomputer of the Barcelona Supercomputing Center (BSC–CNS), within the Red Española de Supercomputación (RES). C.R.B. is a co-founder and scientific advisory board member of GanNA Bio, Neuravid, Firefly Bio, Lycia Therapeutics, Palleon Pharmaceuticals, Enable Bioscience, Redwood Biosciences (a subsidiary of Catalent), OliLux Bio, Grace Science and InterVenn Biosciences. C.R.B. is a member of the Board of Directors of Alnylam, Xaira Therapeutics, Acepodia and OmniAb.

