## Supporting Information for "Iterative Bump-and-hole engineering creates a bioorthogonal reporter for *N*-acetylglucosaminyltransferase I"

[d] Institució Catalana de Recerca i Estudis Avançats (ICREA), 08020 Barcelona, Spain.

[e] Department of Biochemistry, Dorothy Crowfoot Hodgkin Building, University of Oxford, Oxford OX1 3QU, United Kingdom.

[f] The Kavli Institute for Nanoscience Discovery, Dorothy Crowfoot Hodgkin Building, University of Oxford, Oxford OX1 3QU, United Kingdom.

[g] Department of Chemical Engineering, Imperial College London, London, UK

[h] Stanford Sarafan ChEM-H, Stanford University, Stanford, California 94305, USA.

[i] Department of Chemistry, Stanford University, Stanford, California 94305, USA.

[j] Howard Hughes Medical Institute, Stanford University, Stanford, California 94305, USA.

[k] Faculty of Chemistry and Food Chemistry, TUD Dresden University of Technology, 01067 Dresden, Germany.

[l] Current address: College of Chemistry and Molecular Engineering, Peking University, Beijing 100871, China.

[#] These authors contributed equally.

**Supporting Figures**

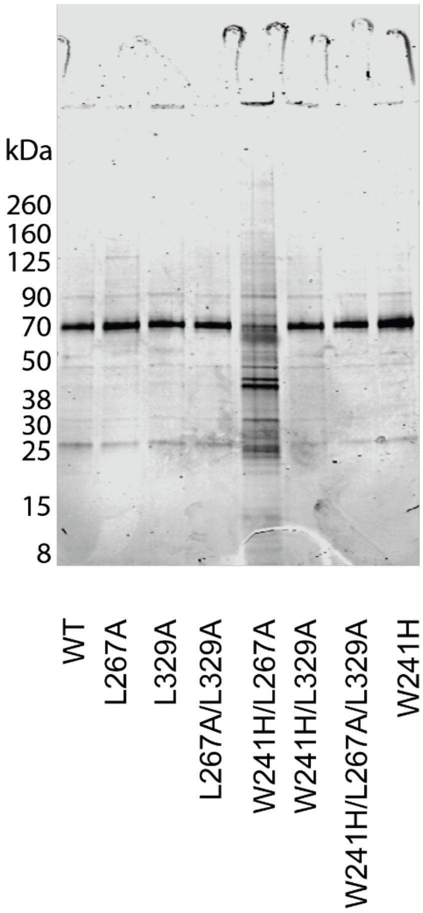


**Supporting Fig. 1:** Recombinant MGAT1 variants as assessed by SDS-PAGE and Coomassie staining.

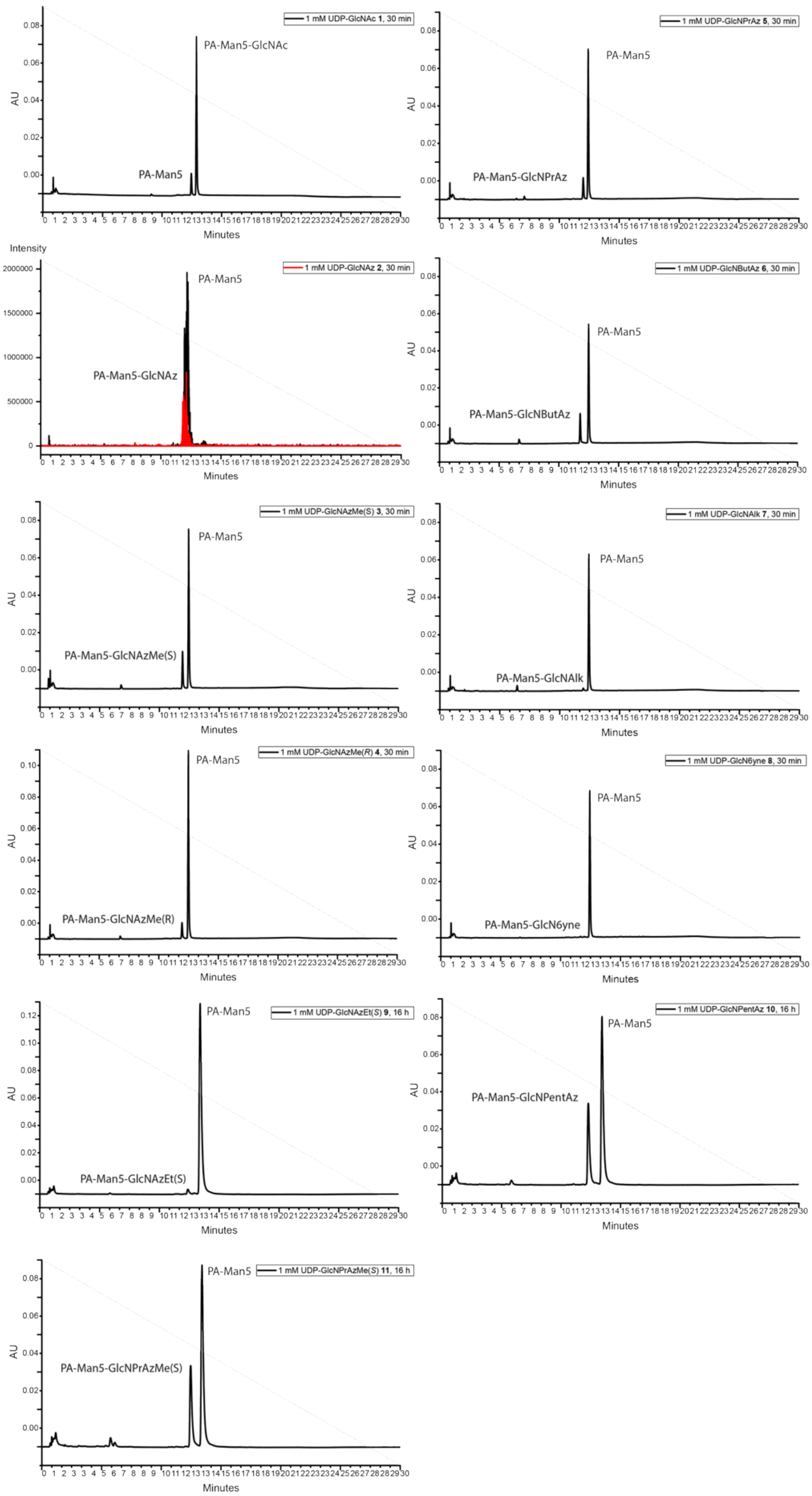


**Supporting Fig. 2:** Representative traces for incorporation of GlcNAc analogs into a procainamide (PA)-tagged Man5 sugar (Man5-PA). Reactions were carried out using 1 mM UDP-GlcNAc or UDP-GlcNAc analogs, 0.2 mM Man5-PA, and 250 nM BH-MGAT1. Turnover of UDP-GlcNAc **1** and analogs **3-11** into the corresponding product glycan was measured by the near-UV trace (302 nm absorption) of procainamide in the acceptor substrate and the product glycans. The conversion of analog **2** was measured by MS due to co-elution of the substrate and the product glycans.

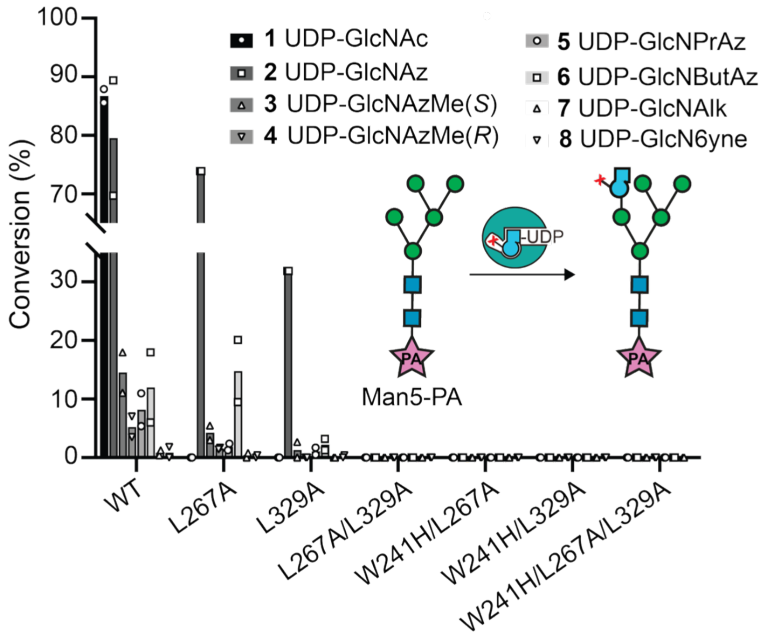


**Supporting Fig. 3:** MGAT1 variants were subjected to UDP-GlcNAc **1** and UDP-GlcNAc analogs **2-8** in endpoint enzymatic assays, using Man5-PA as an acceptor substrate and analyzed by absorption at 302 nm or MS on UHPLC. Data are individual data points and means from two independent replicates.

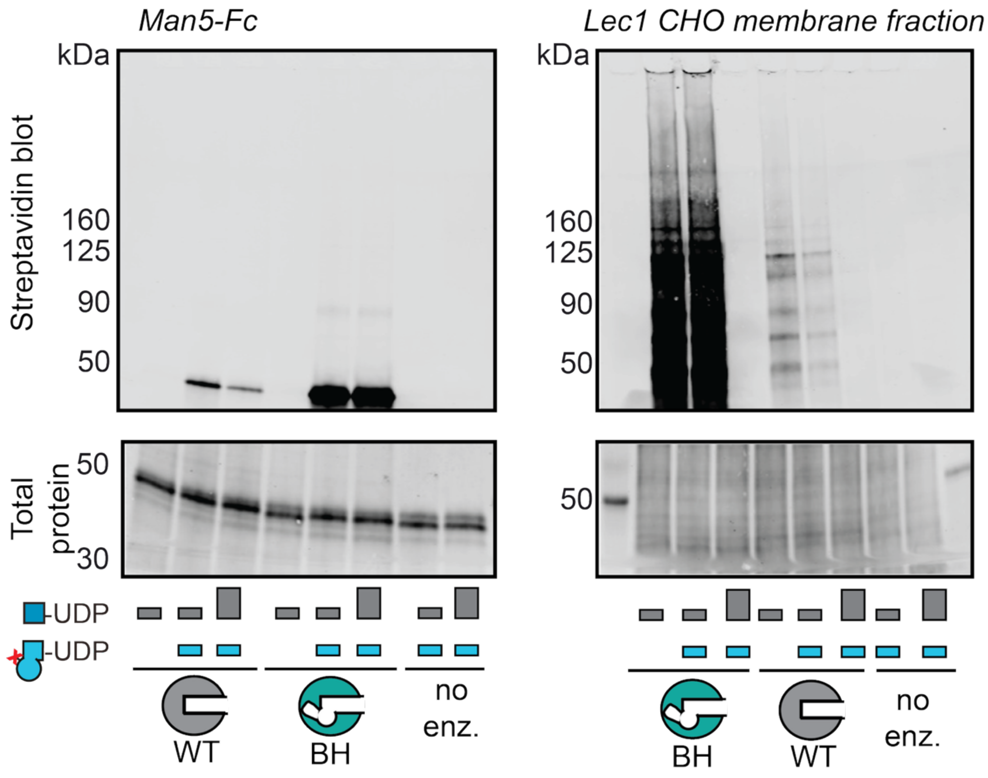


**Supporting Fig. 4:** Second independent biological replicate of *in vitro* glycosylation of Man5-containing glycoprotein substrates by BH-engineered MGAT1. Glycoproteins were treated with WT- or BH-MGAT1 and the substrates UDP-GlcNAc (grey bars) and UDP-GlcNButAz **6** (blue bars) in either 0.2 mM (small bars) or 0.8 mM (big bars) concentration. *In vitro* glycosylation was applied to Man5-Fc (left) or a Lec1 CHO cell membrane fraction (right). Reaction mixtures were treated with biotin-alkyne under CuAAC conditions and bioorthogonal tagging analyzed by streptavidin blot.

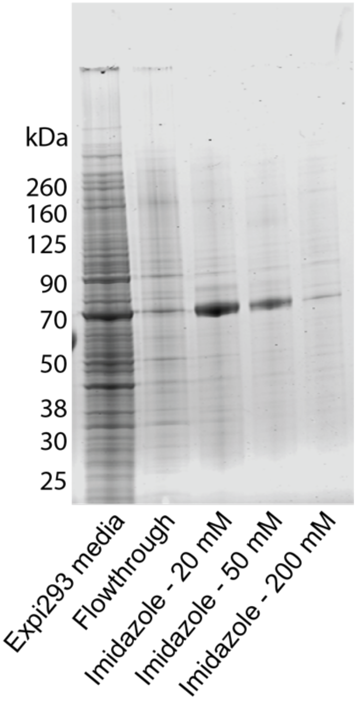


**Supporting Fig. 5:** Ni-NTA-purified recombinant MGAT2 variants were assessed by SDS-PAGE followed by Coomassie staining.

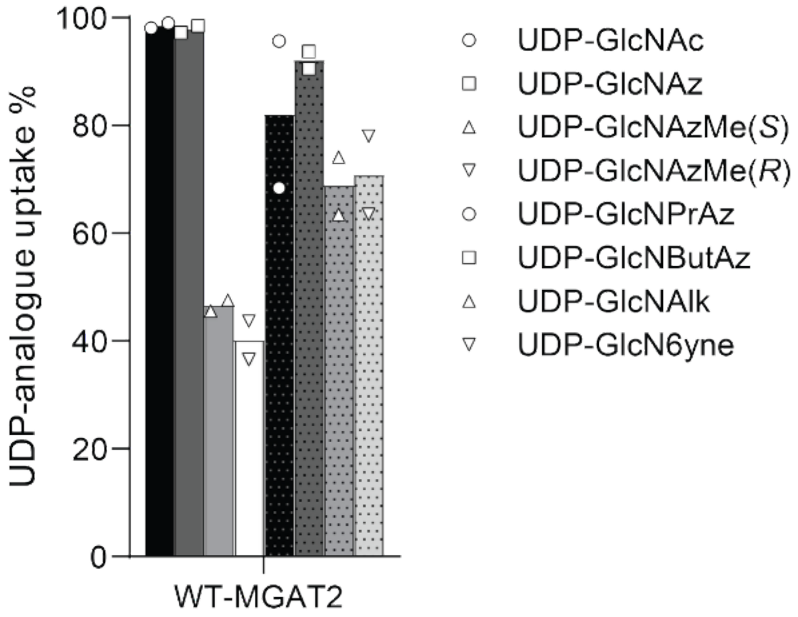


**Supporting Fig. 6:** Wildtype MGAT2 was subjected to UDP-GlcNAc **1** and UDP-GlcNAc analogs **2-8** in endpoint enzymatic assays, using a procainamide (PA)-tagged A1 sugar as an acceptor substrate and analyzed by absorption at 302 nm or MS on UPLC-MS. Data are individual data points and means from two independent replicates.

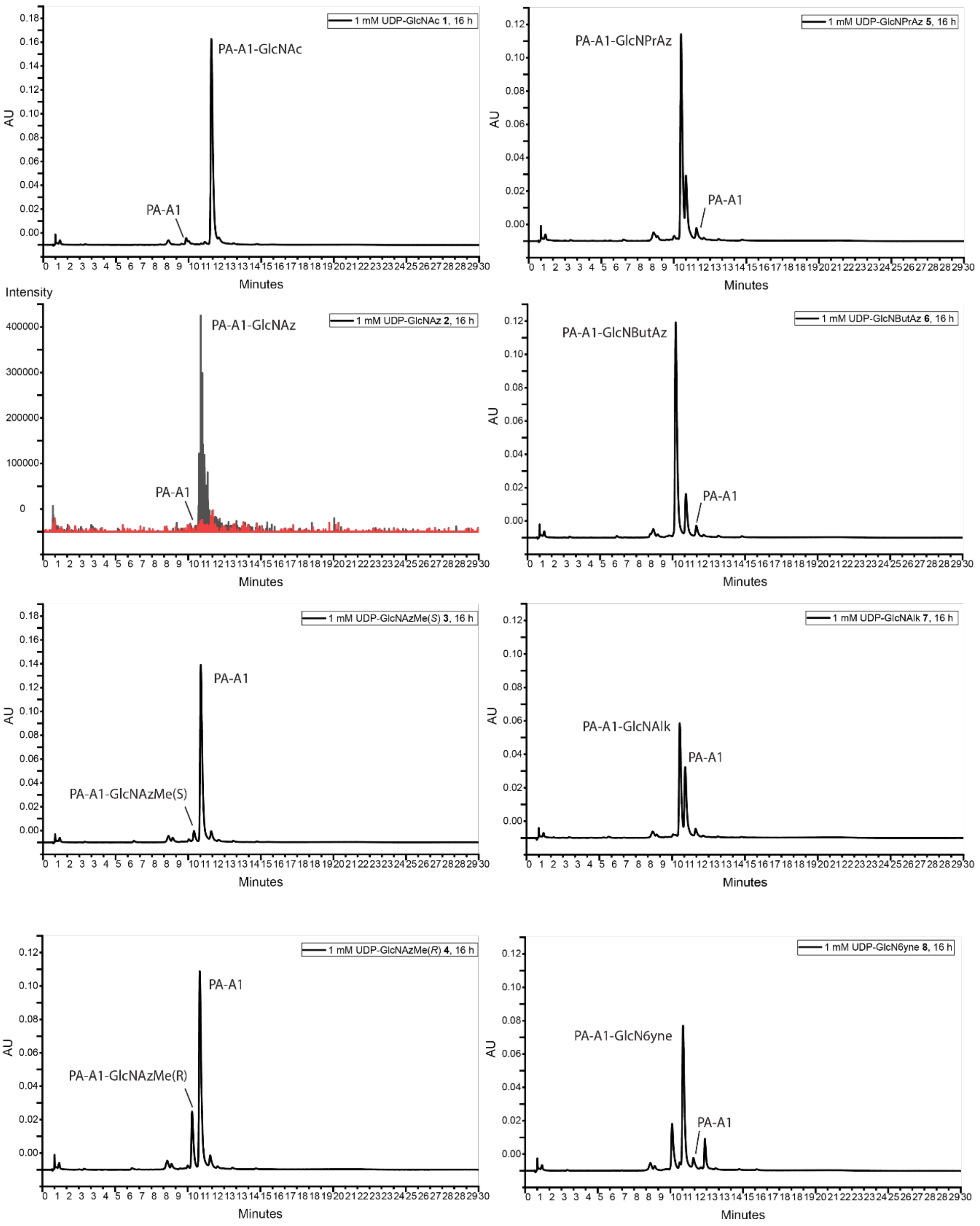


**Supporting Fig. 7:** Representative traces for incorporation of GlcNAc analogs into A1-PA. Reactions were carried out using 1 mM UDP-GlcNAc or UDP-GlcNAc analogs, 0.2 mM A1-PA, and 130 nM WT-MGAT2. Turnover of UDP-GlcNAc **1** and analogs **3-8** into the corresponding product glycan was measured by the near-UV trace (302 nm absorption) of procainamide in the acceptor substrate and the product glycans. Conversion of analog **2** was measured by MS due to co-elution of the substrate and the product glycans.

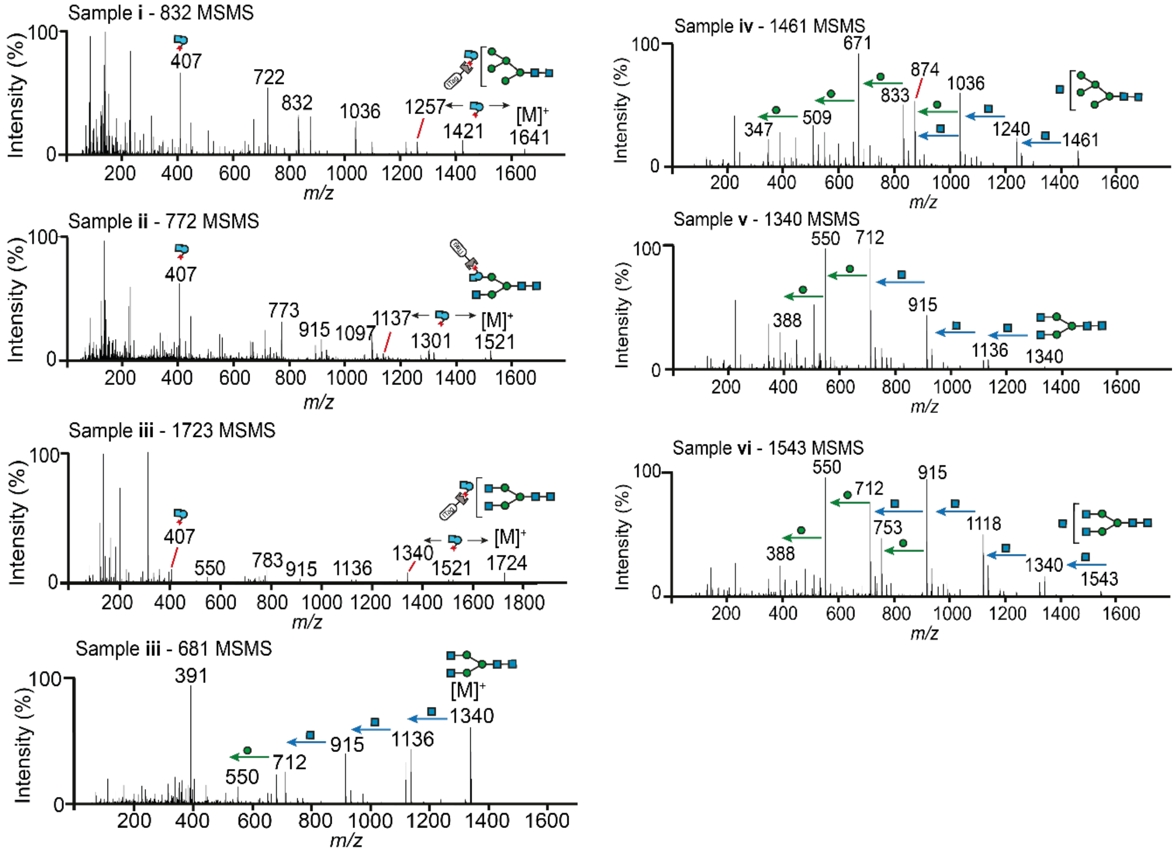


**Supporting Fig. 8**: Released N-glycan MS/MS spectra of Man5-Fc after differential N-glycoprotein tagging with either UDP-GlcNButAz **6** and BH-MGAT1 (i), WT-MGAT2 (ii) or BH-MGAT5 (iii) followed by click reaction with ITag-alkyne, or UDP-GlcNAc **1** and WT-MGAT1 (iv), WT-MGAT1 (v) or WT-MGAT5 (vi). Data are from one experiment.

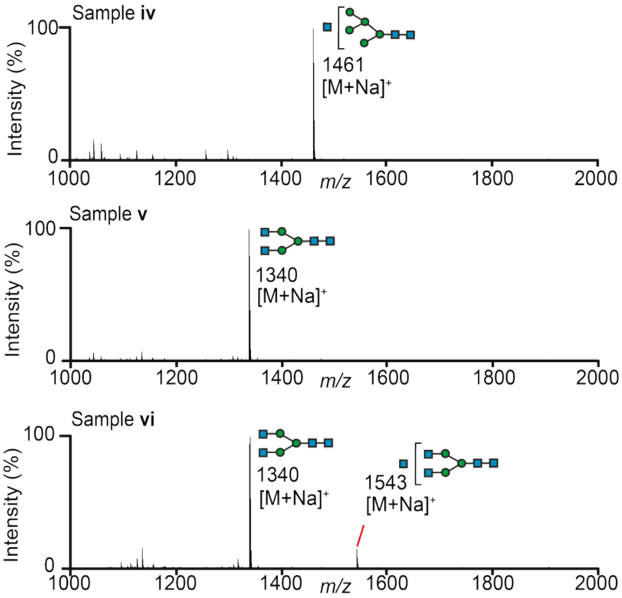


**Supporting Fig. 9**: Released N-glycan MS spectra of Man5-Fc after differential N-glycoprotein tagging with UDP-GlcNAc **1** and WT-MGAT1 (iv), WT-MGAT2 (v) or WT-MGAT5 (vi). Data are from one experiment.

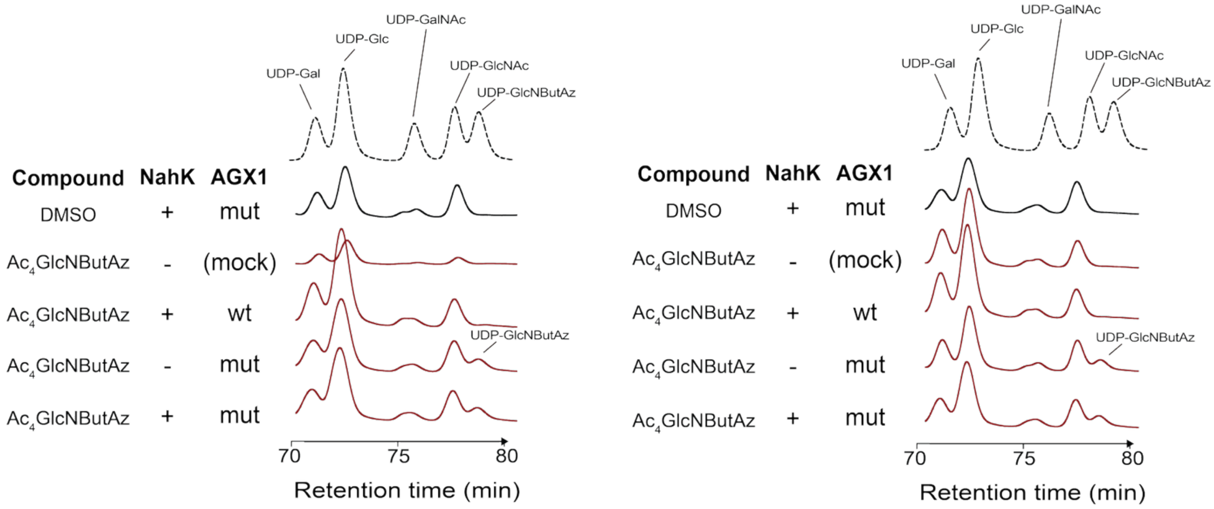


**Supporting Fig. 10:** Biosynthesis of UDP-GlcNButAz **6** in cells stably expressing a combination of NahK, and wildtype or mutant AGX1 (AGX1$^{F383A}$), as assessed by high performance anion exchange chromatography (HPAEC), in comparison with a mixture of synthetic **6** and synthetic UDP-Gal, UDP-Glc. UDP-GalNAc and UDP-GlcNAc (dashed line). Cells were fed with 50 µM $Ac_4$GlcNButAz **12** or DMSO and extracts were assessed by anion exchange chromatography. Experiment was performed in two independent biological replicates.

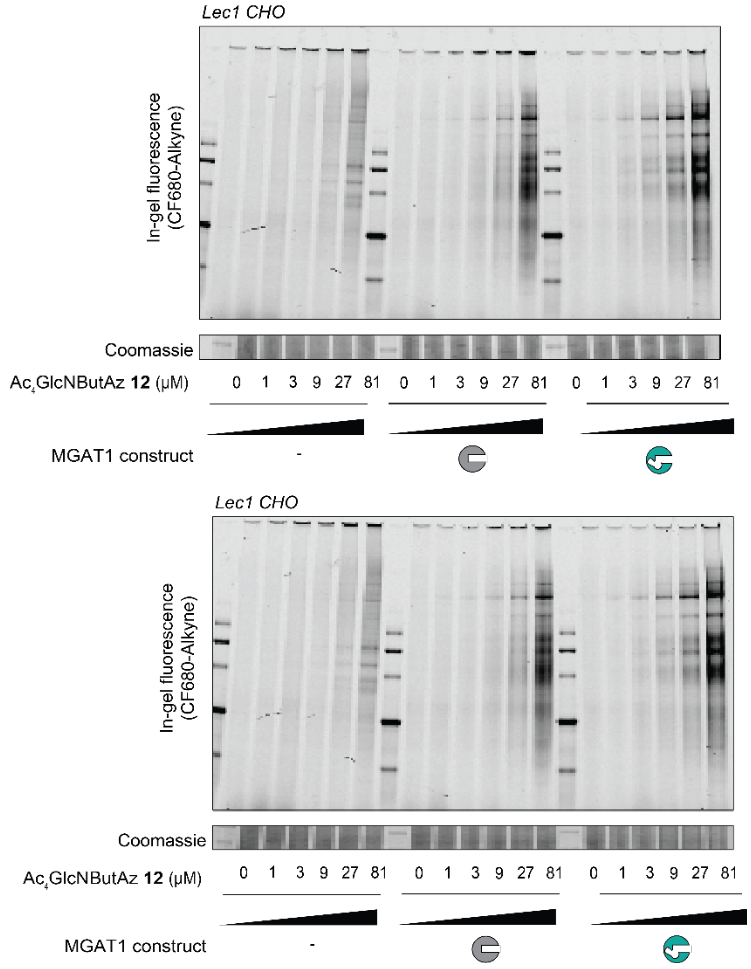


**Supporting Fig. 11:** Cell-surface labeling of Lec1 CHO cells expressing NahK and AGX1$^{F383A}$ alone (left), NahK, AGX1$^{F383A}$ and WT-MGAT1 (center) or NahK, AGX1$^{F383A}$ and BH-MGAT1 (right). Cells were stably transfected with plasmids as indicated, fed with increasing concentrations of Ac$_4$GlcNButAz **12**, subjected to on-cell CuAAC with CF680-alkyne, and fluorescent tagging was assessed by in-gel fluorescence.

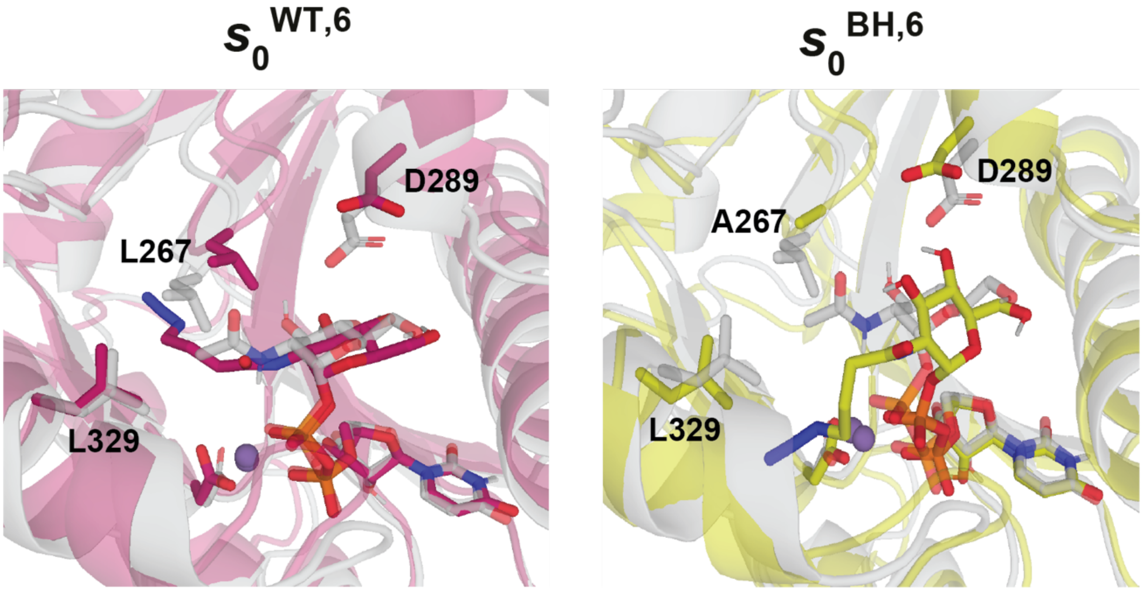


**Supporting Fig. 12:** VAMPnets binding poses comparing the binding of UDP-GlcNButAz **6** to WT- and BH-MGAT1. The two minor states, $s_0^{WT,6}$ and $s_0^{BH,6}$, are shown.

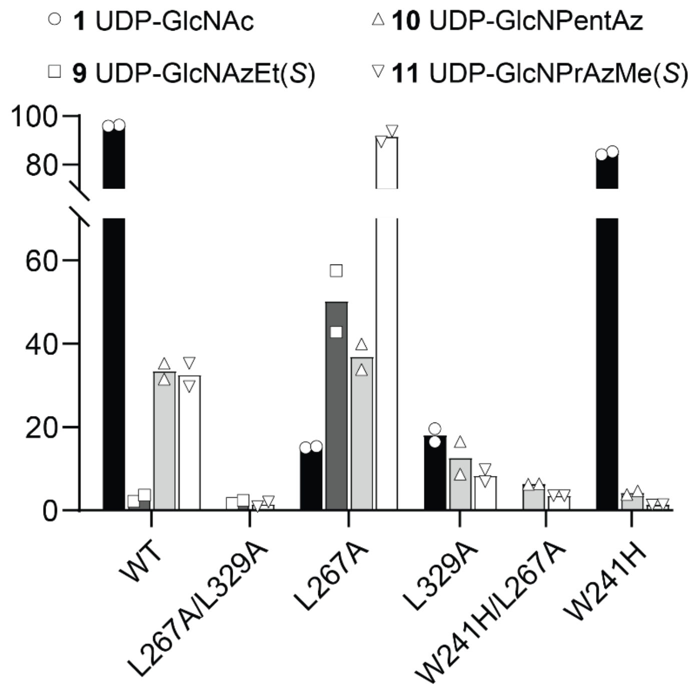


**Supporting Fig. 13:** WT- and BH-MGAT1 variants were subjected to UDP-GlcNAc **9-11** analogs in endpoint enzymatic assays, using Man5-PA as an acceptor substrate and analyzed by UV absorption at 302 nm on UPLC-MS. Data are individual data points and means from two independent replicates.

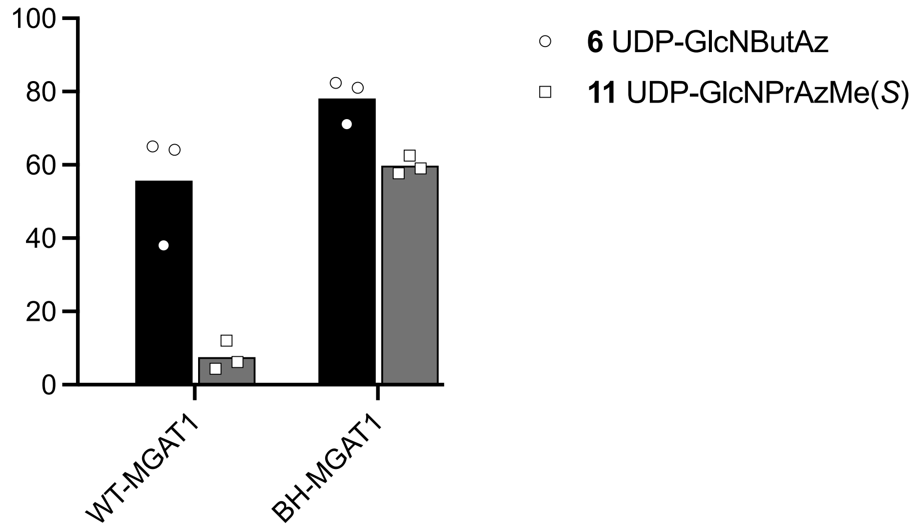


**Supporting Fig. 14**: WT- and BH-MGAT1 (MGAT1$^{L267A}$) were subjected to UDP-GlcNAc analogs **6** and **11** in endpoint enzymatic assays, using Man5**-**PA as an acceptor substrate and analyzed by absorption at 302 nm on UPLC-MS. Data are individual data points and means from three independent replicates.

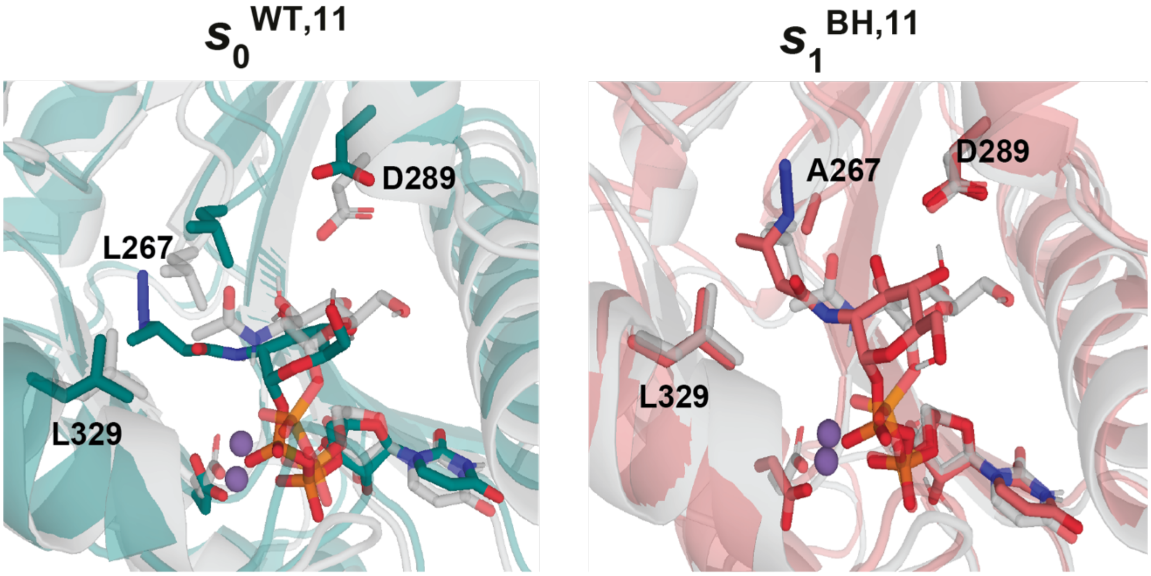


**Supporting Fig. 15:** VAMPnets binding poses comparing the binding of UDP-GlcNButAz **6** to WT- and BH-MGAT1. The two minor states, $s_0^{WT,11}$ and $s_1^{BH,11}$, are shown.

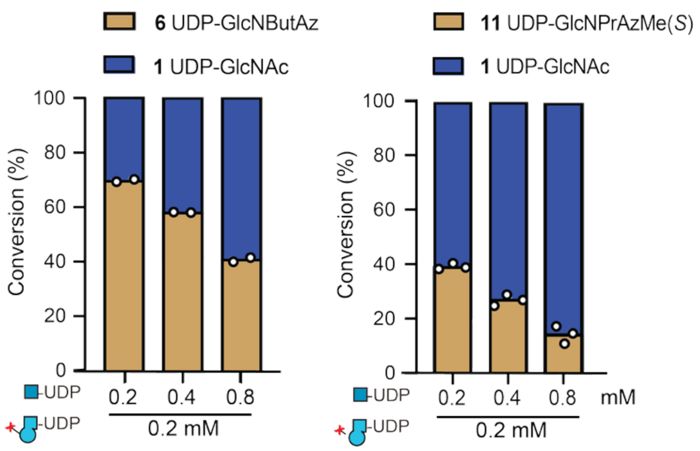


**Supporting Fig. 16:** Wildtype MGAT2 was subjected to UDP-GlcNAc analogs **6** and **11** in endpoint enzymatic assays in the presence of excess UDP-GlcNAc, using A1-PA as an acceptor substrate and analyzed by absorption at 302 nm on UPLC-MS. Data are individual datapoints and means from two (UDP-GlcNButAz) or three (UDP-GlcNPrAzMe(*S*)) independent replicates.

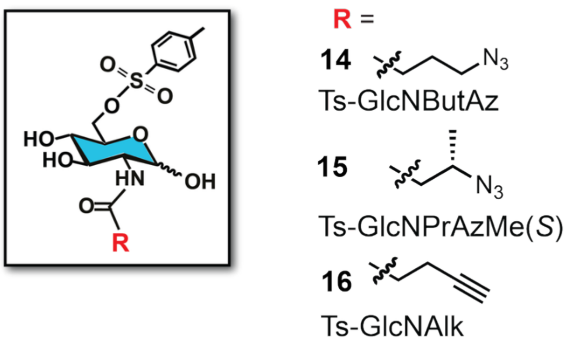


**Supporting Fig. 17:** Collection of GlcNAc analogs **14-16** containing a 6-OTs group.

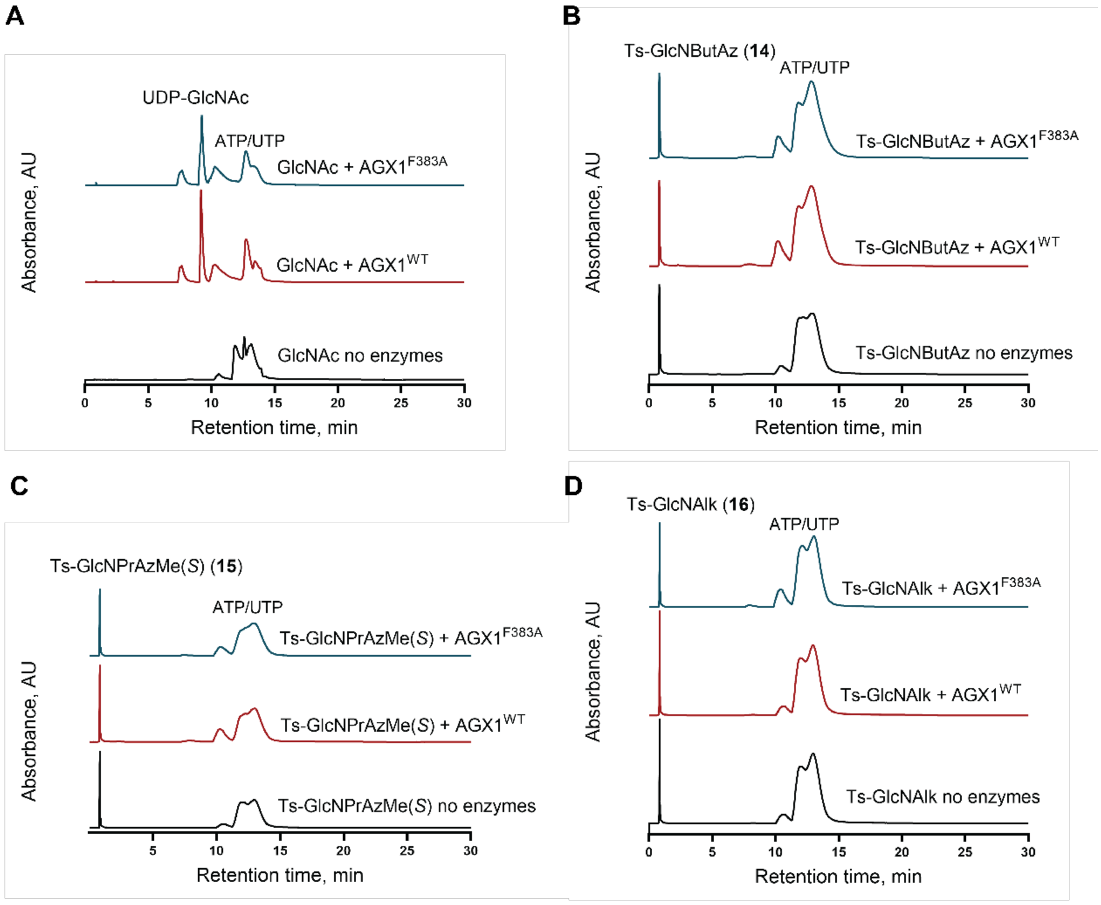


**Supporting Fig. 18:** *In vitro* evaluation of tosylated GlcNAc analogs as substrates for the biosynthetic enzymes NahK and AGX1$^{WT}$ and AGX1$^{F383A}$. GlcNAc (*A*), **14** (*B*), **15** (*C*) and **16** (*D*) were subjected to NahK and WT (red trace) or AGX1$^{F383A}$ (blue) in the presence of ATP and UDP. Product formation was evaluated by the presence or absence of product peak at 260 nm on UPLC-MS. Blanks with the above reaction mixture and without enzymes were included (black). GlcNAc was converted to UDP-GlcNAc indicated with additional peak at 8 min. The tosylated analogs **14**, **15** and **16** were not converted to the corresponding UDP-sugars indicated by the lack of additional UV and ion peaks with the expected *m/z*. UV traces are representative of two independent replicates.

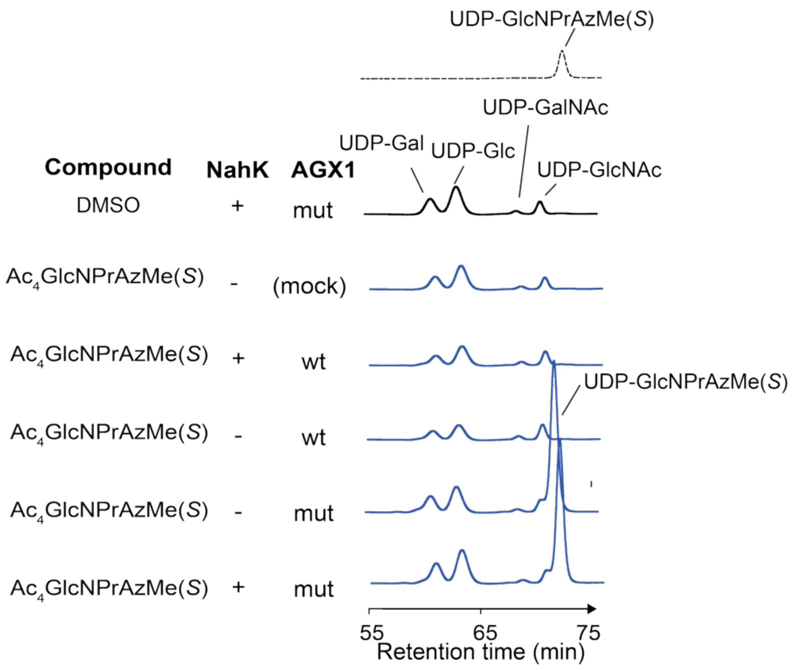


**Supporting Fig. 19:** Second independent biological replicate of biosynthesis of UDP-GlcNPrAzMe(*S*) in cells stably expressing a combination of NahK, and wildtype or mutant AGX1 (AGX1$^{F383A}$), as assessed by HPAEC, in comparison with synthetic **11** (dashed line). Cells were fed with 250 µM Ac$_4$GlcNPrAzMe(*S*) **13** or DMSO and extracts assessed by anion exchange chromatography.

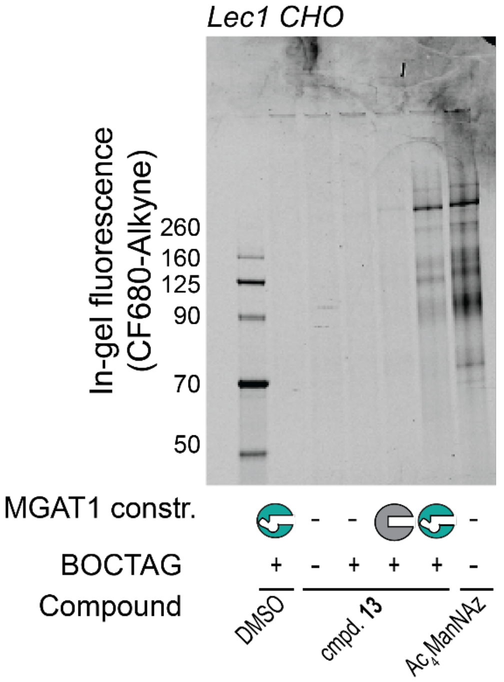


**Supporting Fig. 20:** Second independent biological replicate of cell-surface labeling of Lec1 CHO cells. Cells were stably transfected with plasmids as indicated, fed either DMSO, 9 µM $Ac_4$GlcNPrAzMe(*S*) **13** or 2.5 µM $Ac_4$ManNAz. Cells were subjected to on-cell CuAAC with CF680-alkyne, and fluorescent tagging was assessed by in-gel fluorescence.

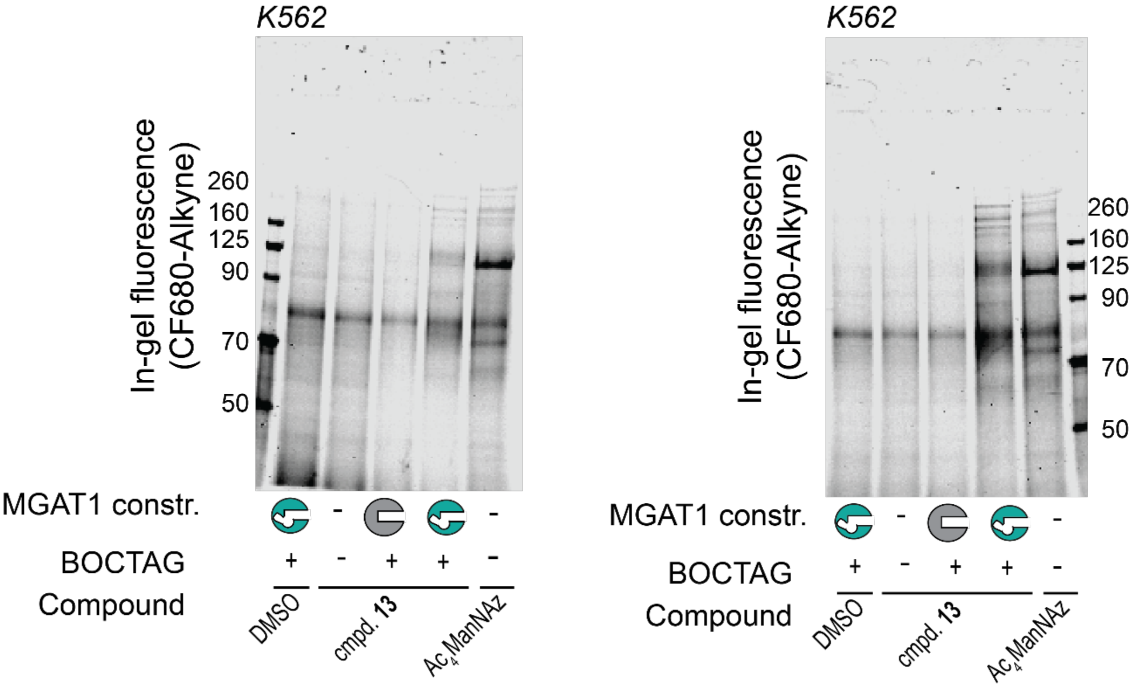


**Supporting Fig. 21** Cell-surface labeling of K562 cells. Cells were stably transfected with plasmids as indicated, fed either DMSO, 9 µM $Ac_4GlcNPrAzMe(S)$ **13** or 2.5 µM $Ac_4ManNAz$. Cells were subjected to on-cell CuAAC with CF680-alkyne, and fluorescent tagging was assessed by in-gel fluorescence. Two independent biological replicates are shown.

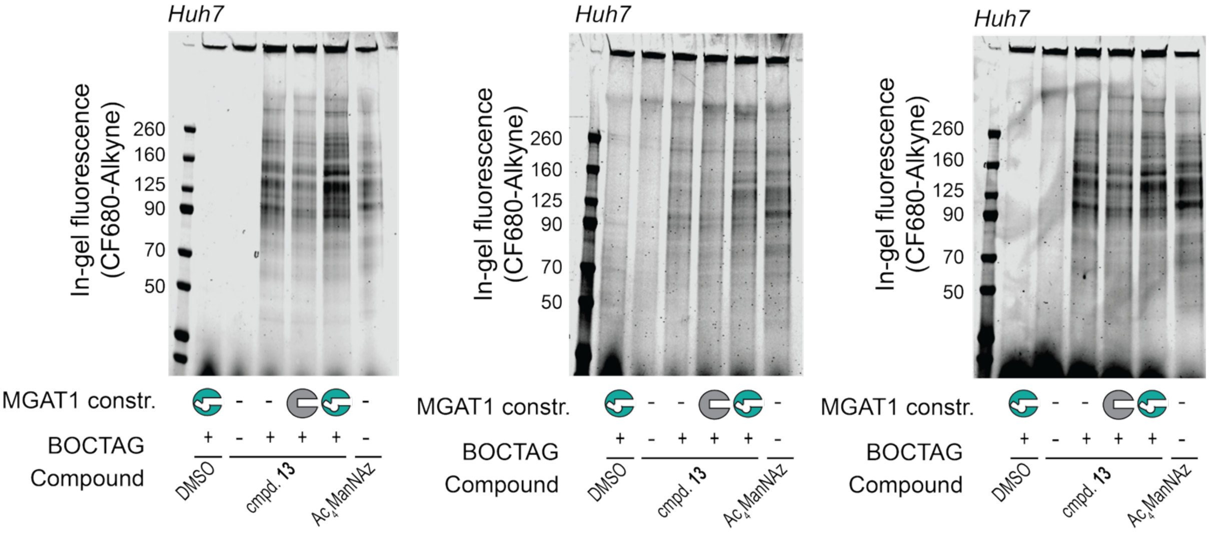


**Supporting Fig. 22:** Cell-surface labeling of Huh7 cells. Cells were stably transfected with plasmids as indicated, fed either DMSO, 27 µM $Ac_4$GlcNPrAzMe(*S*) **13** or 1 µM $Ac_4$ManNAz. Cells were subjected to on-cell CuAAC with CF680-alkyne, and fluorescent tagging was assessed by in-gel fluorescence. Three independent biological replicates are shown.

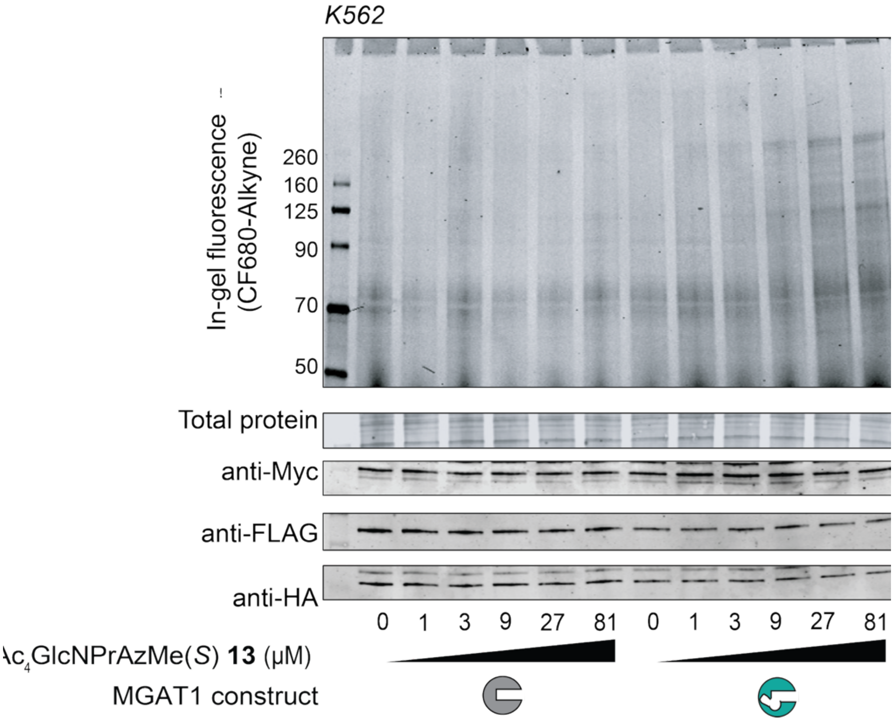


**Supporting Fig. 23** Second independent biological replicate of dose-dependent incorporation of $Ac_4$GlcNPrAzMe(*S*) **13** into K562 cells. Cells were stably transfected with plasmids as indicated, fed increasing concentrations of $Ac_4$GlcNPrAzMe(*S*) **13**, subjected to on-cell CuAAC with CF680-alkyne, and fluorescent tagging was assessed by in-gel fluorescence.

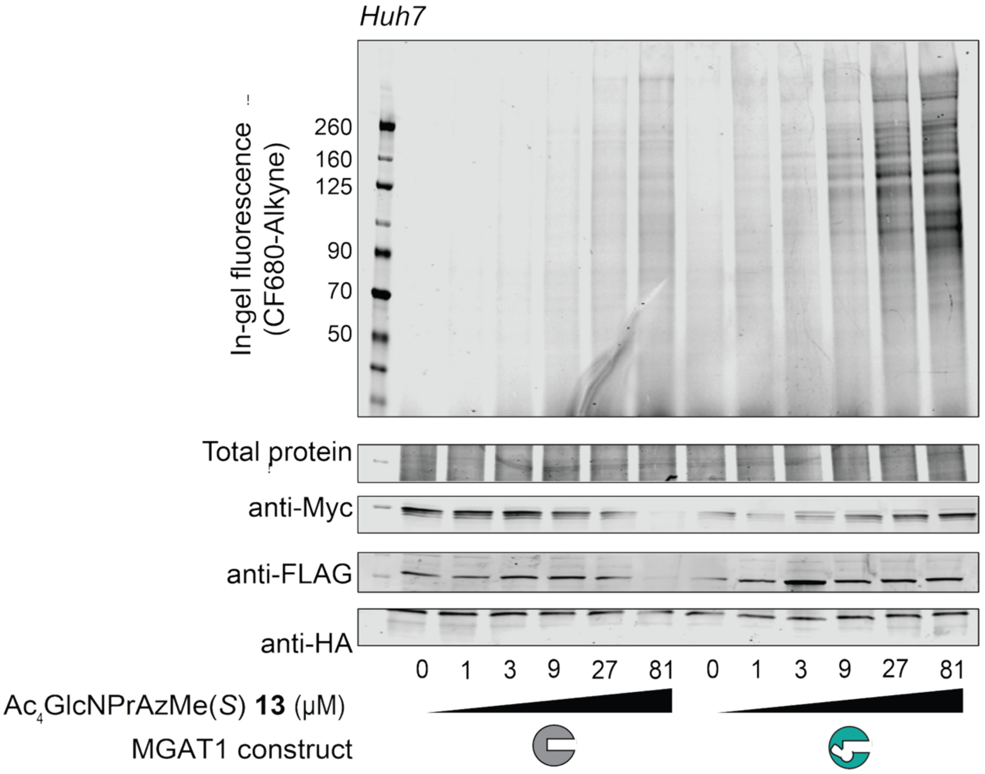


**Supporting Fig. 24** Second independent biological replicate of dose-dependent incorporation of $Ac_4$GlcNPrAzMe(*S*) **13** into Huh7 cells. Cells were stably transfected with plasmids as indicated, fed increasing concentrations of $Ac_4$GlcNPrAzMe(*S*) **13**, subjected to on-cell CuAAC with CF680-alkyne, and fluorescent tagging was assessed by in-gel fluorescence.

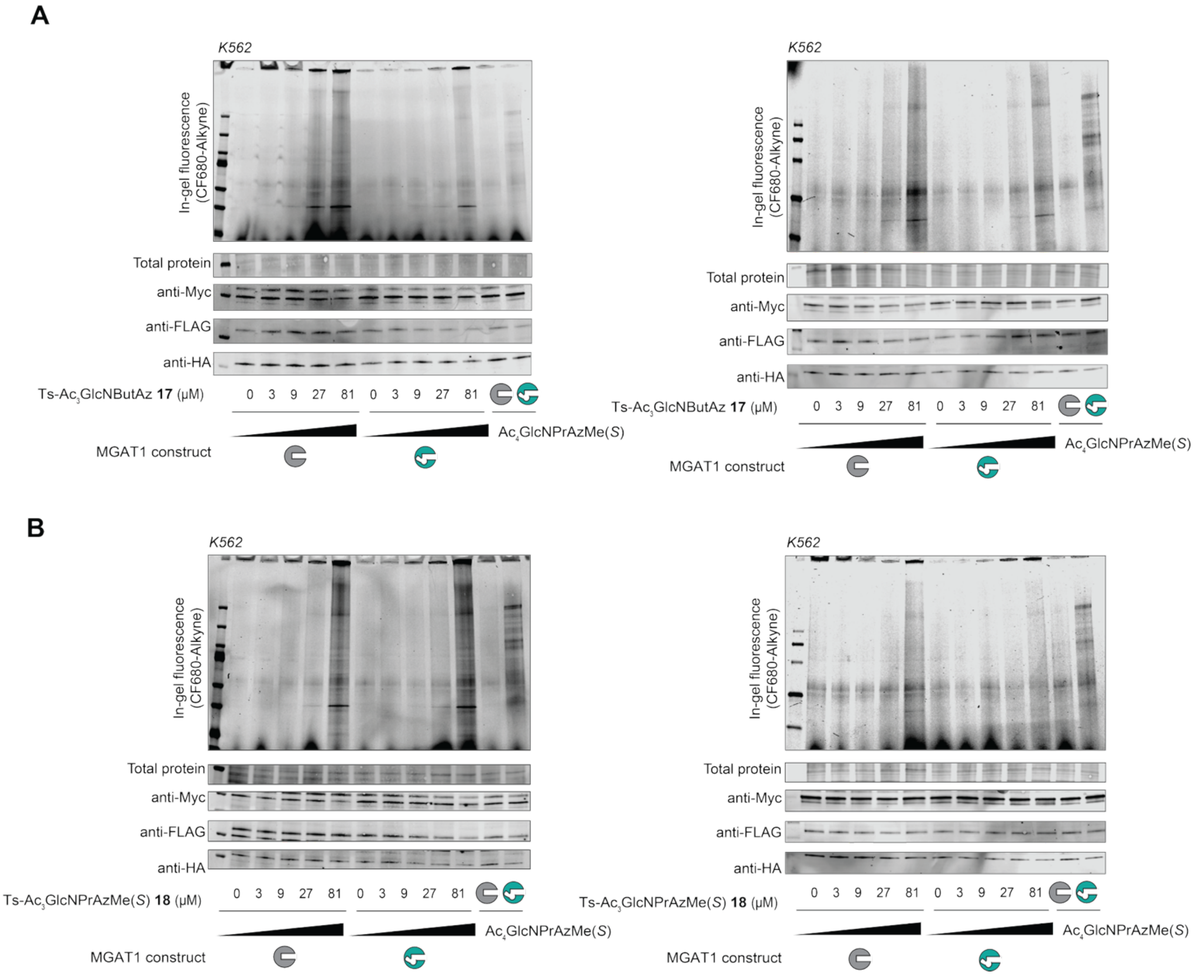


**Supporting Fig. 25:** Cell-surface labeling of K562 cells with tosylated compounds. Cells were stably transfected with plasmids as indicated, fed with increasing concentrations of either Ts-$Ac_3$GlcNButAz **17** (*A*) or Ts-$Ac_3$GlcNPrAzMe(*S*) **18** (*B*) and subjected to on-cell CuAAC with CF680-alkyne. Fluorescent tagging was assessed by in-gel fluorescence. $Ac_4$GlcNPrAzMe(*S*) **13** (9 µM) was used as a positive control. Two independent biological replicates are shown in each panel.

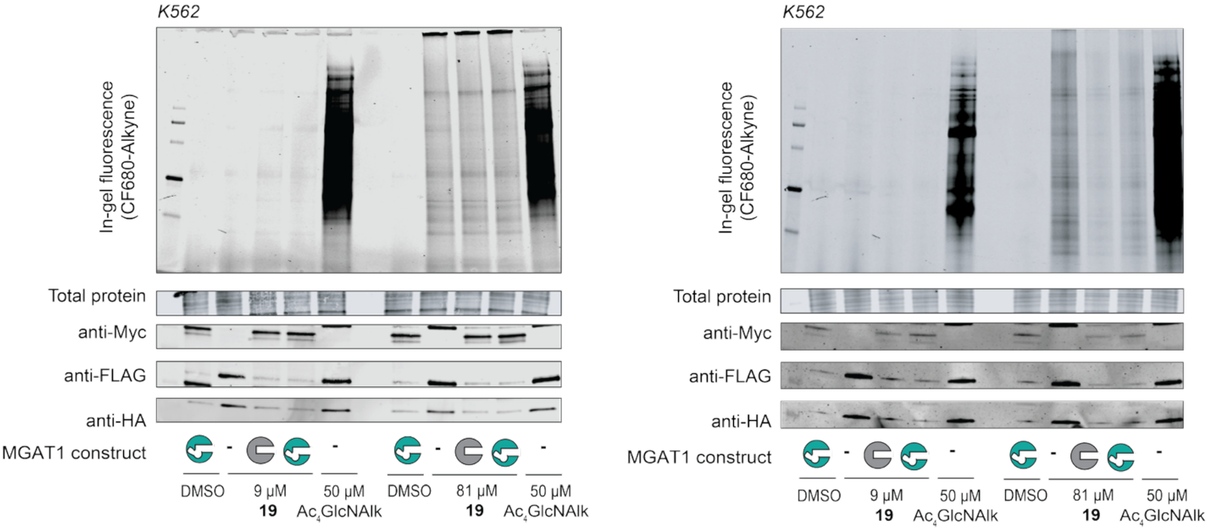


**Supporting Fig. 26:** Cell-surface labeling of K562 cells using Ts-$Ac_3$GlcNAlk **19**. Cells were stably transfected with plasmids as indicated, fed either 9 µM or 81 µM of Ts-$Ac_3$GlcNAlk **19**, subjected to on-cell CuAAC with CF680-alkyne, and fluorescent tagging was assessed by in-gel fluorescence. $Ac_4$GlcNAlk (50 µM) was used as a positive control. Two independent biological replicates are shown.

**Experimentals**

**Supporting Table 1. Oligonucleotides used for site directed mutagenesis**

| | |
|---|---|
| **L267A Forward** | ACTTTTTCCCTGGCGCAGGCTGGCTGCTGTTG |
| **L267A Reverse** | CAACAGCAGCCAGCCTGCGCCAGGGAAAAAGT |
| **L329A Forward** | CAGTTCTTTGACCAGCACGCCAAGTTTATCAAGCTGAAC |
| **L329A Reverse** | GTTCAGCTTGATAAACTTGGCGTGCTGGTCAAAGAACTG |
| **W241H Forward** | GGTGCGTCTCGGCCCATAATGACAACGGCAAG |
| **W241H Reverse** | CTTGCCGTTGTCATTATGGGCCGAGACGCACC |
| **W241HL 267A Forward** | CCACCCTCAGTCAGCGCCTTGGACGGGGATCCT |
| **W241HL 267A Reverse** | GAGGTTGATTGGATCCAAGCTATCAACCACTTTGTACAAGAAAGCTGGGT CCTAGTTCCAGGATGGGTCGTATCC |
| **W241HL 329A Forward** | CCACCCTCAGTCAGCGCTCTTGACGGGGATCCTG |
| **W241HL 329A Reverse** | GAGGTTGATTGGATCCAAGCTATCAACCACTTTGTACAAGAAAGCTGGGT CCTAGTTCCAACTGGGGTCGTACC |
| **W241HL 267A L329A Forward** | CCACCCTCAGTCAGCGCTCTGGACGGTGATCCTG |
| **W241HL 267A L329A Reverse** | GAGGTTGATTGGATCCAAGCTATCAACCACTTTGTACAAGAAAGCTGGGT CCTAGTTCCAAGATGGGTCGTACCC |

**Cloning, expression and purification of truncated MGAT1**

A construct encoding the soluble version of MGAT1 comprising N-terminal $His_8$ tag, N-terminal GFP tag, followed by amino acids 30-445 of human MGAT1 in pGEn2-DEST was from DNASU (HsCD00413193) and originally made by Moremen and colleagues.[1] Single-point mutations of MGAT1 were derived using a Gibson assembly method for site directed mutagenesis. All primers were obtained from Integrated DNA Technologies (Coralville, USA). Gibson solution of MGAT1 variants L267A, L329A, W241H contained 0.5 µM forward primer, 0.5 µM reverse primer, 0.1 ng/µL wild-type MGAT1 in pGEn2-DEST, and 1 U/µL Phusion High-Fidelity DNA Polymerase (Thermo Fisher Scientific, Waltham, USA) in a final reaction volume of 50 µL. Doubly mutated variant L267AL329A was derived from pGEn2 plasmid of MGAT1 L267A using Gibson solution that contained the forward and reverse primers of L329A. Double mutants W241HL267A, W241HL329A, and triple mutant W241HL267AL329A were ordered from Twist Bioscience (South San Francisco, USA) in

Twist plasmids. Fragments of interest were amplified by PCR in a solution containing 0.5 µM forward primer, 0.5 µM reverse primer, 0.4 ng/µL Twist plasmid, and 1 unit/µL Q5 High-Fidelity DNA Polymerase (New England Biolabs, Ipswich, USA) in a final reaction volume of 25 µL. PCR fragments were cloned into a AfeI/BamHI-digested pGEn2-DEST by using the In-Fusion HD Cloning Kit (Takara, Tokyo, Japan). Mutations were verified by whole-plasmid sequencing. A total of 50 µg of plasmid DNA diluted in 2.5 mL of Opti-MEM I Reduced Serum Medium (Thermo Fisher Scientific, Waltham, USA) was mixed with 135 µL of Expifectamine transfection reagent (Thermo Fisher Scientific, Waltham, USA) diluted in 2.5 mL of Opti-MEM medium according to manusfacturer's instruction, before transfectioninto 50 mL of Expi293F® cells (Thermo Fisher Scientific, Waltham, USA) grown to a density of 3.5 million cells/mL at higher than 97% viability rate.  MGAT1 constructs were expressed and secreted to the cell medium. After 72 h, the supernatants were purified over Ni-NTA Agarose beads (QIAGEN, Hilden, Germany) exactly as described before for WT-MGAT1.[2] On average, approx. 0.34 mg recombinant enzyme was obtained from 100 million cells through this protocol, with no obvious difference in expression levels between different MGAT1 constructs.

##### Expression and purification of truncated MGAT2

A construct encoding the soluble version of MGAT2 comprising N-terminal $His_8$ tag, N-terminal GFP tag, followed by amino acids 30-447 of human MGAT2 in pGEn2-DEST was from DNASU (HsCD00413203) and originally made by Moremen and colleagues.[1] A total of 50 µg of plasmid was used to transfect 50 mL of Expi293F® cells at a density of 3.5 million cells/mL and higher than 97% viability rate. MGAT2 construct was expressed and secreted to the cell medium. After 72 h, the supernatant was purified over Ni-NTA Agarose beads exactly as described before for WT-MGAT2.[2] On average, approx. 0.39 mg recombinant MGAT2 was obtained from 100 million cells through this protocol. Enzyme fractions were buffer exchanged to 20 mM HEPES pH 7.5 with 200 mM NaCl, 1 mM DTT and 20% Glycerol, aliquoted and flash-frozen in liquid nitrogen and stored at -80 °C.

##### Expression and purification of truncated MGAT5

A double mutant of MGAT5, F458V/F517L, which is able to specifically make use of a 4-azidobutyramide-containing UDP-GlcNAc analog, was cloned into a pOMNIBac plasmid, expressed from High Five cells (*Trichoplusia ni*) and purified from HisTrap column (Cytiva, Marlborough, USA) as described before. [2,3]

##### *In vitro* glycosylation assay using synthetic glycan acceptor substrate

Commercial procainamide-labelled pentamannosylated N-glycan precursor (Man5-PA; Ludger Ltd, Abingdon, UK) was used as an acceptor substrate for MGAT1 to assess conversion by near UV (302 nm) absorbance-based assay. Reactions were carried out using 1 mM UDP-GlcNAc or UDP-GlcNAc analogs **2-8** or analog **11**, 0.2 mM Man5-PA, and 250 nM MGAT1 constructs, in a total volume of 5 µL Reaction Buffer I (50 mM MES pH 6.5, 25 mM NaCl, 5 mM $MnCl_2$) at 37 °C for 30 min. UDP-GlcNAc analogs **9-11** were treated in the same way, but reactions were stopped after 16 h. For competition experiment, reactions were performed using 0.1 mM Man5-PA, 0.2 mM UDP-GlcNAc analog and either 0.2 mM, 0.4 mM or 0.8 mM UDP-GlcNAc, and 250 nM MGAT1 constructs, in a total volume of 10 µL

Reaction Buffer I at 37 °C for 30 min (for analog **6**) or 16 h (for analog **11**). Reactions were stopped by addition of an equal volume of acetonitrile, followed by centrifugation at 13,000 x g, at 4 °C for 30 min. Of each sample, 9 µL of the supernatants were injected onto an Acquity H-Class PLUS QDa UPLC-MS (Waters, Milford, USA) equipped with an ACQUITY UPLC Glycan BEH Amide column (130 Å, 1.7 µm, 2.1 x 100 mm, Waters, Milford, USA). Samples were run at flow rate of 0.35 mL/min using Buffer A (10 mM ammonium formate at pH 4.5); Buffer B (10 mM ammonium formate in ACN: water 90:10 (v/v)) with a gradient of 90-55% buffer B over 17 min. The percentage of turnover of procainamide-labelled substrate glycan into the corresponding product glycan was calculated by integration of the near-UV peak (302 nm absorption) of the acceptor substrate and the product glycan or by integration of the MS peaks in the scenario where acceptor substrate and the product glycan co-eluted, and determined as Turnover % = Peak Area of Product Glycan / Peak Area of (Substrate Glycan + Product Glycan) %.

Commercial procainamide-labelled A1 N-glycan precursor (A1-PA; Ludger Ltd, Abingdon, UK) was used as an acceptor substrate for MGAT2 to assess conversion by near UV (302 nm) absorbance-based assay. Reactions were carried out using 1 mM UDP-GlcNAc or UDP-GlcNAc analogs **2-8**, 0.2 mM A1-PA, and 130 nM MGAT2, in a total volume of 5 µL Reaction Buffer II (50 mM Tris pH 6.8, 5 mM $MnCl_2$, 0.1% BSA in PBS, 0.1% Triton) at 37 °C for 16 h. For competition experiment, reactions were performed using 0.1 mM A1-PA, 0.2 mM UDP-GlcNAc analog and either 0.2 mM, 0.4 mM or 0.8 mM UDP-GlcNAc, and 130 nM MGAT2, in a total volume of 10 µL Reaction Buffer II at 37 °C for 16 h. Reactions were stopped by addition of an equal volume of acetonitrile, followed by centrifugation at 13,000 x g, at 4 °C for 30 min. Of each sample, 9 µL of the supernatants were injected onto an Acquity H-Class PLUS QDa UPLC-MS equipped with an ACQUITY UPLC Glycan BEH Amide column (130 Å, 1.7 µm, 2.1 x 100 mm). Samples were run at flow rate of 0.35 mL/min using Buffer A (10 mM ammonium formate at pH 4.5); Buffer B (10 mM ammonium formate in ACN: water 90:10 (v/v)) with a gradient of 90-55% buffer B over 17 min. The percentage of turnover of procainamide-labelled substrate glycan into the corresponding product glycan was calculated by integration of the near-UV peak (302 nm absorption) of the acceptor substrate and the product glycan, and determined as Turnover % = Peak Area of Product Glycan / Peak Area of (Substrate Glycan + Product Glycan) %.

**Enzyme kinetics**

Recombinant WT-MGAT1 at a concentration of 278 nM was incubated for 30 min with serially diluted UDP-GlcNAc or UDP-GlcNAc analogs **6** or **11** from 0.02 to 1 mM and 0.2 mM Man5-PA in Reaction Buffer I in a total volume of 5 µL, and turnover measured by UPLC-MS as described above. To quantify the formation of product, standard curves were employed by plotting concentrations of serially diluted Man5-PA against the integrated peak area under the near-UV peak (302 nm absorption). The kinetics curve was plotted with Prism 10 (GraphPad, San Diego, USA) and fitted with a Michaelis Menten function to calculate $k_{cat}$, $K_M$ and $V_{max}$. Recombinant BH-MGAT1 at a concentration of 195 nM was incubated for 30 min with serially diluted UDP-GlcNAc analog **6** from 0.02 to 1 mM or analog **11** from 0.01 to 0.4 mM and 0.2 mM Man5-PA in Reaction Buffer I in a total volume of 5 µL, and turnover measured by UPLC-MS as described above. To quantify the formation of product, standard curves were employed by plotting concentrations of serially diluted Man5-PA against the integrated peak area under the near-UV peak (302 nm absorption). The kinetics curve was plotted with Prism 10 and fitted with a Michaelis Menten function to calculate $k_{cat}$, $K_M$ and $V_{max}$.

##### Bioorthogonal labelling of Man5-containing glycoprotein substrate by BH-MGAT1, WT-MGAT2, and BH-MGAT5

Monomeric antibody Fc containing a single Man5 N-glycan per polypeptide was expressed in the *P. pastoris* SuperMan5 (Biogrammatics, Carlsbad, USA) as described previously. *In vitro* glycosylation was performed using 20 µg Man5-Fc in 30 µL reaction volume containing 50 mM MES pH 6.5, 5 mM NaCl, 1 mM $MnCl_2$, 0.2 mM $ZnCl_2$ and 1 mM UDP-GlcNAc at 37 °C for 16 h in the presence of 250 nM WT-MGAT1 or in the presence of 250 nM BH-MGAT1 and 200 µM UDP-GlcNAc analog **6**. *In vitro* elaboration was performed using 20 µg Man5-Fc in 30 µL reaction volume containing 50 mM MES pH 6.5, 5 mM NaCl, 1 mM $MnCl_2$, 0.2 mM $ZnCl_2$ and 1 mM UDP-GlcNAc at 37 °C for 16 h in the presence of 250 nM WT-MGAT1 and 250 nM drosophila α-mannosidase (MAN2A1, which was a gift from Zach Armstrong), followed by incubation with either 1 mM UDP-GlcNAc, 0 or 200 µM UDP-GlcNAc analog **6** in the presence of 250 nM WT-MGAT2 in a total 40 µL reaction volume. Alternative *in vitro* elaboration was performed using 20 µg Man5-Fc in 30 µL reaction volume containing 50 mM MES pH 6.5, 5 mM NaCl, 1 mM $MnCl_2$, 0.2 mM $ZnCl_2$ and 1 mM UDP-GlcNAc at 37 °C for 16 h in the presence of 250 nM WT-MGAT1, 250 nM drosophila α-mannosidase, and 250 nM WT-MGAT2, followed by incubation with either 1 mM UDP-GlcNAc, either 0 or 200 µM UDP-GlcNAc analog **6** in the presence of 500 nM $MGAT5^{F458V/F517L}$ (BH-MGAT5)[2]. Reactions were heat-inactivated at 95 °C for 20 s and cooled to 4 °C. Azide-containing reaction mixtures were sequentially treated with 50 µM biotin-alkyne (Biotium, Fremont, USA), 1200 µM BTTAA (Click Chemistry Tools, Scottsdale, USA), 600 µM $CuSO_4$, 5 mM aminoguanidinium chloride and 5 mM sodium ascorbate (final concentrations). Reactions were quenched by the addition of 0.5 M EDTA. Reaction mixtures were then subjected to SDS-PAGE and blotted on nitrocellulose membranes. The total protein amount was assessed using the REVERT protein staining kit (LI-COR Biosciences, Lincoln, USA), and biotinylation was detected using IRDye 800CW Streptavidin (LI-COR Biosciences, Lincoln, USA) according to the manufacturer's instructions.

##### Glycan release and ion mobility–mass spectrometry (IM-MS) analysis of bioorthogonally labelled Man5-Fc

N-glycans were released from 50 µg of protein with 2 µg PNGaseF (produced in-house) at 37 °C for 16 h. Released N-glycans were desalted using a 0.6 µL C18 ZipTip (Merck Millipore, Darmstadt, Germany) overlaid with porous graphitized carbon resin (HyperSep™ Hypercarb™ Thermo Fisher Scientific, Waltham, USA). De-salted N-glycans were lyophilized and resuspended in 50% methanol immediately prior to IM-MS analysis. IM-MS measurements were performed on a SYNAPT XS IMS instrument (Waters, Manchester, UK). For each sample analysis, 2 µL of N-glycan sample material was ionized by nano-electrospray ionization (nano-ESI) from gold-coated borosilicate glass capillaries prepared in-house.[4] Data was acquired in positive ion mode with the settings as follows: capillary voltage 1.4 kV, cone voltage 150 V, source offset voltage 5 V, source temperature 100 °C. Collision-induced dissociation was performed in the transfer with argon as the collision gas with a collision energy (CE) between 50-150 V. Data acquisition was carried out using Waters MassLynx™ (version 4.2) and annotation of spectra was done manually.

**Bioorthogonal labelling of Man5-containing cellular fraction by BH-MGAT1**

The Lec1 Chinese hamster ovary (CHO) cell line was a kind gift from Pamela Stanley (Albert Einstein College of Medicine, New York, USA).[5,6] One 75-$cm^2$ flask of cells were grown to confluency in MEMα medium (Fisher Scientific UK Ltd, Loughborough, UK) containing 10% (v/v) fetal bovine serum (FBS), 100 U/mL penicillin-streptomycin (Thermo Fisher Scientific, Waltham, USA). Cells were rinsed three times with phosphate-buffered saline, harvested at 500 x g for 5 min. Pelleted cells were resuspended in ice-cold PBS and transferred to a 1.5 mL microcentrifuge tube, centrifuged at 500 x g, 4 °C, for 3 min and supernatant removed. The Subcellular Protein Fractionation Kit for Cultured Cells (Thermo Fisher Scientific, Waltham, USA) was used to obtain the membrane fraction. Protein concentration was determined using the Rapid Gold BCA Protein Assay Kit (Thermo Fisher Scientific, Waltham, USA). The membrane fraction was concentrated to 16 mg/mL using an Amicon Ultra-0.5 centrifugal filter (Merck Millipore, Darmstadt, Germany).

*In vitro* glycosylation reactions were performed using 60 µg Lec1 membrane fraction in 25 µL reaction volume containing 200 mM MES pH 6.5, 25 mM NaCl, 5 mM $MnCl_2$, 200 µM UDP-GlcNAc analog and either 200 or 800 µM UDP-GlcNAc at 37 °C for 16 h in the presence of 500 nM WT- or BH-MGAT1. Reactions were heat-inactivated at 95 °C for 20 s and cooled to 4 °C. Azide-containing reaction mixtures were sequentially treated with one-third volume of 400 µM biotin-alkyne, 4800 µM BTTAA, 1200 µM $CuSO_4$, 20 mM aminoguanidinium chloride and 20 mM sodium ascorbate (final concentrations 100 µM biotin probe, 1200 µM BTTAA, 300 µM $CuSO_4$, 5 mM aminoguanidinium chloride and 5 mM sodium ascorbate). Reaction mixtures were then subjected to SDS-PAGE and blotted on nitrocellulose membranes. The total protein amount was assessed using the REVERT protein staining kit, and biotinylation was detected using IRDye 800CW Streptavidin according to the manufacturer's instructions.

**Molecular Dynamics**

The computational models were based on WT-MGAT1 (RCSB PDB: 2APC). The crystallographic C-linked GlcNAc-UDP was converted to the O-linked natural substrate and in silico mutation L267A was made to form BH-MGAT1 complex. Ligands were docked into both variants with their common substructures kept rigid and the natural substrate as reference, using AutoDock Vina.[7,8] For each ligand–enzyme pair, the top-ranked pose by predicted affinity seeded the MD simulations. Enzymes, carbohydrate fragments, and non-sugar ligand moieties were described with Amber ff14SB[9], GLYCAM[10], and GAFF2[11], respectively; solvent was TIP3P water[12]. Protonation states were assigned by H++[13,14] at pH 6.5. Crystal waters were retained, and each complex was embedded in a periodic solvent box with at least 10 Å padding from any protein atom, then neutralized with $Na^+$ /$Cl^-$ to a bulk concentration of 0.25 mM (12 $Na^+$ and 16 $Cl^-$). Topologies were prepared with AmberTools23 leap[15], converted to GROMACS format using ACPYPE[16,17], and simulated with GROMACS 2022.3[18,19]. Energy minimization used steepest descent for up to 50,000 steps or until the maximum force fell below 10 kJ $mol^{-1}$ $nm^{-1}$. NVT equilibration employed the V-rescale thermostat[20] at 300 K with coupling time $\tau_T$ = 0.1 ps for 2 ns, followed by 2 ns of NPT equilibration with a Parrinello–Rahman barostat[21] at 1 bar, and $\tau_P$ = 2.0 ps. Harmonic position restraints (1000 kJ $mol^{-1}$ $nm^{-1}$) were applied to heavy atoms pre-production. Production MD used a 2 fs time step with all bonds to hydrogens constrained by LINCS, PME electrostatics with default Coulomb and Lennard-Jones cutoffs and long-range dispersion corrections, the Verlet cutoff scheme, and periodic boundary conditions in all directions with the neighbor list updated every 10 steps. For each ligand–enzyme variant pair, five independent replicas of 1 μs were run, yielding 40 μs total sampling (4 ligands × 2 variants × 5 replicas = 40 trajectories).

Trajectory frames were featurized as protein–ligand contact maps (40 Å cutoff) with an exponential distance transform (σ = 2.0 Å), using the first frame of each replica as reference. A two-state VAMPnet[22] (input dimension 342; MLP 342–128–64–32–32–2 with dropout[23] 0.1 on the first two hidden layers) was trained for 300 epochs (batch size 428, learning rate $10^{-5}$) at system-specific lag times τ. We turned the network's outputs at lag time τ into discrete state labels and used these to construct a two-state MSM. Implementation used PyTorch[24] and deeptime[25]; training was deterministic (seed 0) and ran on CPU. For comparison, all stationary probabilities and kinetics are reported at a common 2 lag of 25 ns. VAMP-2 scores, implied-timescale curves, and Chapman–Kolmogorov tests guided hyperparameter and lag selection. The system-specific training lags were: **1**–WT: 2.5 ns, **1**–BH: 25 ns, **6**–WT: 25 ns, **6**–BH: 10 ns, **11**–WT: 25 ns, and **11**–BH: 2.5 ns. RBFE calculations were performed with OpenFE[26] using OpenMM[27] as the MD driver. A ligand–ligand alchemical network over **1, 6,** and **11** was constructed as a minimal-spanning network, scored by LoMAP[28] and seeded with the RDKit[29] MCS. For each edge, transformations were run in both solvent and complex legs against WT and BH. Small-molecule parameters used OpenFF 2.1.1[30] with AM1–BCC[31] partial charges; proteins used Amber ff14SB and water was TIP3P. The OpenFE solvent component employed $Na^+$ /$Cl^-$ at 0.25 mM with neutralization, and all simulations were conducted at 300 K using the Langevin middle integrator[32]. Each window comprised 5,000 steps of minimization, 0.5 ns of equilibration, and 10 ns of production. We used 20 λ windows per leg, and each edge was repeated three times with independent initializations, yielding 20 × 10 ns × 2 legs = 400 ns per repeat and 1.2 μs total sampling per transformation.

##### Cloning of full-length MGAT1 into pSBbi plasmids

Human MGAT1 gene in a pDONR221 plasmid was from DNASU (HsCD00041466) and a pUC57 plasmid encoding MGAT1 L267A was ordered from GenScript. A myc tag was appended to the C-terminus of the MGAT1 constructs and fragments of interest were amplified by PCR and in a solution containing 0.5 μM forward primer, 0.5 μM reverse primer, 0.4 ng/μL parental plasmid, and 1 U/μL Q5 High-Fidelity DNA Polymerase in a final reaction volume of 25 μL. PCR fragments were cloned into a SfiI-digested pSBbi plasmid that encodes $AGX1^{F383A}$ and NahK by using the In-Fusion HD Cloning Kit. Mutations were verified by whole-plasmid sequencing.

##### Generation of cell lines overexpressing wildtype or L267A variant of MGAT1

Human lymphoblast K562 cells, Chinese hamster ovary Lec1 cells and hepatocarcinoma Huh7 cells were used for stable transfection with pSBbi MGAT1/$AGX1^{F383A}$/NahK, pSBbi $MGAT1^{L267A}$/$AGX1^{F383A}$/NahK plasmids or empty pSBbi-GH. A total of 2.4 μg of pSBbi plasmid and 125 ng of pCMV(CAT)T7-SB100 plasmid DNA were used to transfect 800,000 cells seeded at 500,000 cells/mL in a 6-well plate using Lipofectamine LTX (Thermo Fisher Scientific, Waltham, USA) according to the manufacturer's instructions. Stable cell lines were generated via antibiotic selection. Specifically, K562, Lec1, and Huh7 cells were each subjected to selection in RPMI1640, MEMα, and DMEM growth media supplemented with 10% (v/v) fetal bovine serum (FBS), 100 μg/mL penicillin-streptomycin, and hygromycin B (Thermo Fisher Scientific, Waltham, USA) was used at 150 μg/mL, 600 μg/mL, and 100 μg/mL respectively. The selection process commenced 24 hours post-transfection and continued for 7–10 days until stable polyclonal populations were established. Cells were maintained at 37 °C in a humidified atmosphere containing 5% $CO_2$.

##### Cellular biosynthesis of UDP-GlcNAc analogs

K562 cells were seeded at 500,000 cells/mL in 10 cm diameter petri dishes in 10 mL of RPMI growth media supplemented with 10% (v/v) FBS, 100 μg/mL penicillin-streptomycin and incubated for 16 h with either DMSO, 50 μM $Ac_4$GlcNButAz **12** or 250 μM $Ac_4$GlcNPrAzMe(*S*) **13**. After 16-h incubation, cells were transferred to 15 mL falcon tube and centrifuged for 5 min at 500 x g at 4 °C. Media was aspirated and cells were resuspended in 1 mL PBS (without $MgCl_2$ and $CaCl_2$) and transferred to 1.5 mL microcentrifuge tubes. After centrifugation for 5 min at 300 x g, cells were washed two more times with PBS. Cell pellets were mixed with Zirconia/silica beads (0.1 mm, BioSpec, Bertlesville, USA) at a similar volume and 1 mL of 1:1 ethanol/milliQ water was added. Cells were lysed by 3 cycles of vortex shaking for 20 s and were placed on ice for 30 s between shaking steps. Samples were centrifuged for 10 min at 14,000 x g at 4 °C, and the resulting supernatant was transferred to a 1.5 mL tube. The solvent was removed by SpeedVac. The residue was resuspended in 0.3 mL of milliQ $H_2O$ and the solution was membrane-filtered for 40 min at 14,000 x g using a 3 kDa cut-off filter (Centricon, Merck, Darmstadt, Germany). 100 μL of water was add the filter and centrifuged for a further 20 min. The washing step with water was performed twice. The filters were pre-washed three times with 200 μL of Milli-Q prior to use. The flow-through was concentrated by SpeedVac and resuspended in 60 μL of Milli-Q water. Alkaline phosphatase treatment was performed on 20 μL of each lysate. A 200 μU/μL solution of calf intestinal alkaline phosphatase (CIAP, ThermoFisher) was prepared by diluting 1 μL of 20 U/μL CIAP in 99 μL of dilution buffer (25 mM tris pH=7.6, 0.1mM $ZnCl_2$, 1mM $MgCl_2$, 50% glycerol). To 20 μL of each lysate, 2 μL of CIAP (200 μU/μL) was added, and incubated for 1 h at 37 °C. The reaction was quenched by heating at 65 °C for 15 min and centrifuged at 18,000 x g for 5 min. The supernatant was injected onto an ArcPremier water HPLC equipped with a Dionex CarboPac PA1 IC column (2 mm x 150 mm, Thermo Fisher Scientific, Waltham, USA, Cat#057178). Samples were run at a flow rate of 0.25 mL/min using buffer A (1 M NaOAc/1 mM NaOH); buffer B (1 mM NaOH) with the following gradient: 0 min, 5% A, 95% B; 20 min 40% A, 60% B; 60 min, 40% A, 60% B; 63 min, 50% A, 50% B; 87 min, 80% A, 20% B; 95 min, 80% A, 20% B; 96 min, 5% A, 95% B; 101 min, 5% A, 95% B.

##### Immunofluorescence microscopy

Stably transfected Huh7 cells were imaged using fluorescence microscopy. Cells were plated in DMEM medium supplemented with 10% (v/v) FBS, 100 μg/mL penicillin-streptomycin in 12-well plates containing circular 15 mm coverslips with 0.17 mm thickness and left to incubate for 24 h (37 °C, 5% $CO_2$). Coverslips were then transferred to a clean 12-well plate without growth medium and washed twice with DPBS (containing 100 mg / L $CaCl_2$ and 100 mg/mL $MgCl_2$) before fixing with ice-cold 4% (v/v) formaldehyde (Thermo Scientific, Loughborough, UK) in PBS (without $CaCl_2$ or $MgCl_2$) for 15 min at r.t. in the dark. Cells were washed twice with DPBS with 2% (w/v) BSA, before quenching with a 5-min incubation in 50 mM ammonium chloride in DPBS, followed by three further washes in DPBS with 2% BSA. Permeabilization was carried out with 10 min on-ice incubation with 0.5% Tween-20 (Sigma-Aldrich, Burlington, USA) in DPBS with 2% BSA in the dark. After three more washes with DPBS cells were blocked at 4 °C overnight in a solution of DPBS with 3%

donkey serum (Abcam, Cambridge, UK), 2% BSA, and 0.1% Tween-20 (blocking solution). Cells were incubated overnight at 4 °C with the primary antibodies (see table below) diluted in blocking solution. Cells were washed three times in DPBS with 1% donkey serum, 2% BSA, and 0.1% Tween-20, before incubation in the secondary antibodies (see table below) in the same buffer for 1 h at 37 °C in the dark. The cells were washed again three times in DPBS with 0.1% Tween-20 and incubated for 30 min at r.t. with DAPI (1:1000, Thermo Fisher Scientific, Waltham, USA, Cat#62248) in the same buffer. The cells were washed in PBS three times before mounting onto glass slides using ProLong Gold antifade reagent (Thermo Fisher Scientific, Waltham, USA, Cat#10144). Cells were imaged using a VisiTech iSIM microscope.

Primary Antibodies:

| Reference | Supplier | Anti- | Host | Mono/Polyclonal | Dilution |
|---|---|---|---|---|---|
| 05-724-25UG | Sigma-Aldrich | Anti-c-Myc | Mouse | Monoclonal | 1:800 |
| ab52649 | abcam | GM130 | Rabbit | Monoclonal | 1:200 |

Secondary Antibodies:

| Reference number | Supplier | Anti- | Host | Fluorophore | Dilution |
|---|---|---|---|---|---|
| A21206 | Thermo Fisher | Rabbit | Donkey | Alexa Fluor 488 | 1:1000 |
| A31571 | Thermo Fisher | Mouse | Donkey | Alexa Fluor 647 | 1:1000 |

##### *In vitro* biosynthesis evaluation of GlcNAc analogs

E. coli recombinant inorganic pyrophosphatase PmPpA from *Pasteurella multocida* and NahK from *B. longum* were purchased from Chemily Glycoscience (Peachtree Corners, USA). Recombinant WT-AGX1 and variants have been expressed before.[33]

Analytical scale (12 µL) one-pot multienzyme (OPME) reactions with NahK and AGX1 WT and AGX1$^{F383A}$ were performed to assess if the chemically modified GlcNAc analogs are substrates for the biosynthetic enzymes. Each reaction mixture contained the corresponding GlcNAc analog (GlcNAc as a control and compounds **14, 15 and 16**) (2.5 mM), ATP (5 mM), UTP (5 mM), $MgCl_2$ (5 mM), BSA (1 mg/mL), kinase NahK (2.5 µg), PmPpA (0.045 U) and either AGX1$^{WT}$ (500 nM) or AGX1$^{F383A}$ (500 nM) in 100 mM Tris-HCl pH 8. Blanks with the above reaction mixture and without enzymes were included in each set of experiments to account for potential noise signal at products' retention time. Reactions were run for 16 h at 37 °C. After 16 h, reactions were stopped by adding equal volume of ice-cold acetonitrile and further cooled on ice for 30 min. The cooled reaction mixtures were centrifugated at 16,200 x g for 30 min at 4 °C to remove the precipitated enzymes. The supernatants were analyzed on UPLC-MS equipped with ACQUITY UPLC BEH Glycan 1.7 µm 2.1 x 50 mm column and gradient of 90-55% buffer B over 17 min at flow rate of 0.35 mL/min and column temperature at 50 °C; buffer A: 10 mM ammonium formate pH 4.5, buffer B: 10 mM ammonium formate in 90:10 v/v acetonitrile: water). Product formation was monitored by absorption at 260 nm and further confirmed by mass detection in negative mode. Turnover was assessed by the presence or absence of product peak.

##### Cell surface glycoprotein labelling

Cells stably transfected with pSBbi plasmids were seeded at a density of 250,000 cells/mL (K562 cells) or 500,000 cells/mL (Lec1 cells and Huh7 cells) into 6-well plates in their respective growth media without hygromycin. Cells were treated with DMSO, Ts-$Ac_3$GlcNButAz **17**, Ts- $Ac_3$GlcNPrAzMe(*S*) **18**, $Ac_4$GlcNPrAzMe(*S*) **13**, Ts-$Ac_3$GlcNAlk **19** or $Ac_4$GlcNAlk at the indicated concentrations. Cells were grown for 16 h. For in-gel fluorescence, cells were harvested in a V-shaped 96 well plate and washed twice with 0.2 mL 2% FBS in PBS (Resuspension Buffer). Cells were resuspended in 35 µL Resuspension Buffer and treated with 200 µM $CuSO_4$, 1 mM BTTAA, 10 mM sodium ascorbate, 10 mM ammonium guanidine chloride and 200 µM CF680 Alkyne or 200 µM CF680 picolyl Azide (Biotium, Fremont, USA) for 7 min at room temperature. The click reaction was quenched with equal volume of 3 mM bathocuproinedisulfonic acid in PBS. Cells were centrifuged, washed twice with Resuspension Buffer and once with PBS. Cells were lysed with 0.1 mL Lysis Buffer containing 50 mM Tris-HCl pH 8, 150 mM NaCl, 1% (v/v) Triton X-100, 0.5% (v/v) sodium deoxylcholate, 0.1% (w/v) SDS, 1 mM $MgCl_2$, and 100 mU/µL benzonase (Merck, Darmstadt, Germany) containing cOmplete protease inhibitors for 20 min at 4 °C on an orbital shaker. After centrifugation, supernatant containing labelled glycoproteins was transferred to a new plate and bicinchoninic acid assay was used to measure protein concentration. Reaction mixture was separated on a 4-20% Criterion™ gel (Bio-Rad, Hercules, USA) by SDS-PAGE and imaged on an Odyssey CLx imager (LI-COR Biosciences, Lincoln, USA). Total protein was stained with Coomassie. Protein expression was assessed by Western blot using mouse anti-myc antibody (Sigma-Aldrich, Gillingham, UK, Cat#05-724) diluted 1:1000 in Intercept TBS blocking buffer (LI-COR Biosciences, Lincoln, USA, Cat#927-60001), rabbit anti-FLAG antibody (Invitrogen, Carlsbad, USA, Cat#PA1-984B) diluted 1:1000 in Intercept TBS blocking buffer, and mouse anti-HA antibody (Abcam, Cambridge, UK, Cat#ab18181) diluted 1:1000 in Intercept TBS blocking buffer, followed by incubation with IRdye CW800 donkey anti-mouse (LI-COR Biosciences, Lincoln, USA, Cat#926-32212) or IRdye CW800 donkey anti-rabbit IgG secondary antibody (LI-COR Biosciences, Lincoln, USA, Cat#926-32213,) diluted 1:10,000 in Intercept TBS blocking buffer.

##### Chemical synthesis

Solvents and reagents were of synthesis and HPLC grade from Sigma Aldrich unless specified otherwise. Anhydrous conditions under inert gas were used unless specified otherwise. Thin layer chromatography was performed on DC-Fertigfolie Polygram SIL G/UV254 pre-coated with silica polyester sheets (0.2 mm thickness) (Macherey-Nagel, Düren, Germany). Spots were developed with sugar stain (0.1% (v/v) 3-methoxyphenol, 2.5% (v/v) sulfuric acid in EtOH) dipping solution. Solvents were removed under reduced pressure using a rotary evaporator and high vacuum. Medium pressure chromatography was performed on an Isolera system (Biotage, Uppsala, Sweden).

$^1$H and $^{13}$C NMR spectra were measured with Bruker Avance-400, 600 or 700 MHz spectrometers at 298 K. Chemical shifts (σ) are reported in parts per million (ppm) relative to the respective residual solvent peaks ($CDCl_3$: σ 7.26 in $^1H$ and 77.16 in $^{13}C$ NMR; $CD_3OH$: σ 3.31 in $^1H$ and 49.00 in $^{13}C$ NMR; $D_2O$: σ 4.79 in $^1H$). The following abbreviations are used to indicate peak multiplicities: s singlet; d doublet; dd doublet of doublets; dt doublet of triplets; m multiplet. Coupling constants (*J*) are reported in Hertz (Hz). Low resolution mass spectrometry spectrometry by electrospray ionization (ESI-LRMS) was performed on an Acquity UPLC-MS (Waters, Milford, USA) equipped with ACQUITY UPLC® BEH C18 column.

Compounds **3-10** were made previously.[2,34] Compounds **1** and **2** (UDP-GlcNAc and UDP-GlcNAz) are commercially available from Sigma-Aldrich (Gillingham, UK) and Chemily Glycoscience (Peachtree Corners, USA) respectively.

#### Compound characterization

##### (*S*)-3-azidobutyric acid (SI-11)

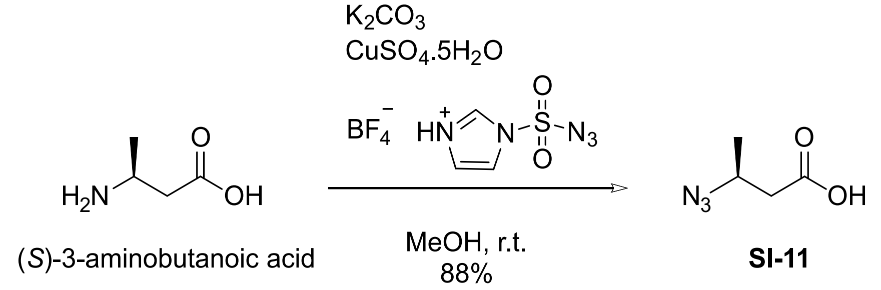


**SI-11**

(*S*)-3-azidobutyric acid was prepared by modification of an established procedure.[35] (*S*)-3-aminobutyric acid (Sigma Aldrich) (257.8 mg, 2.5 mmol), potassium carbonate (518 g, 3.75 mmol, 1.5 eq.) and copper (II) sulfate pentahydrate (5 mg, 20 µmol) were dissolved in methanol (10 mL). Imidazole-sufonyl azide tetra-fluoroborate (783 mg, 3 mmol, 1.2 eq.) was added and the reaction mixture was stirred at r.t. for 6 h. The solvent was reduced under reduced pressure and the aqueous slurry was diluted with 0.25 M phosphate buffer pH 6.2 (100 mL) and washed with EtOAc to remove the sulfonamide by-product. The aqueous phase was acidified with conc. HCl to pH 2 and the product extracted with EtOAc. The organic layer was dried over $MgSO_4$ and concentrated, but not fully dried, to provide the titled **SI-11** in solution of 1:1.75 product:EtOAc and 90% purity as determined by NMR, 284 mg, 2.2 mmol, 88 %. The crude product was taken forward without further purification.

##### 2-[(*S)*-3-Azidobutanamido]-2-deoxy- D-glucopyranose (11a)

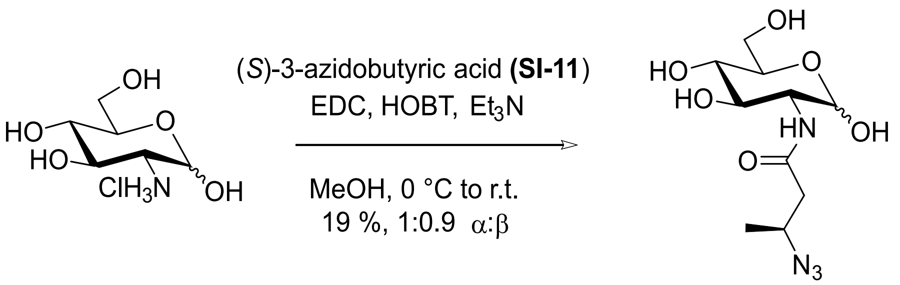


**11a**

To a stirred solution of (S)-3-azidobutyric acid, **SI-11** (168 mg, 1.3 mmol, 1.4 eq.) in anhydrous methanol (10 mL) were added D-glucosamine hydrochloride (200 mg, 0.93 mmol, 1 eq.) and $Et_3N$ (335 µl, 2.4 mmol, 2.78 eq.). The solution was cooled to 0 °C and EDC (173 mg, 1.1 mmol, 1.2 eq.) and HOBT (149 mg, 1.1 mmol, 1.2 eq.) were added. The reaction was warmed to r.t. and stirred for 16 h. On completion, the solvent was evaporated and the crude mixture purified on medium pressure flash chromatography with pre-packed silica column (Sfär Silica D Duo 25 g cartridge; A: DCM, B: methanol; 20 CV linear gradient from 0% to 60% B) to give the amide as yellow solid that still contained $Et_3N$ and HOBT. The mixture was further purified on Agilent 1260 Infinity II MDAP system equipped with C18 (100Å, 5 µm, 21.2 mm x 50 mm) column and buffers: A: water containing 0.1% formic acid; B: acetonitrile containing 0.1% formic acid. Fractions containing pure product were collected and lyophilized to give **11a** (51.8 mg, 0.18 mmol, 19 %, 1:0.8 α:β). $^1H$ NMR (400 MHz, $D_2O$) δ 5.22 (d, *J* = 3.5 Hz, 0.55H), 4.73 (d, *J* = 8.4 Hz, 0.45H), 4.03–3.67 (m, 5H), 3.60–3.41 (m, 2H), 2.59–2.44 (m, 2H), 1.31 (dd, *J* = 6.6, 1.2 Hz, 3H); $^{13}C$ NMR (100 MHz, $D_2O$) δ 173.6, 94.9, 90.7, 75.9, 73.7, 71.5, 70.5, 70.0 (d, *J* = 20.6 Hz), 60.6 (d, *J* = 15.6 Hz), 56.6, 55.0,

54.1, 42.2 (d, *J* = 38.9 Hz), 18.5. MS (ESI) calcd. for $C_{10}H_{18}N_4O_6$ (M-H$^+$) 289.12 found 289.00 *m/z*.

**(2S,3R,4R,5S,6R)-6-(acetoxymethyl)-3-(4-azidobutanamido)tetrahydro-2H-pyran-2,4,5-triyl triacetate (12)**

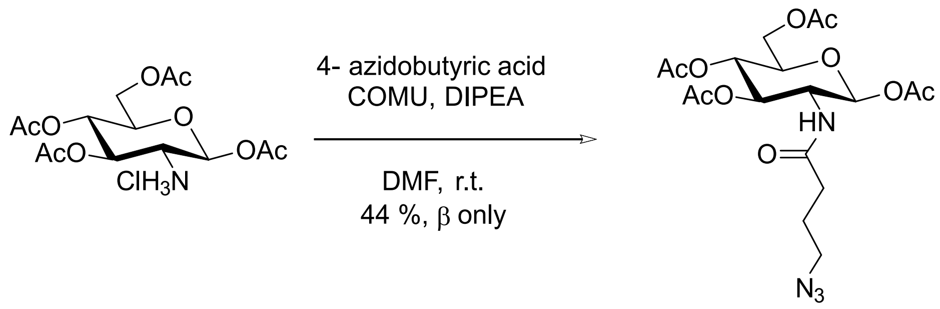


**SI-1** **12**

Tetraacetate glucosamine hydrochloride **SI-1** was prepared by an established procedure.[36] To a stirred solution of 4-azidobutyric acid (40 mg, 0.31 mmol, 1.2 eq.) in DMF (3 mL) were added tetraacetate **SI-1** (100 mg, 0.26 mmol, 1 eq.), DIPEA (136 µL, 0.78 mmol, 3 eq.) and COMU (134 mg, 0.31 mmol, 1.2 eq.). The solution was left to stir at room temperature for 16 h. After completion, the reaction mixture was diluted with EtOAc and washed with 0.1M HCl, sat. $NaHCO_3$ and brine. The organic layer was dried, concentrated, and purified on medium pressure flash chromatography and pre-packed silica column Biotage Sfär Silica D Duo 10 g (cyclohexane: EtOAc; 30-70% EtOAc over 5 CV) to give **12** (52.4 mg, 0.11 mmol, 44 %, β-anomer only) as a white solid. $^1$H NMR (400 MHz, $CDCl_3$) δ 5.73 (d, *J* = 8.8 Hz, 1H), 5.23–5.04 (m, 2H), 4.33–4.20 (m, 2H), 4.13 (dd, *J* = 12.5, 2.3 Hz, 1H), 3.80 (ddd, *J* = 9.6, 4.6, 2.1 Hz, 1H), 3.31 (t, *J* = 6.5 Hz, 2H), 2.21 (t, *J* = 7.2 Hz, 2H), 2.11 (s, 3H), 2.09 (s, 3H), 2.07–2.02 (m, 6H), 1.86 (p, *J* = 6.9 Hz, 2H); $^{13}$C NMR (100 MHz, $CDCl_3$) δ 171.9, 171.3, 170.8, 169.6, 169.4, 92.7, 73.1, 72.6, 67.8, 61.8, 53.4, 50.6, 33.2, 29.9, 24.6, 21.0, 20.9, 20.8. MS (ESI) calcd. for $C_{16}H_{23}N_4O_8^+$ (oxocarbenium ion) 399.15 found 399.08 *m/z*.

**(2S,3R,4R,5S,6R)-6-(acetoxymethyl)-3-((S)-3-azidobutanamido)tetrahydro-2H-pyran-2,4,5-triyl triacetate (13)**

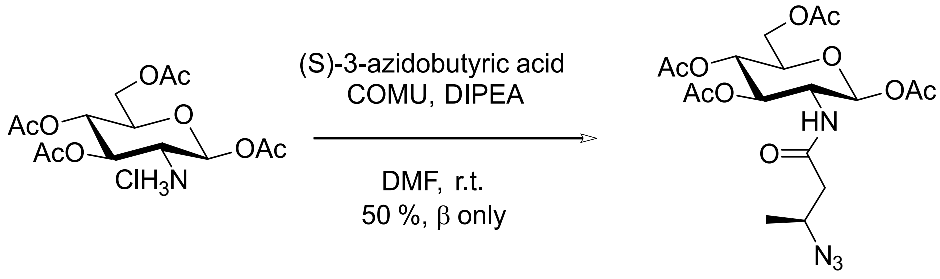


**SI-1** **13**

Tetraacetate glucosamine hydrochloride **SI-1** was prepared by an established procedure.[36] To a stirred solution of (S)-3-azidobutyric acid **11a** (40 mg, 0.31 mmol, 1.2 eq.) in DMF (3 mL) were added tetraacetate **SI-1** (100 mg, 0.26 mmol, 1 eq.), DIPEA (136 µL, 0.78 mmol, 3 eq.) and COMU (134 mg, 0.31 mmol, 1.2 eq.). The solution was left to stir at room temperature for 16 h. After completion, the reaction mixture was diluted with EtOAc and washed with 0.1M HCl, sat. $NaHCO_3$ and brine. The organic layer was dried, concentrated, and purified on medium pressure flash chromatography and pre-packed silica column Biotage Sfär Silica D Duo 10 g (cyclohexane: EtOAc; 10-70% EtOAc over 10 CV) to give **13** (58.5 mg, 0.13 mmol, 50 %, β-anomer only) as a white solid. $^1$H NMR (400 MHz, $CDCl_3$) δ 5.71 (d, *J* = 8.8 Hz, 1H), 5.26 – 5.05 (m, 2H), 4.36–4.23 (m, 2H), 4.12 (dd, *J* = 12.6, 2.3 Hz,

1H), 4.01–3.92 (m, 1H), 3.84 (ddt, *J* = 9.4, 4.0, 2.1 Hz, 1H), 2.27 (dd, *J* = 14.6, 4.6 Hz, 1H), 2.18 (dd, *J* = 14.6, 8.7 Hz, 1H), 2.10 (d, *J* = 1.2 Hz, 3H), 2.08 (d, *J* = 1.5 Hz, 3H), 2.06 (s, 3H), 2.05–2.02 (m, 3H), 1.29 (dd, *J* = 6.6, 1.2 Hz, 3H); $^{13}$C NMR (100 MHz, $CDCl_3$) δ 171.4, 170.7, 169.8, 169.4, 169.3, 92.4, 72.9, 72.4, 67.9, 61.7, 54.5, 52.8, 43.1, 20.9, 20.7, 20.6 (d, *J* = 1.9 Hz), 19.2. MS (ESI) calcd. for $C_{16}H_{23}N_4O_8^+$ (oxocarbenium ion) 399.15 found 399.05 *m/z*.

**Chemoenzymatic synthesis of UDP-GlcNPrAzMe(*S*) (11)**

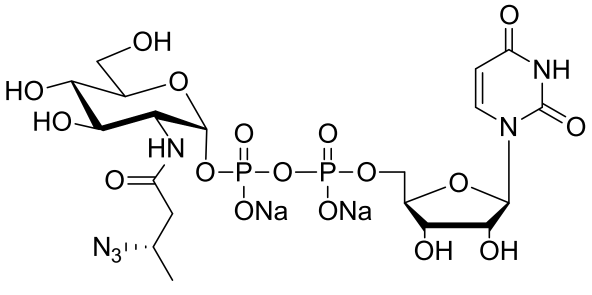


UDP-GlcNPrAz3Me(S) **11**

CIAP (Calf Intestine Alkaline Phosphatase) was purchased from Thermo Fisher Scientific (Waltham, USA).

Analytical scale one-pot multienzyme (OPME) reactions with NahK, AGX1 variants and compound **11a** were performed to establish the optimal system for the generation of preparative scale UDP-sugar analog **11**, following established procedure.[37]

Compound **11** was prepared using NahK and AGX1$^{F383A}$, according to previously established method.[4] The reaction mixture contained **11a** (2.5 mM), ATP (5 mM), UTP (5 mM), $MgCl_2$ (5 mM), BSA (1 mg/mL), kinase NahK (2.5 µg), PmPpA (0.045 U) and AGX1 F383A (500 nM) in 100 mM Tris-HCl pH 8 and total volume of 15 mL. After 16 h, CIAP (12.5 U) was added and the mixture left to incubate for further 16 h, after which the reaction was quenched by the addition of an equal volume (15 mL) of ice-cold ethanol. The mixture was left on ice for 30 min, centrifugated and the supernatant collected. The ethanol from the supernatant was removed under vacuum and the remaining aqueous solution dried by lyophilisation. The dried product was desalted by passing it through a pre-equilibrated 10 g C18 Sep Pak column (Waters, Milford, USA) and product eluted with 20% acetonitrile in water. Fractions containing the product were collected, dried on a freeze-drier and further purified on Agilent 1260 Infinity II MDAP system (Agilent Technologies, UK) equipped with HILIC XBridge® BEH Amide OBD™ Prep (130Å, 5 µm, 10 mm x 100 mm) column. Buffers were: A: 10 mM ammonium formate at pH 4.5; B: 10 mM ammonium formate in 90:10 (v/v) acetonitrile: water; flow rate: 19 mL/min. Gradient was from 90% to 65% Buffer B over 10 min. The lyophilized final product was passed through a short (4 g resin) ion exchange column of Dowex 50W X8 $Na^+$ form and concentrated to give **11** as white solid, 16.5 mg, 14 µmol, 65 % yield.

**UDP-GlcNPrAzMe(*S*) (11)** white solid, 65%; $^1$H NMR (400 MHz, $D_2O$) δ 7.88 (d, *J* = 8.1 Hz, 1H), 5.96–5.84 (m, 2H), 5.44 (dd, *J* = 7.2, 3.3 Hz, 1H), 4.32–4.26 (m, 2H), 4.15 (dddt, *J* = 24.7, 14.9, 8.7, 3.0 Hz, 3H), 3.97–3.69 (m, 6H), 3.47 (dd, *J* = 10.1, 9.1 Hz, 1H), 2.54–2.42 (m, 2H), 1.22 (d, *J* = 6.6 Hz, 3H); $^{13}$C NMR (100 MHz, $D_2O$) δ 173.7, 166.3, 151.8, 141.7, 102.7, 94.5, 88.5, 83.2, 83.2, 73.8, 73.0, 70.8, 69.6, 64.9, 60.3, 55.0, 53.7, 42.0, 18.6. MS (ESI) calcd. for $C_{19}H_{28}N_6O_{17}P_2$ (M-H$^+$) 675.11 found 675.21 *m/z*.

**2-(4-Azidobutanamido)-2-deoxy-6-O-toluenesulfonyl-D-glucopyranose (14)**

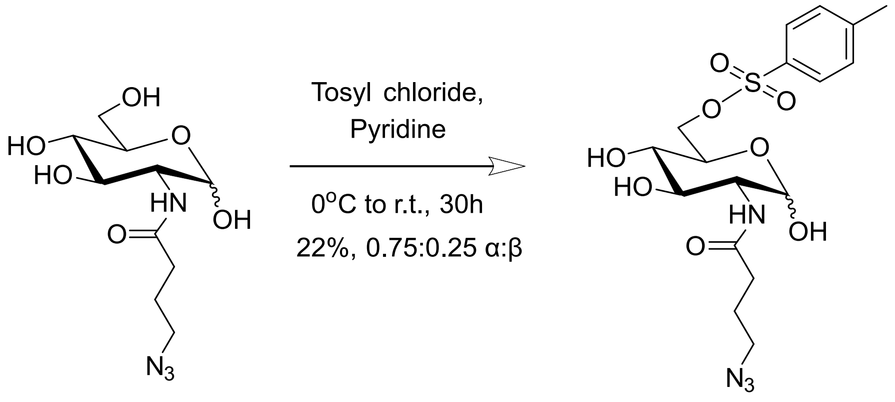


**GlcNButAz** **14**

GlcNButAz prepared according to a previously reported method[2] (48 mg, 0.164 mmol, 1 eq) was dissolved in pyridine (2.5 mL) and cooled to 0°C. P-toluene sulfonyl chloride (81 mg, 0.425 mmol, 1.3 eq) was added and the reaction was stirred under nitrogen and left to warm up to r.t. for a further 30 h. On completion, the reaction was quenched by the addition of methanol (MeOH, 0.625 mL) and the solvent removed under vacuum. The dried residue was dissolved in a minimal amount of 20% MeOH in water (0.5 mL) and purified on medium pressure flash chromatography C18 column (Sfär C18 12 g cartridge; Buffer A: $H_2O$ with 0.1% FA, Buffer B: ACN with 0.1% FA; linear gradient from 0% to 100% Buffer B over 13 CV), to give **14** (15.9 mg, 0.036 mmol, 22%, 0.75:0.25 α:β) as a white powder. $^1$H NMR (400 MHz, $CD_3OD$) δ 7.76 – 7.65 (m, 2H), 7.37 – 7.25 (m, 2H), 4.89 (d, *J* = 3.4 Hz, 0.75H), 4.44 (d, J = 8.3 Hz, 0.25H), 4.20 (ddd, *J* = 24.7, 10.6, 2.0 Hz, 1H), 4.11 – 3.98 (m, 1H), 3.82 (ddd, *J* = 10.0, 5.4, 2.0 Hz, 1H), 3.67 (dd, *J* = 10.7, 3.5 Hz, 1H), 3.61 – 3.41 (m, 1H), 3.32 – 3.22 (m, 2H), 3.14 (td, J = 10.2, 8.7 Hz, 1H), 2.36 (s, 3H), 2.26 – 2.17 (m, 2H), 1.83 – 1.67 (m, 2H); $^{13}$C NMR (100 MHz, $CD_3OD$) δ 174.0, 145.1, 132.9, 129.7, 127.7, 91.1, 71.1, 70.7, 69.7, 69.3, 54.2, 50.4, 32.4, 24.8, 20.2. MS (ESI) calcd. for $C_{17}H_{23}N_4O_7S^+$ (oxonium ion) 427.13 found 427.27 *m/z*.

**2-[(*S*)-3-Azidobutanamido]-2-deoxy-6-O-toluenesulfonyl-D-glucopyranose (15)**

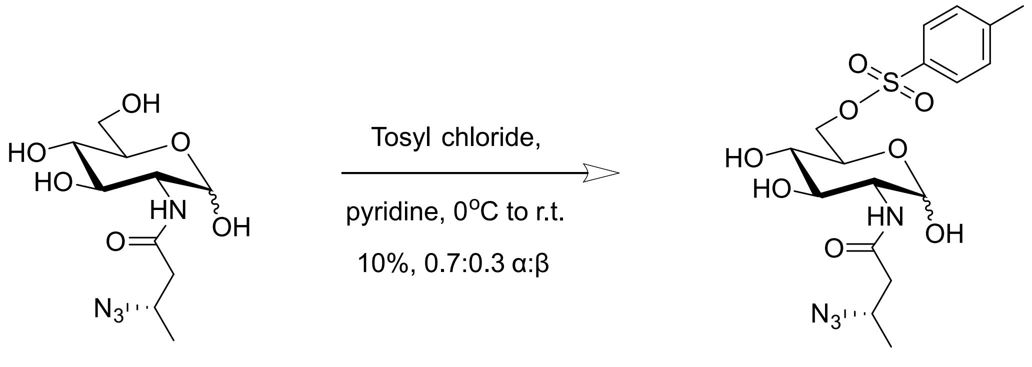


**GlcNPrAzMe(*S*)** **15**

GlcNPrAzMe(*S*) prepared according to a previously reported method [2] (55 mg, 0.276 mmol, 1 eq) was dissolved in pyridine (2.5 mL) and cooled to 0°C. P-toluene sulfonyl chloride (52 mg, 0.414 mmol, 1.5 eq) was added and the reaction was stirred under nitrogen for 24 h, after which additional p-toluene sulfonyl chloride (26 mg, 0.207 mmol, 0.75 eq) was added. After an additional 3 h, more p-toluene sulfonyl chloride (26 mg, 0.207 mmol, 0.75 eq) was added and the reaction stirred at 0°C for further 24 h. After that, the reaction was brought up to room temperature and left to stir for additional 24 h. Finally, additional p-toluene sulfonyl chloride (17 mg, 0.138 mmol, 0.5 eq) was added and left to react for further 3 h before quenching with MeOH (0.625 mL). On completion, the solvent was evaporated, the dried residue dissolved in a minimal amount of 20% MeOH in water (0.5 mL) and purified on medium pressure flash chromatography C18 column (Sfär C18 12 g cartridge; Buffer A: $H_2O$ with 0.1% FA, Buffer B: ACN with 0.1% FA; linear gradient from 0% to 60% Buffer B over 18 CV), to give **15** (11.7 mg, 0.026 mmol, 10%, 0.75:0.25 α:β) as a white powder. $^1$H NMR (400 MHz, $CD_3OD$) δ 7.74 – 7.65 (m, 2H), 7.34 (dtd, *J* = 7.3, 1.7, 0.7 Hz, 2H), 4.89 (t, *J* = 3.3 Hz, 0.75H), 4.44 (d, *J* = 8.3 Hz, 0.25H), 4.20 (ddd, *J* = 24.1, 10.6, 2.0 Hz, 1H), 4.05 (ddd, *J* = 19.8, 10.6, 5.8 Hz, 1H), 3.90 – 3.77 (m, 2H), 3.67 (dd, *J* = 10.7, 3.5 Hz, 1H), 3.61 – 3.49 (m, 1H), 3.20 – 3.08 (m, 1H), 2.40 – 2.18 (m, 5H), 1.18 (dd, *J* = 6.5, 4.3 Hz, 3H); $^{13}$C NMR (100 MHz, $CD_3OD$) δ 171.5, 145.0, 129.6 (d, *J* = 8.3 Hz), 127.7, 91.0, 71.0, 70.7, 69.7, 69.3, 54.8, 54.2, 41.9, 20.2, 18.1. MS (ESI) calcd. for $C_{17}H_{23}N_4O_7S^+$ (oxonium ion) 427.13 found 427.27.

**2-(4-Pentynamido)-2-deoxy-6-O-toluenesulfonyl-D-glucopyranose (16)**

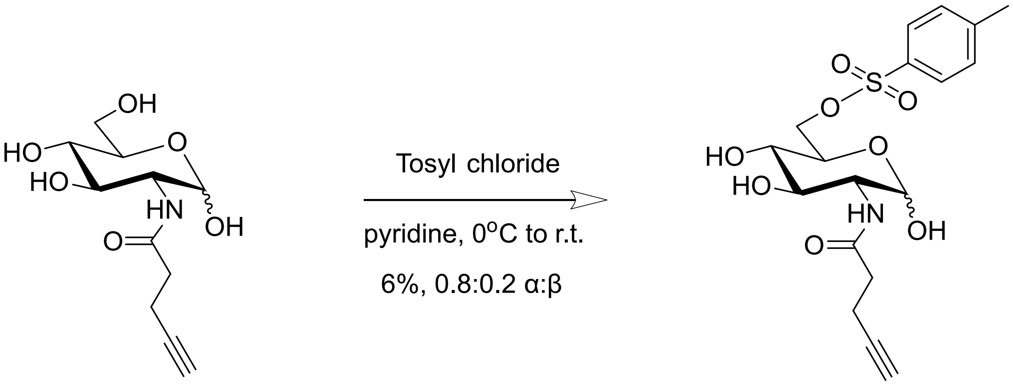


**GlcNAlk** **16**

GlcNAlk prepared according to a previously reported method [2] (150 mg, 0.579 mmol, 1 eq) was dissolved in pyridine (2.5 mL) and cooled to 0°C. P-toluene sulfonyl chloride (143 mg, 0.752 mmol, 1.3 eq) was added and the reaction was stirred under nitrogen for 7 h while warming up to room temperature. The reaction was quenched by the addition of MeOH (0.625 mL). On completion, the solvent was evaporated, the dried residue dissolved in a minimal amount of water (0.5 mL) and purified on medium pressure flash chromatography C18 column (Sfär C18 12 g cartridge; Buffer A: $H_2O$ with 0.1% FA, Buffer B: ACN with 0.1% FA; linear gradient from 0% to 60% Buffer B over 18 CV), to give **16** (13.8 mg, 0.033 mmol, 6%, 0.80:0.20 α:β) as a white powder. $^1$H NMR (400 MHz, $CD_3OD$) δ 7.75 – 7.65 (m, 2H), 7.37–7.27 (m, 2H), 4.88 (d, *J* = 3.4 Hz, 0.80H), 4.44 (d, *J* = 8.3 Hz, 0.22H), 4.20 (ddd, *J* = 24.2, 10.6, 2.0 Hz, 1H), 4.05 (ddd, *J* = 19.5, 10.6, 5.9 Hz, 1H), 3.83 (ddd, *J* = 10.0, 5.4, 2.0 Hz, 1H), 3.68 (dd, *J* = 10.6, 3.5 Hz, 1H), 3.54 (dd, *J* = 10.7, 8.7 Hz, 1H), 3.20–3.08 (m, 1H), 2.35 (d, *J* = 1.6 Hz, 7H), 2.18–2.09 (m, 1H); $^{13}$C NMR (100 MHz, $CD_3OD$) δ 172.9, 145.0, 133.0, 129.6, 127.7, 91.2, 82.2, 71.1, 70.6, 69.7, 69.3, 68.8, 54.2, 34.5, 20.2, 14.2. MS (ESI) calcd. for $C_{18}H_{22}NO_7S^+$ (oxonium ion) 396.11 found 396.26 *m/z*.

**2-(4-Azidobutanamido)-2-deoxy-1,3,4-tri-O-acetyl-6-O-toluenesulfonyl-D-glucopyranose (17)**

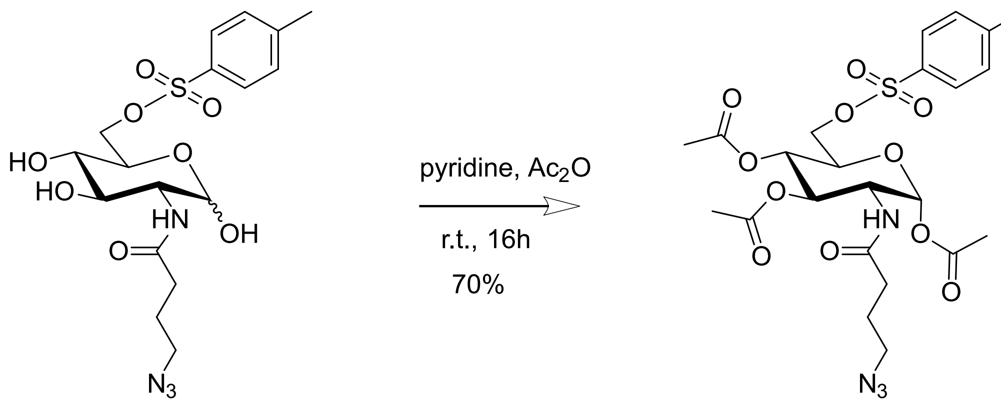

**14** **17**

To a stirred solution of Ts-GlcNButAz **14** (16 mg, 0.036 mmol, 1 eq) in pyridine (0.4 mL), acetic anhydride (0.014 mL, 0.148 mmol, 4.1 eq) was added slowly and the mixture was stirred for 30 min at 0 °C and then at room temperature overnight. After completion, the reaction slurry was diluted with EtOAc (50 mL) and washed with 0.1 M HCl (3 x 40 mL), saturated $NaHCO_3$ (3 x 40 mL) and brine (40 mL). The organic layer was then dried and concentrated to give **17** (14.5 mg, 0.025 mmol, 70%, α-anomer only) as a white powder with minor grease impurity. $^1$H NMR (400 MHz, $CDCl_3$) δ 7.74 – 7.64 (m, 2H), 7.32 – 7.24 (m, 2H), 6.02 (d, *J* = 3.7 Hz, 1H), 5.52 (d, *J* = 8.8 Hz, 1H), 5.13 (dd, *J* = 11.0, 9.5 Hz, 1H), 5.04 – 4.93 (m, 1H), 4.28 (ddd, *J* = 11.0, 8.8, 3.7 Hz, 1H), 4.02 – 3.96 (m, 2H), 3.30 – 3.15 (m, 2H), 2.39 (s, 3H), 2.12 (d, *J* = 7.9 Hz, 5H), 2.04 – 1.91 (m, 7H), 1.84 – 1.73 (m, 2H); $^{13}$C NMR (100 MHz, $CDCl_3$) δ 171.8, 169.2, 168.6, 145.3, 132.3, 129.9, 128.1, 90.0, 70.4, 69.6, 67.9, 67.4, 53.4, 51.0, 50.5, 32.9, 24.4, 21.7, 20.9, 20.7, 20.5. MS (ESI) calcd. for $C_{21}H_{27}N_4O_9S^+$ (oxonium ion) 511.15 found 511.11 *m/z*.

**2-[(*S*)-3-Azidobutanamido]-2-deoxy-1,3,4-tri-O-acetyl-6-O-toluenesulfonyl-D-glucopyranose (18)**

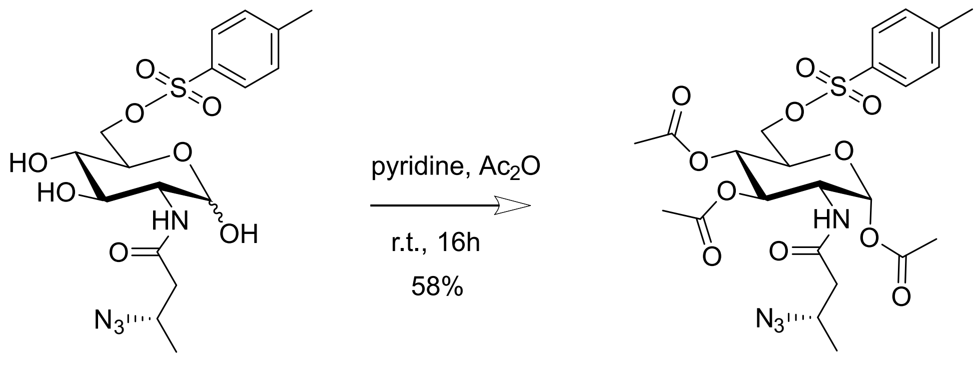

**15** **18**

To a stirred solution of Ts-GlcNPrAzMe(*S*) **15** (7.7 mg, 0.018 mmol, 1 eq) in pyridine (0.4 mL) acetic anhydride (0.007 mL, 0.079 mmol, 4.1 eq) was added slowly and the mixture was stirred for 30 min at 0 °C and then at room temperature overnight. After completion, the reaction slurry was diluted with EtOAc (50 mL) and washed with 0.1 M HCl (3 x 40 mL), saturated $NaHCO_3$ (3 x 40 mL) and brine (40 mL). The organic layer was then dried and concentrated to give **18** (6.0 mg, 0.011 mmol, 58%, α-anomer only) as a white powder with minor grease impurity. $^1$H NMR (400 MHz, $CDCl_3$) δ 7.74 – 7.65 (m, 2H), 7.33 – 7.25 (m, 2H), 6.01 (d, *J* = 3.6 Hz, 1H), 5.72 (d, *J* = 8.8 Hz, 1H), 5.16 (dd, *J* = 11.1, 9.4 Hz, 1H), 5.04 – 4.92 (m, 1H), 4.35 – 4.22 (m, 1H), 4.07 – 3.96 (m, 3H), 3.88 (dqd, *J* = 8.8, 6.6, 4.3 Hz, 1H),

2.39 (s, 3H), 2.21 (dd, $J$ = 14.7, 4.3 Hz, 1H), 2.13 – 2.09 (m, 4H), 1.98 (d, $J$ = 3.1 Hz, 6H), 1.21 (d, $J$ = 6.6 Hz, 3H). $^{13}$C NMR (150 MHz, $CDCl_3$) δ 171.9, 169.7, 169.3, 168.7, 145.4, 132.4, 130.0, 128.3, 90.2, 70.3, 69.8, 68.0, 67.5, 54.6, 51.1, 43.1, 21.8, 21.1, 20.8, 20.7, 19.3. MS (ESI) calcd. for $C_{21}H_{27}N_4O_9S^+$ (oxonium ion) 511.15 found 511.13 *m/z*.

**2-(4-Pentynamido)-2-deoxy-1,3,4-tri-O-acetyl-6-O-toluenesulfonyl-D-glucopyranose (19)**

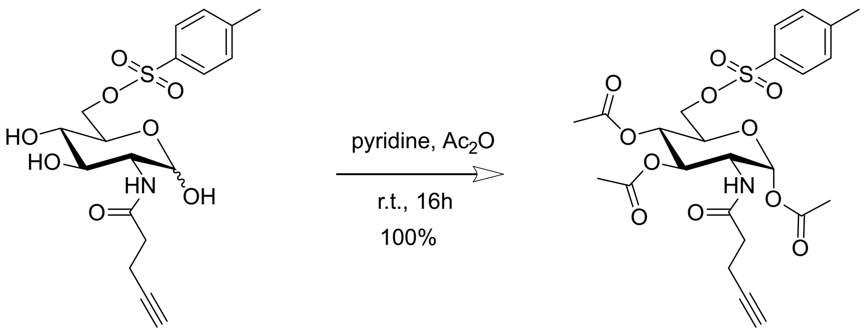


**16** **19**

To a stirred solution of Ts-GlcNAlk **16** (8 mg, 0.019 mmol, 1 eq) in pyridine (0.4 mL), acetic anhydride (0.007 mL, 0.079 mmol, 4.1 eq) was added slowly and the mixture was stirred for 30 min at 0 °C and then at room temperature overnight. After completion, the reaction slurry was diluted with EtOAc (50 mL) and washed with 0.1 M HCl (3 x 40 mL), saturated $NaHCO_3$ (3 x 40 mL) and brine (40 mL). The organic layer was then dried and concentrated to give **19** (10.3 mg, 0.019 mmol, 100%, α-anomer only) as a white powder. $^1$H NMR (400 MHz, $CDCl_3$) δ 7.74–7.62 (m, 2H), 7.30–7.22 (m, 2H), 6.01 (d, $J$ = 3.7 Hz, 1H), 5.67 (d, $J$ = 8.9 Hz, 1H), 5.15 (dd, $J$ = 11.1, 9.4 Hz, 1H), 5.09–4.91 (m, 1H), 4.31 (ddd, $J$ = 11.0, 8.9, 3.7 Hz, 1H), 4.11–3.93 (m, 2H), 2.38 (d, $J$ = 4.9 Hz, 5H), 2.32–2.20 (m, 2H), 2.11 (s, 2H), 2.00–1.92 (m, 6H), 1.30–1.14 (m, 3H); $^{13}$C NMR (100 MHz, $CDCl_3$) δ 171.6, 170.9, 168.5, 153.7, 145.2, 129.9, 128.2, 121.0, 90.1, 82.5, 70.3, 69.6, 68.0, 67.4, 50.9, 35.1, 29.7, 21.7, 20.9, 20.8, 20.5, 14.7. MS (ESI) calcd. for $C_{22}H_{26}NO_9S^+$ (oxonium ion) 480.13 found 480.11 *m/z*.

#### NMR Characterization

$^{1}$H NMR, $D_2O$, 400 MHz

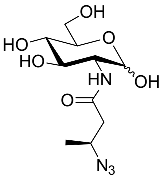


**11a**

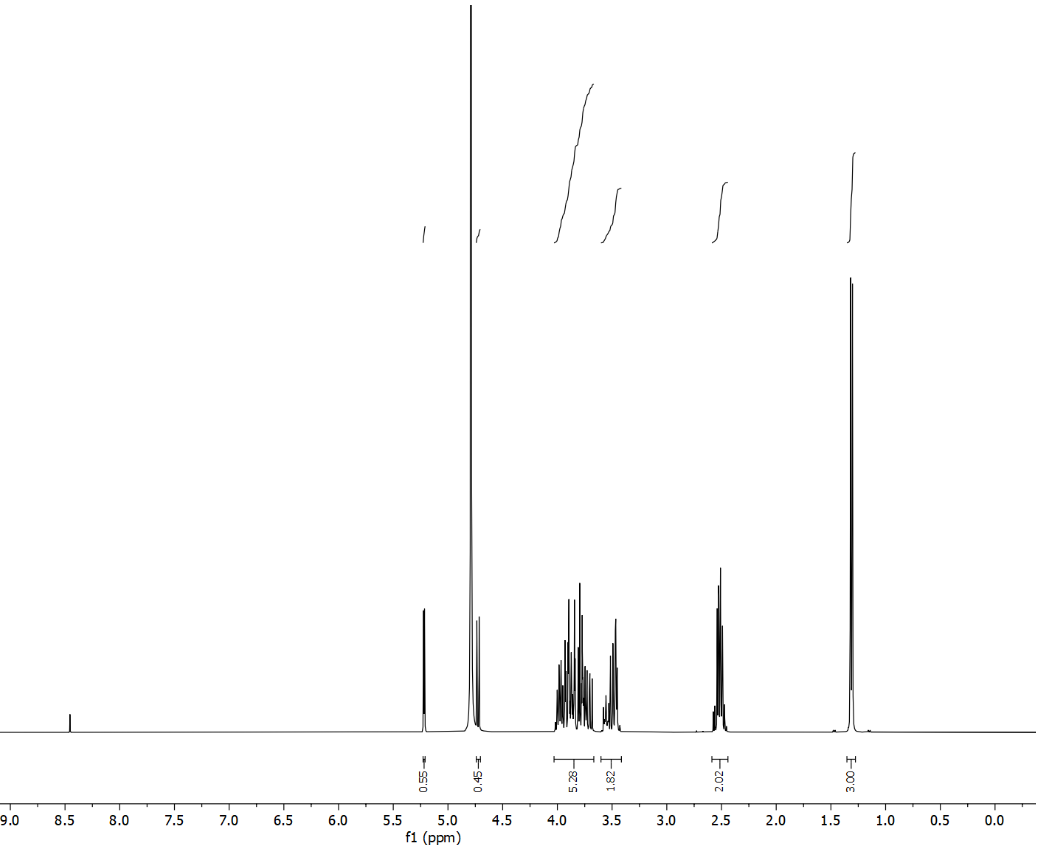


$^{13}$C NMR, $D_2O$, 100 MHz

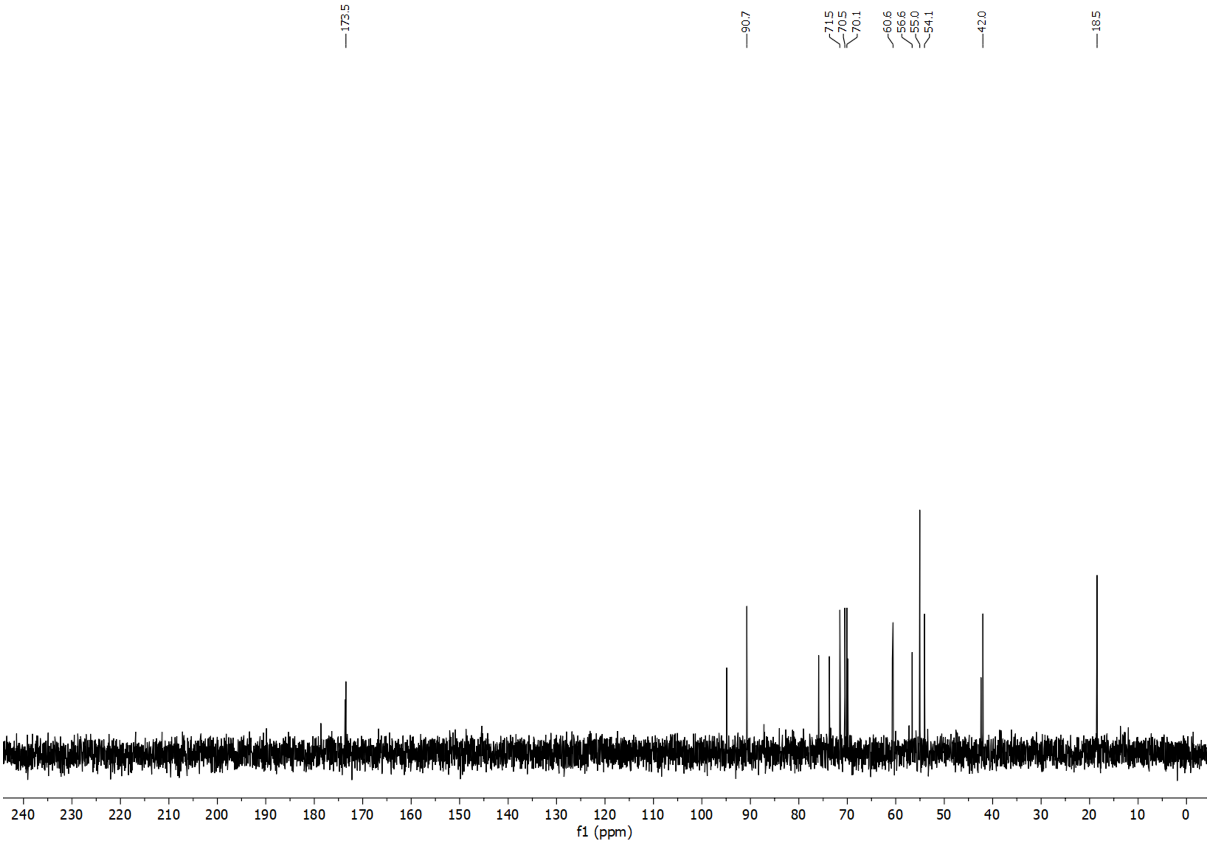

### [1]H NMR, $D_2O$, 400 MHz

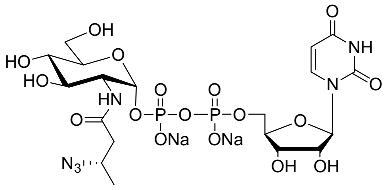


UDP-GlcNPrAz3Me(S) **(11)**

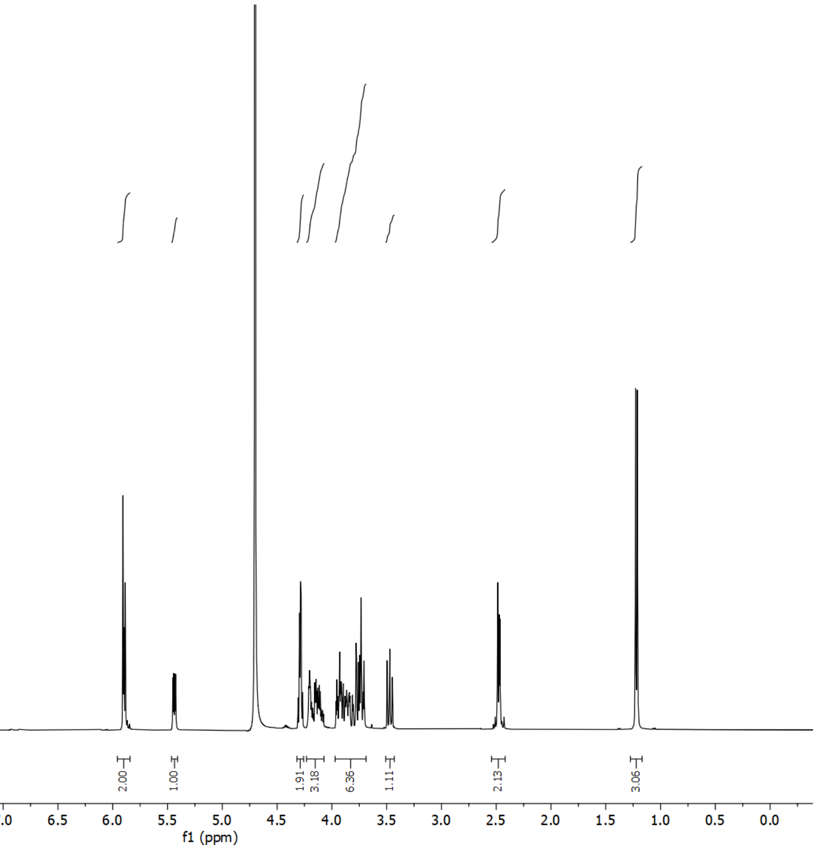


### [13]C NMR, $D_2O$, 100 MHz

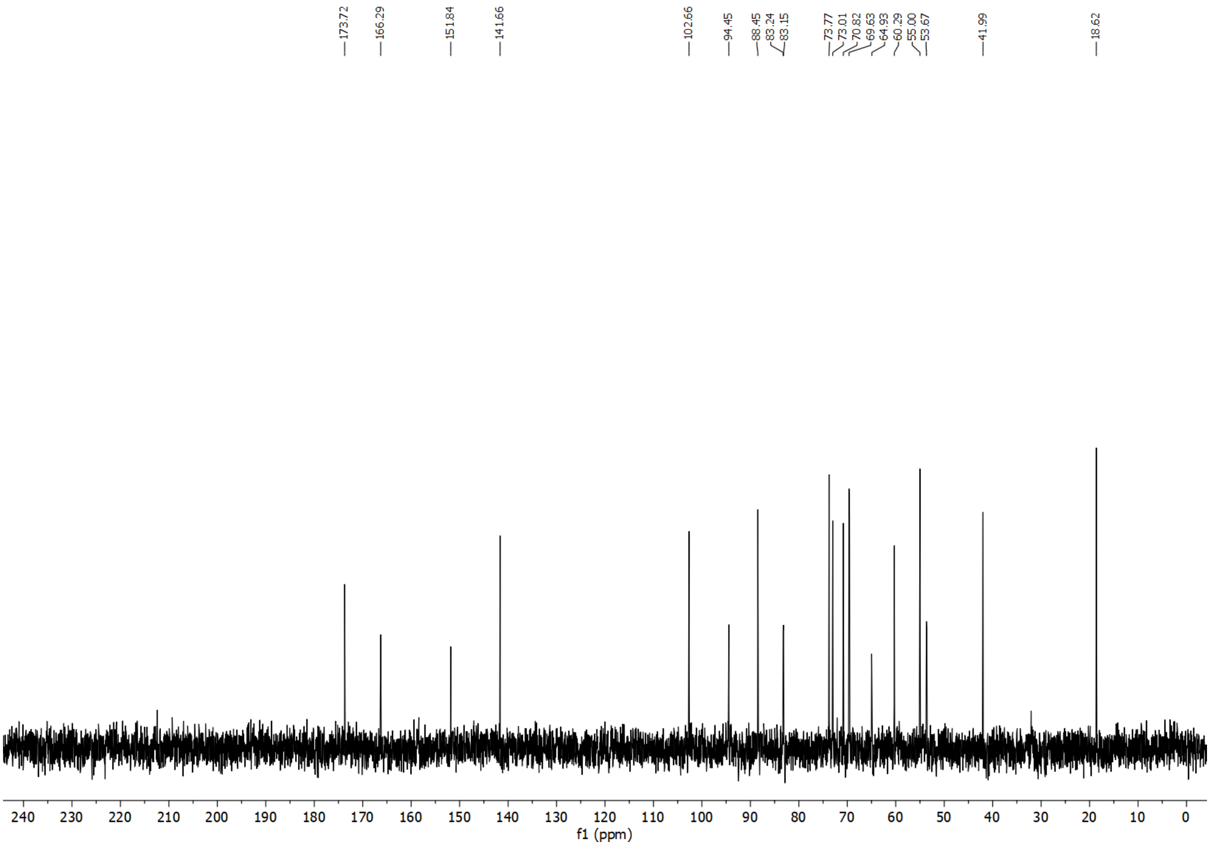

#### $^1H$ NMR, $CDCl_3$, 400 MHz

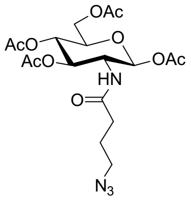


Ac$_4$GlcNButAz **(12)**

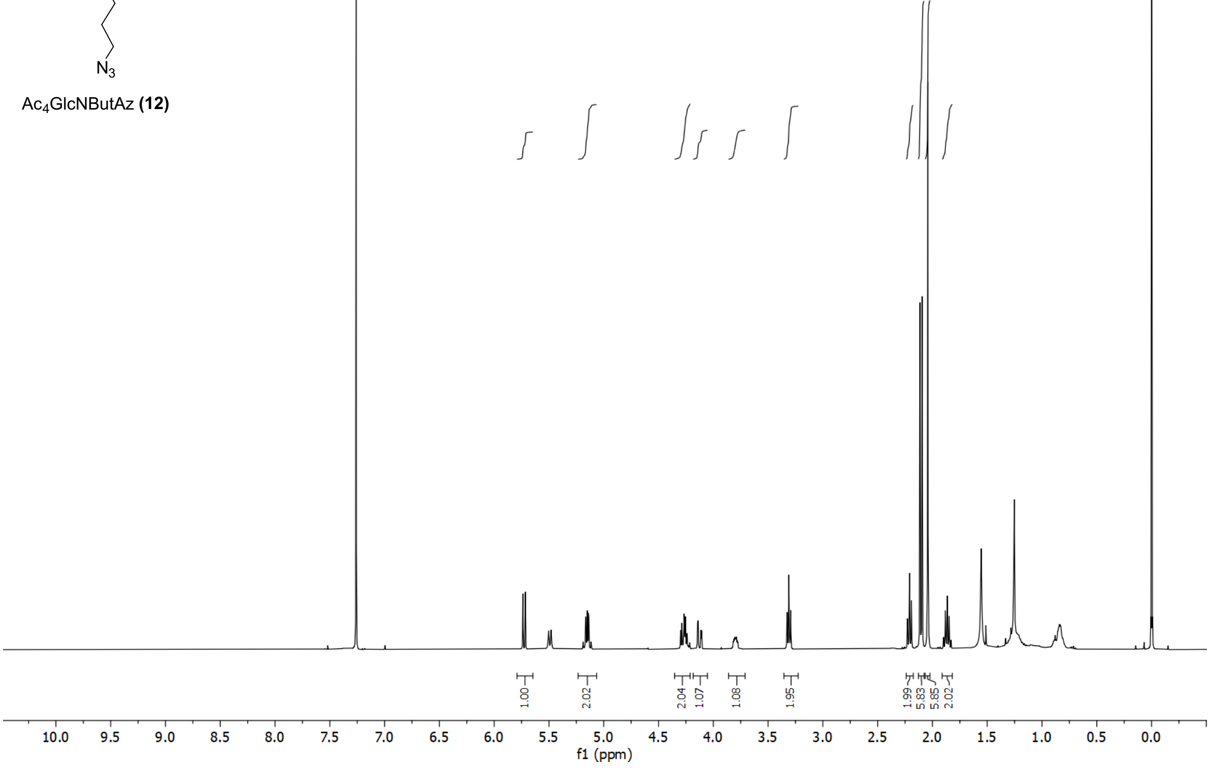


#### $^{13}C$ NMR, $CDCl_3$, 100 MHz

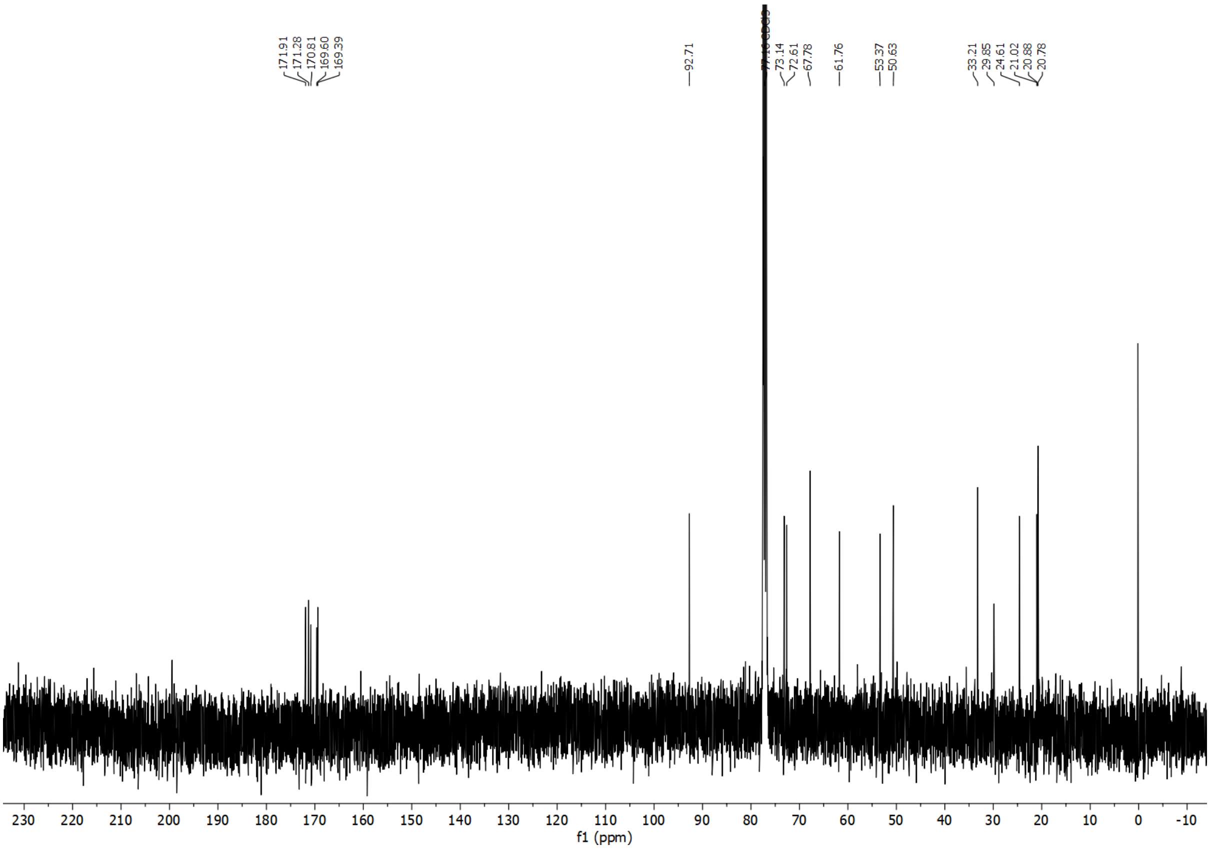

$^{1}$H NMR, $CDCl_3$, 400 MHz

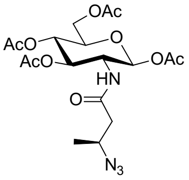


Ac$_4$GlcNPrAzMe(*S*) **(13)**

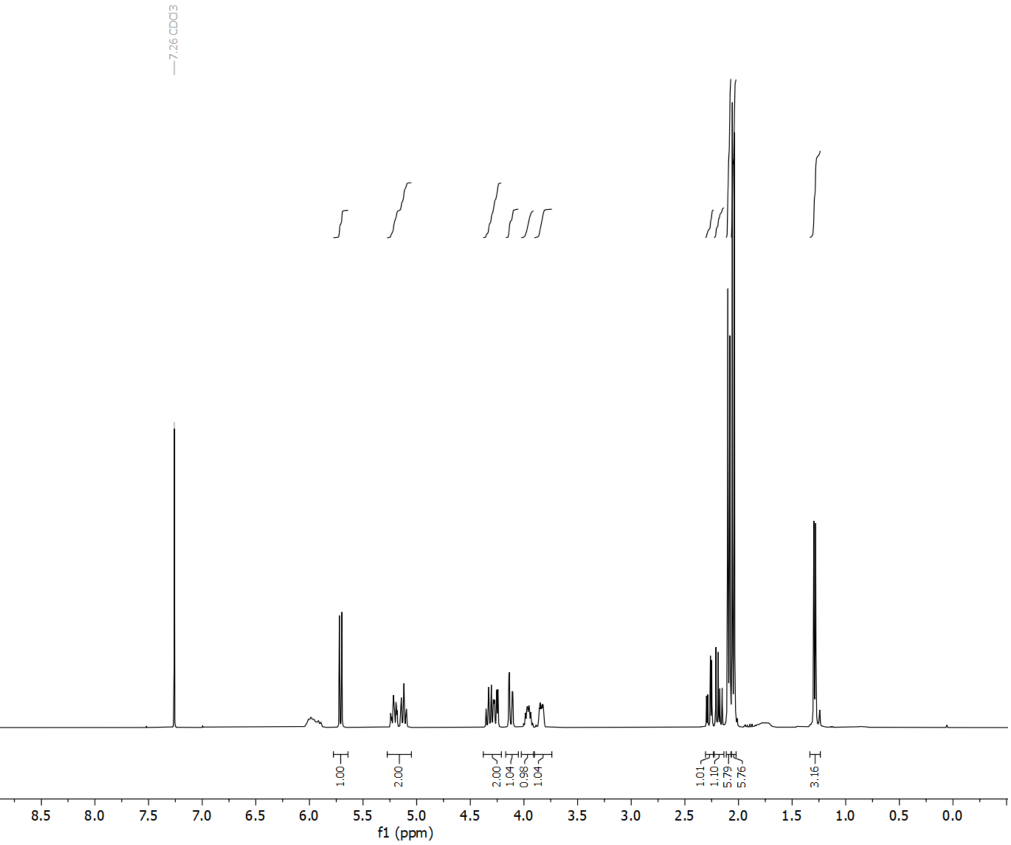


$^{13}$C NMR, $CDCl_3$, 100 MHz

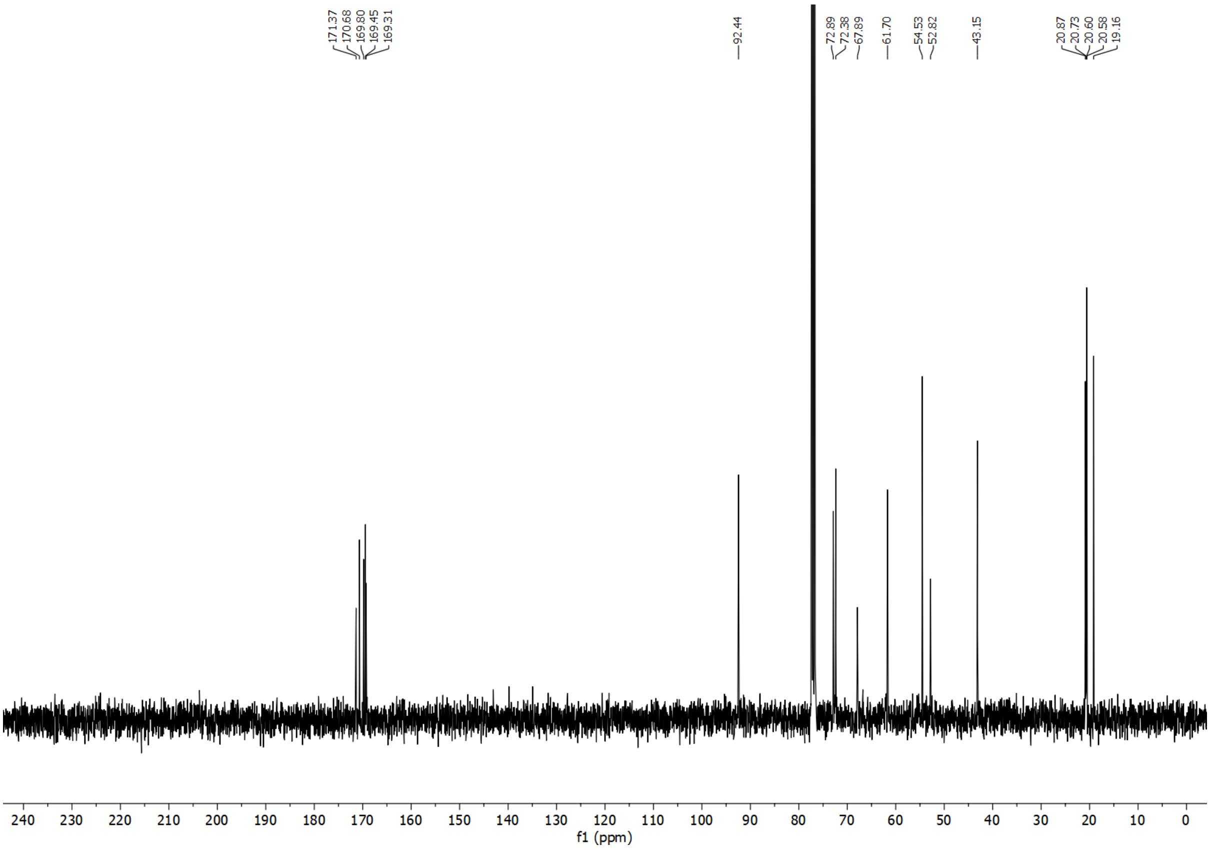

[1]H NMR, $CD_3OD$, 400 MHz

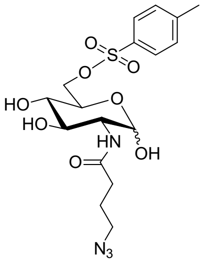


Ts-GlcNButAz (**14**)

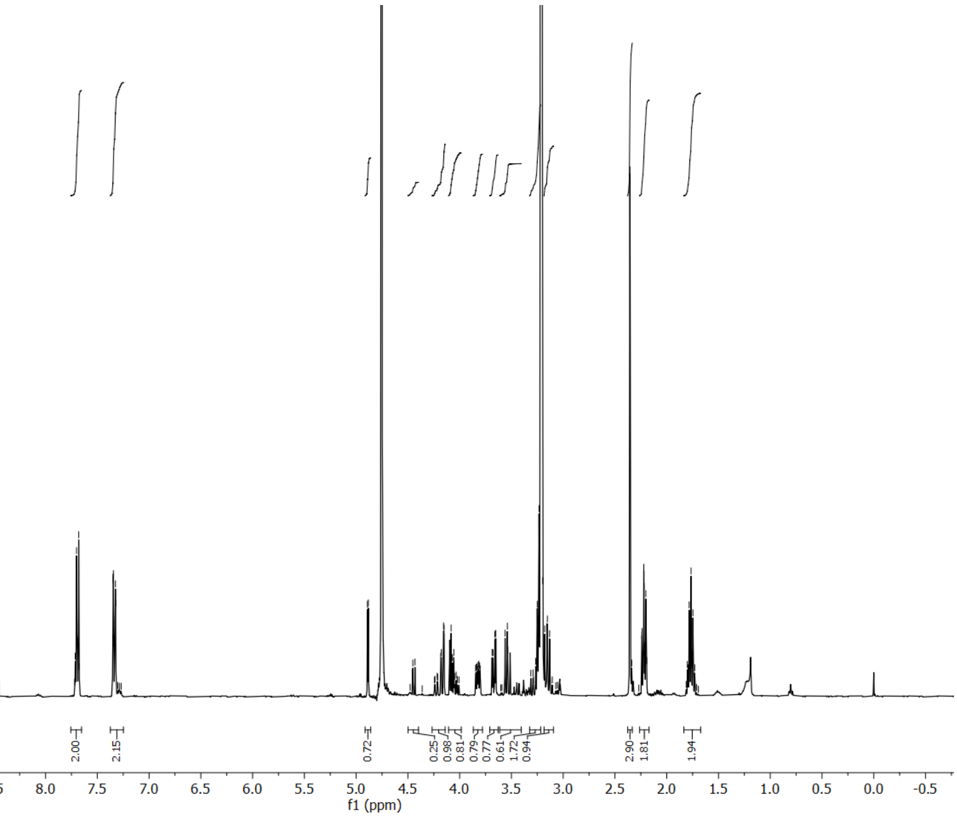


[13]C NMR, $CD_3OD$, 100 MHz

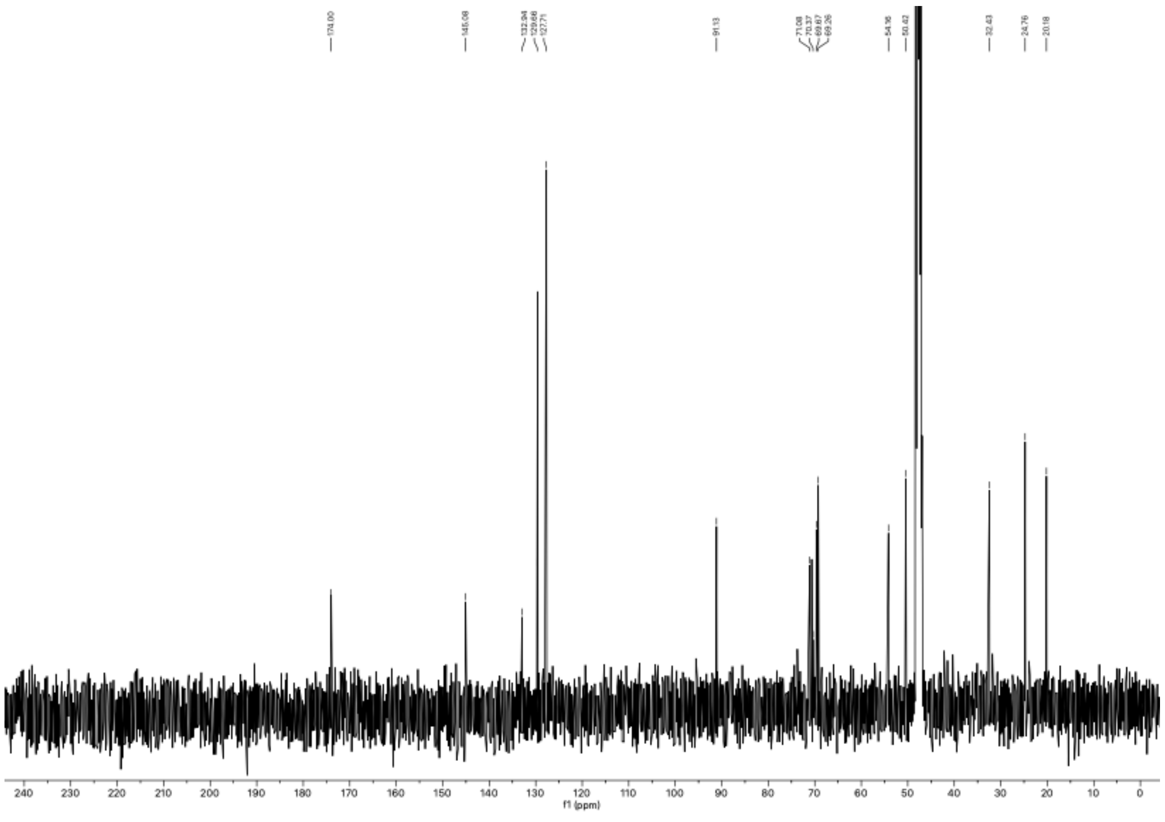

[1]H NMR, $CD_3OD$, 400 MHz

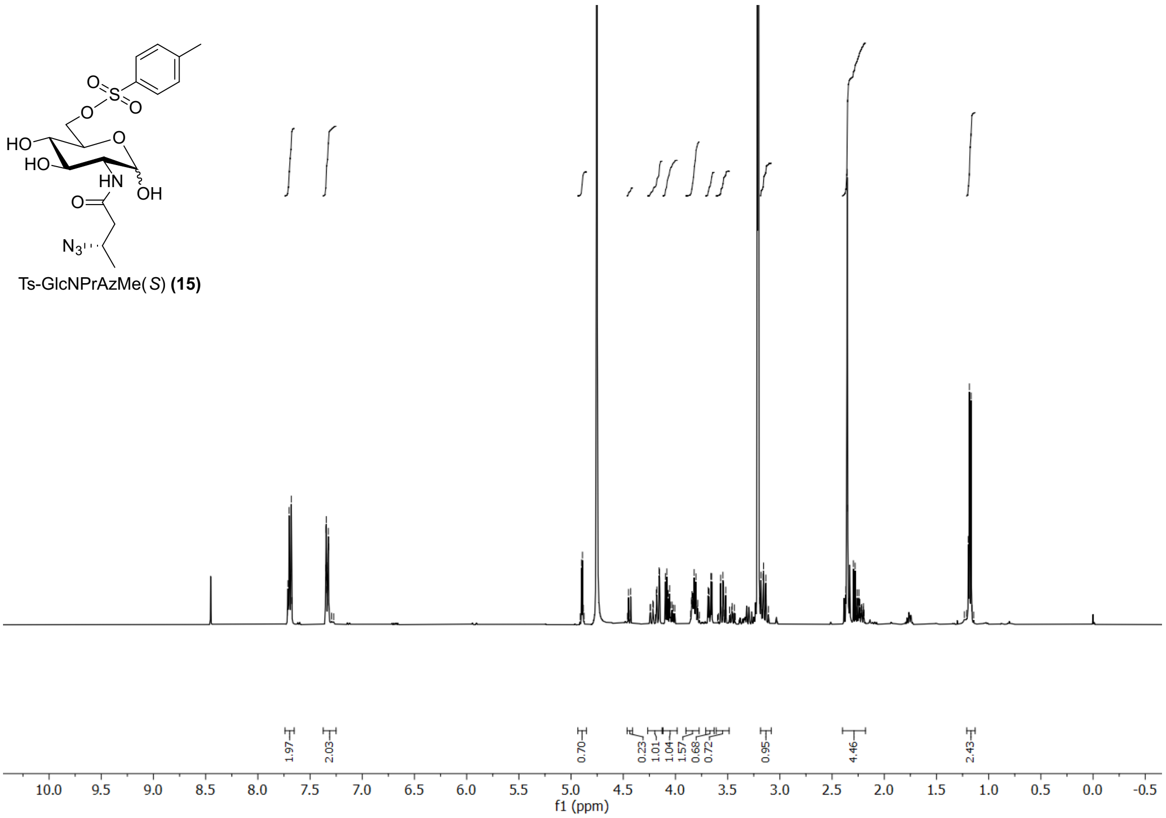


[13]C NMR, $CD_3OD$, 100 MHz

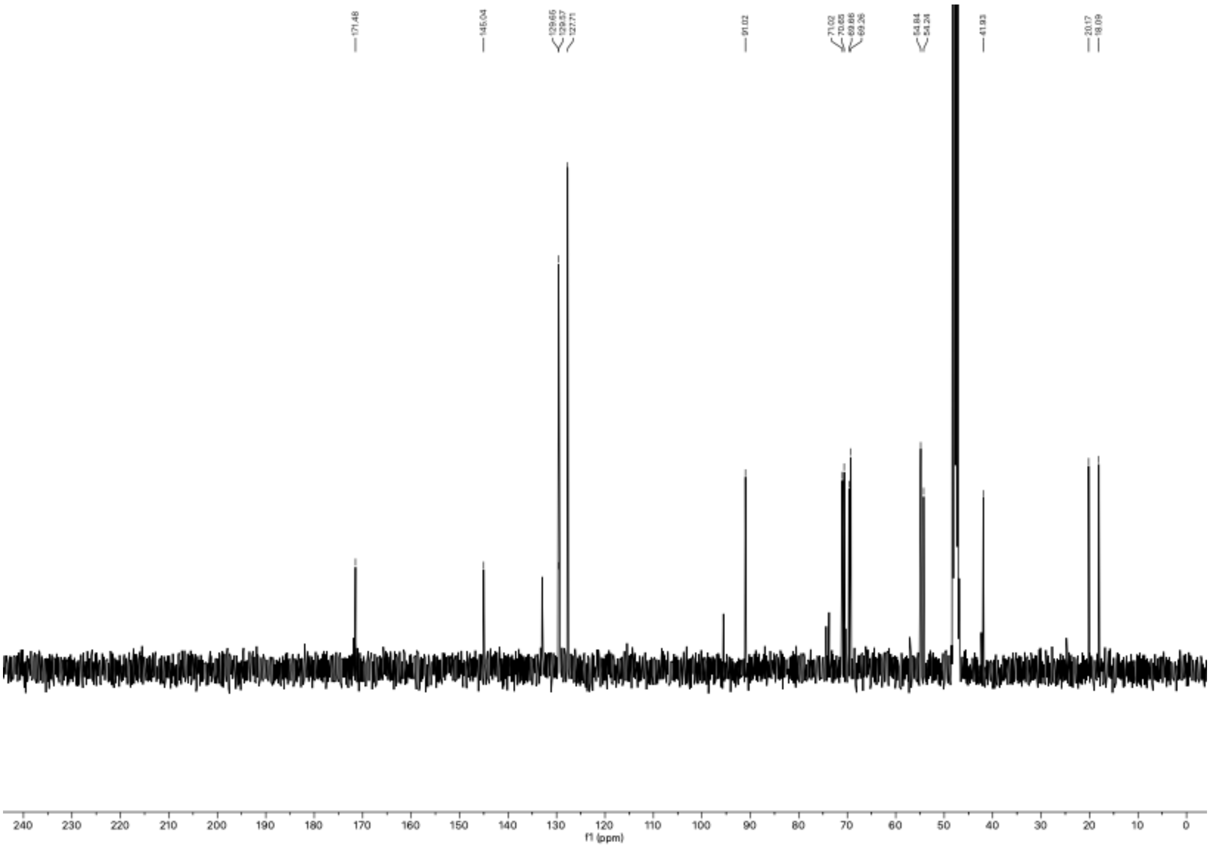

[1]H NMR, $CD_3OD$, 400 MHz

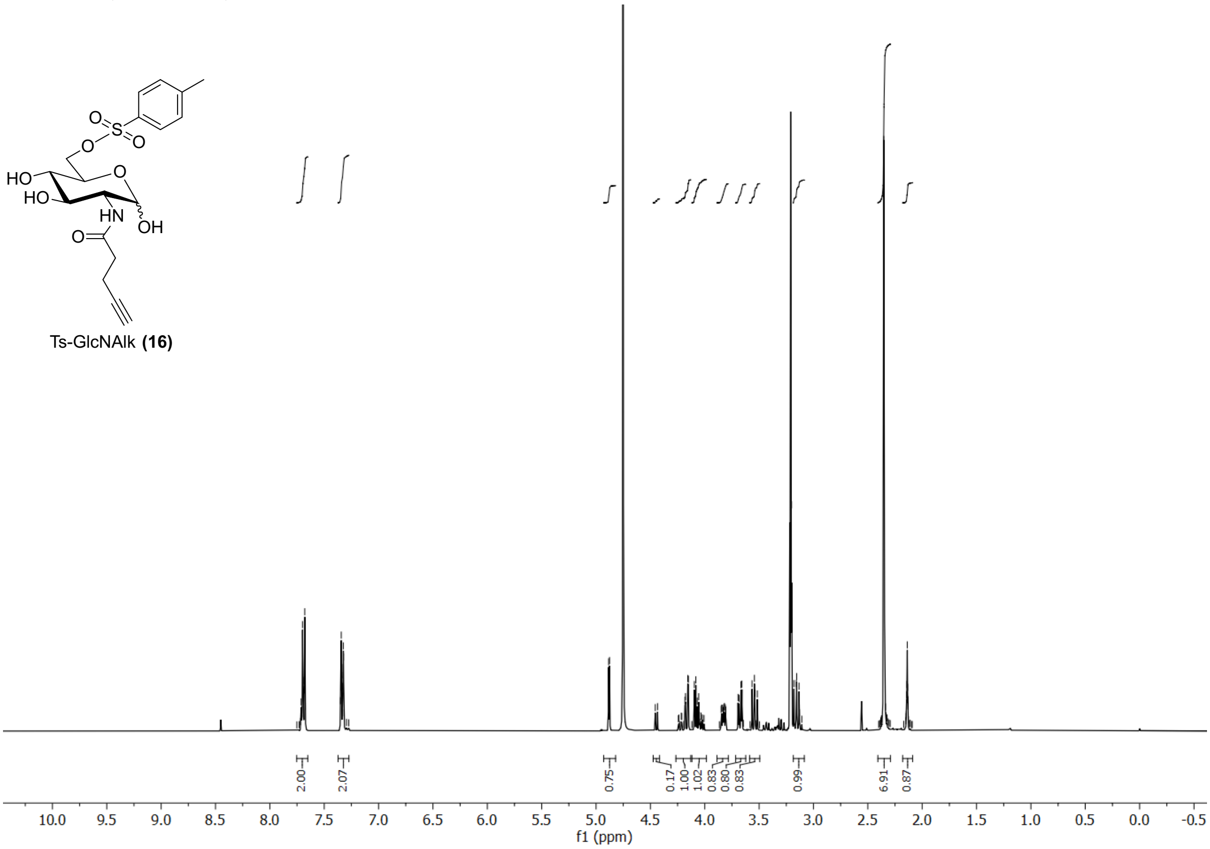


[13]C NMR, $CD_3OD$, 100 MHz

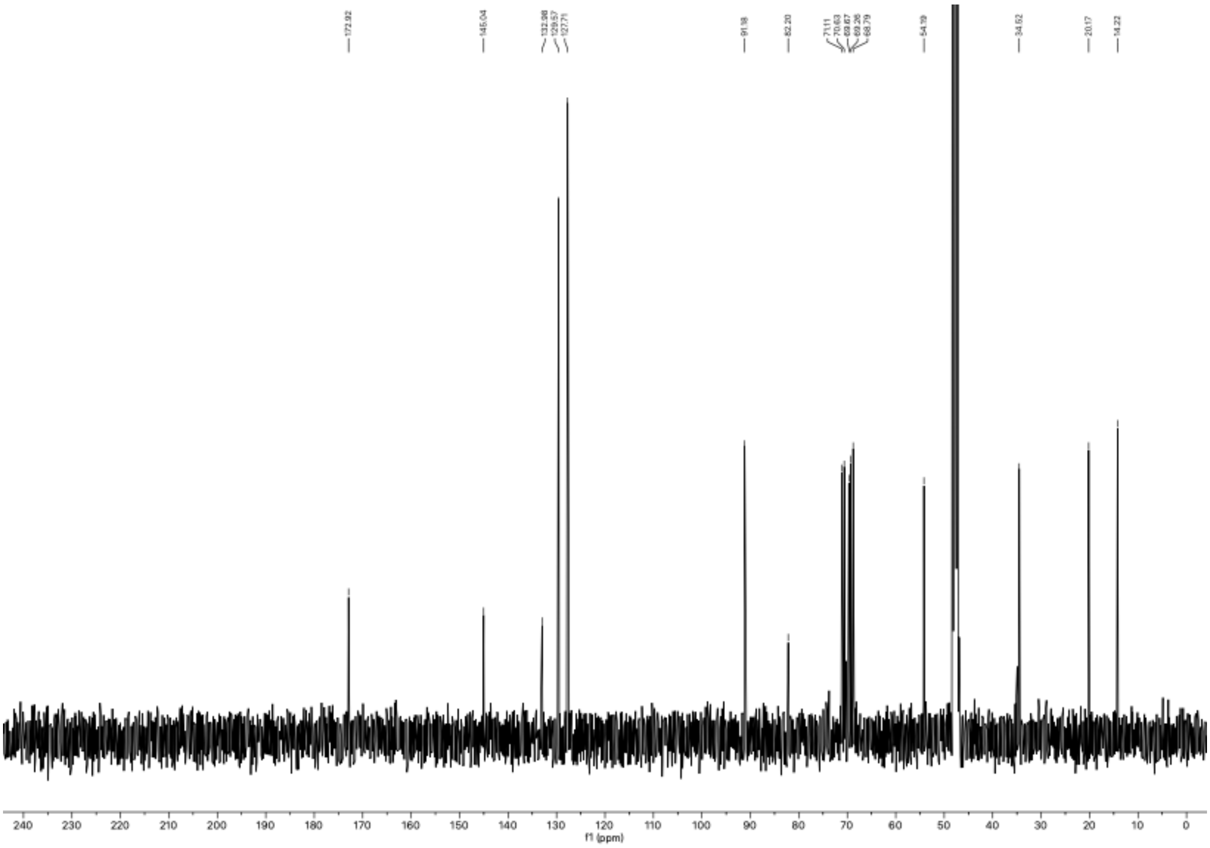

$^{1}H$ NMR, $CDCl_3$, 400 MHz

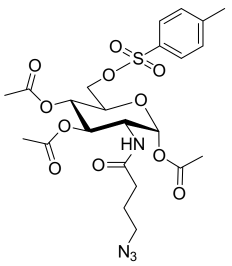

Ts-Ac$_3$GlcNButAz **(17)**

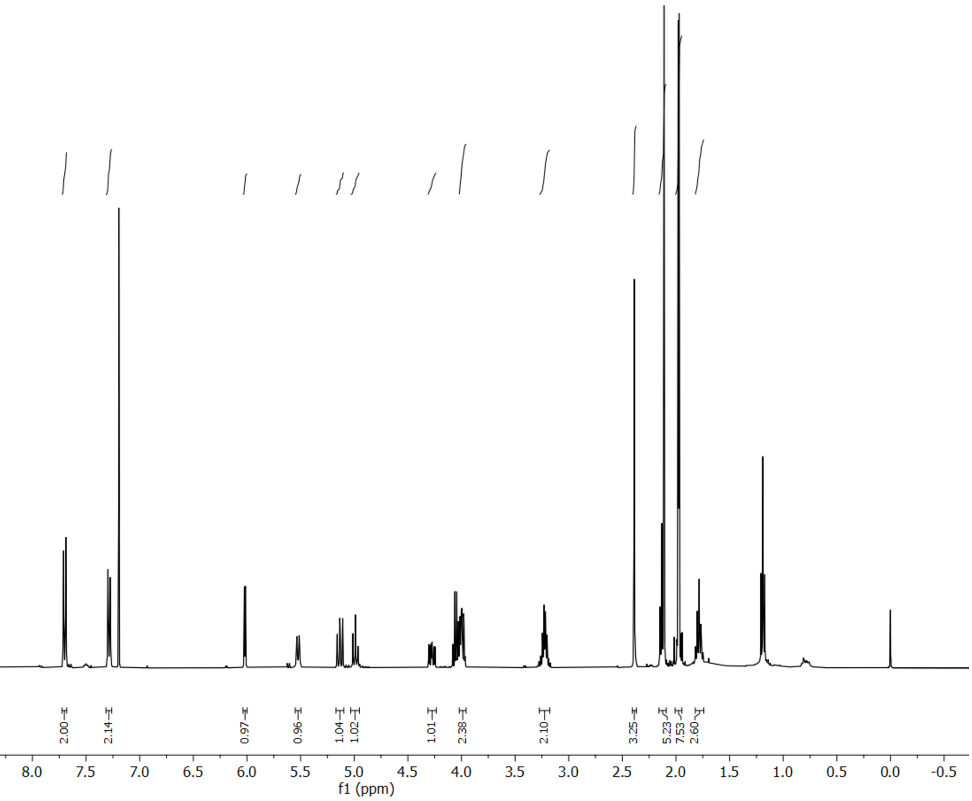

$^{13}C$ NMR, $CDCl_3$, 100 MHz

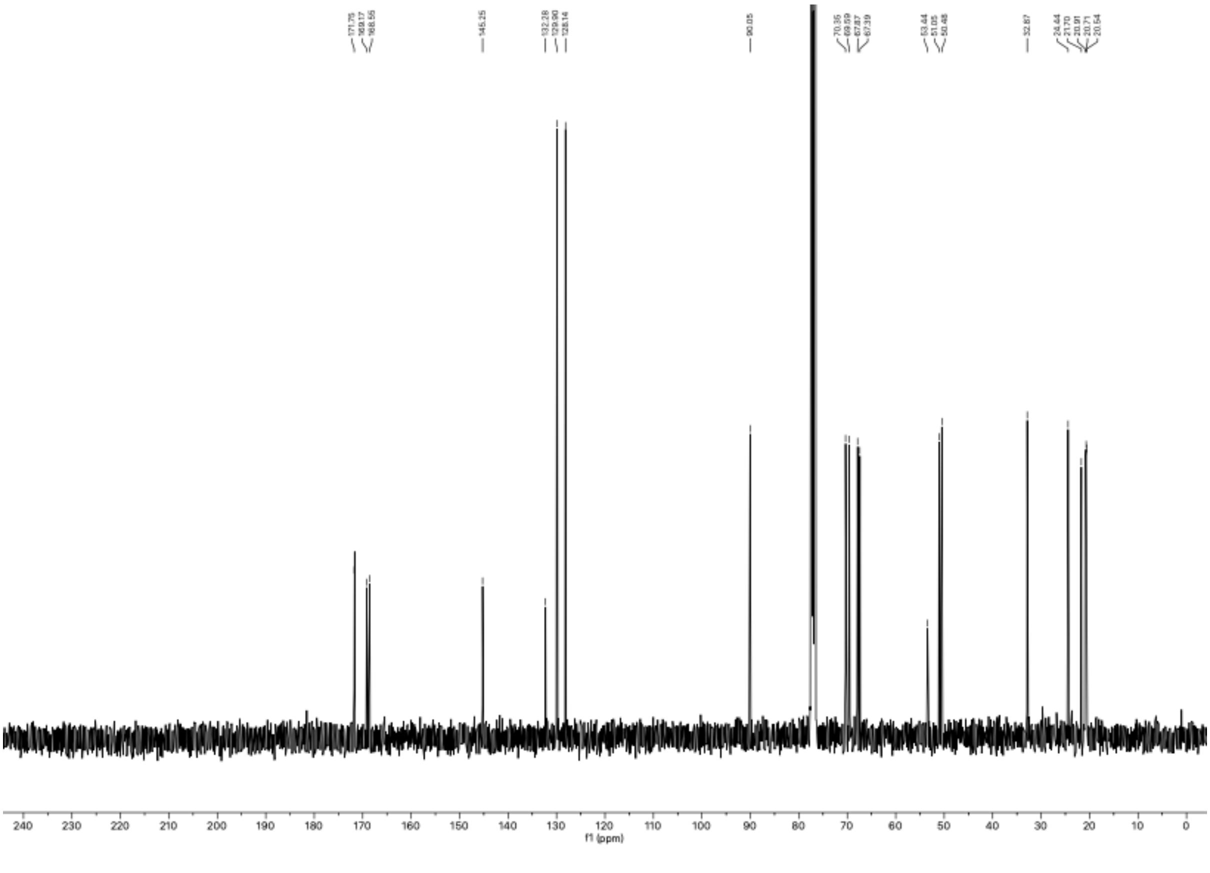

$^{1}H$ NMR, $CDCl_3$ with TMS, 400 MHz

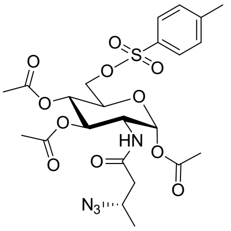


Ts-Ac$_3$GlcNPrAzMe(*S*) **(18)**

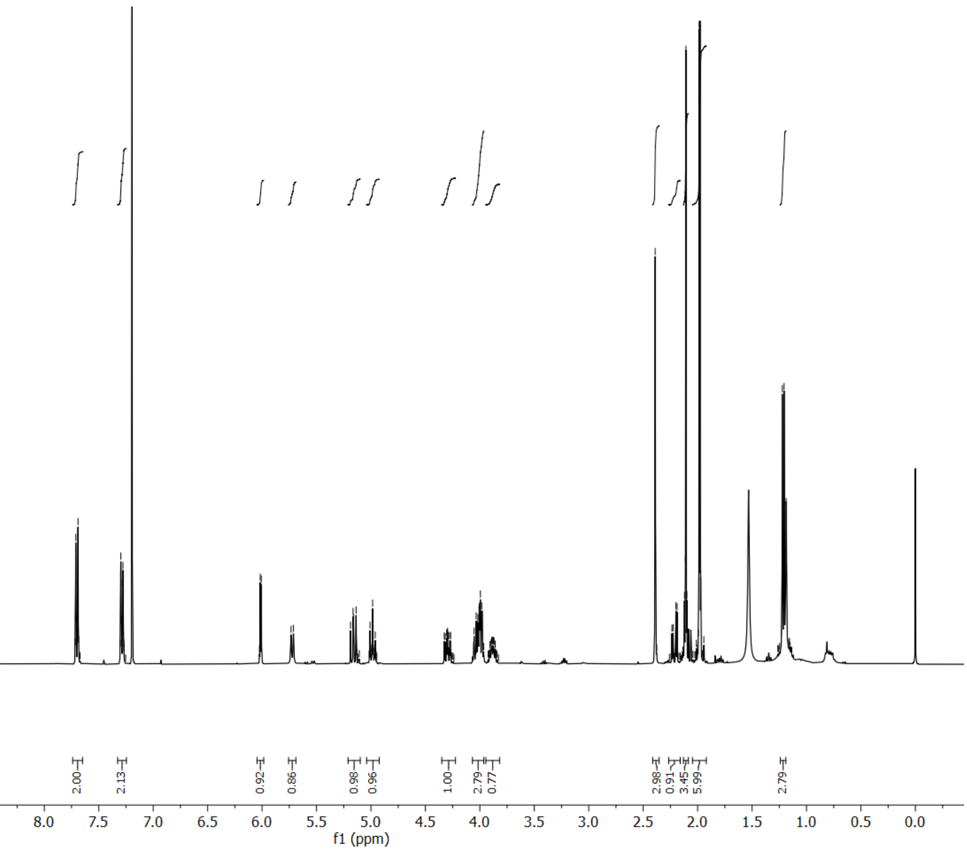


$^{13}C$ NMR, $CDCl_3$ with TMS, 150 MHz

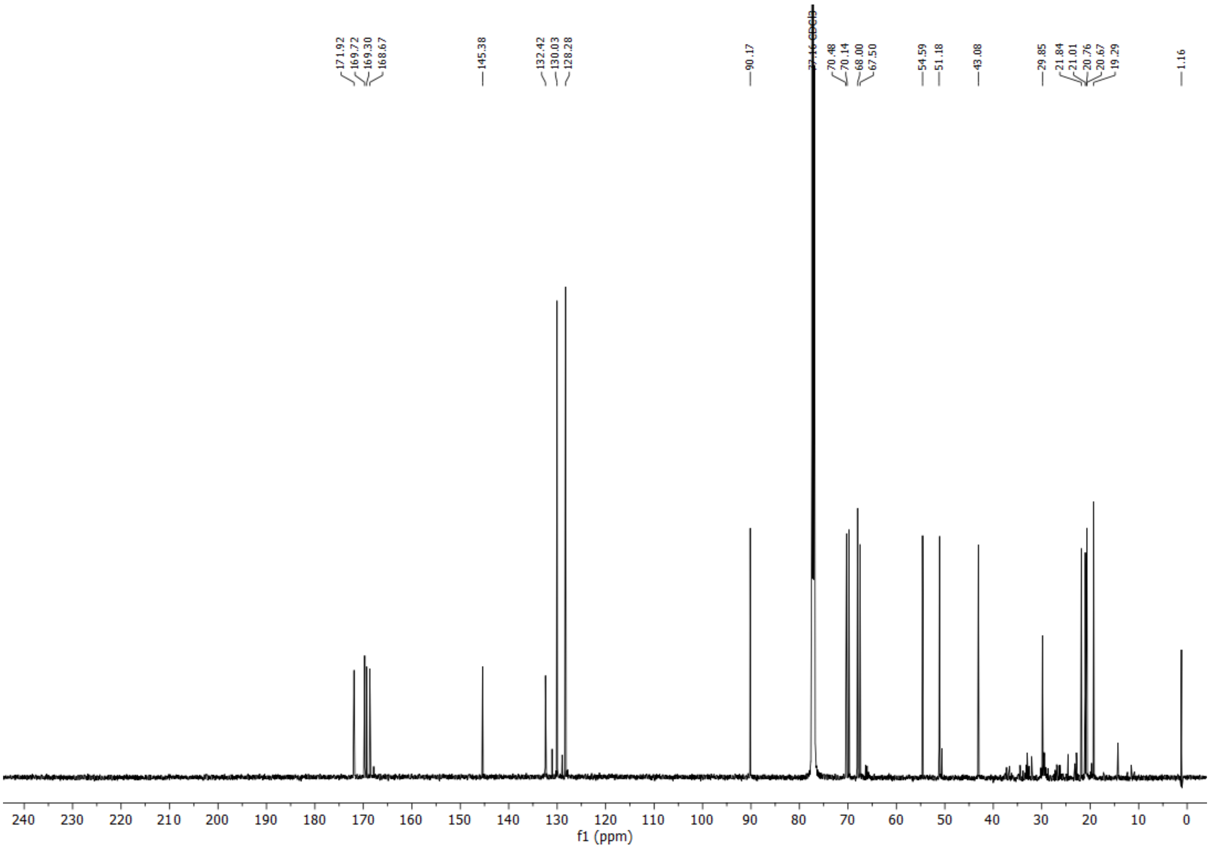

$^1$H NMR, $CDCl_3$, 400 MHz

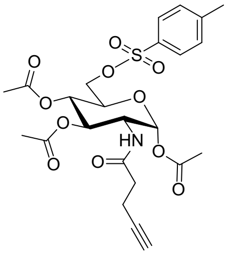

Ts-Ac$_3$GlcNAlk **(19)**

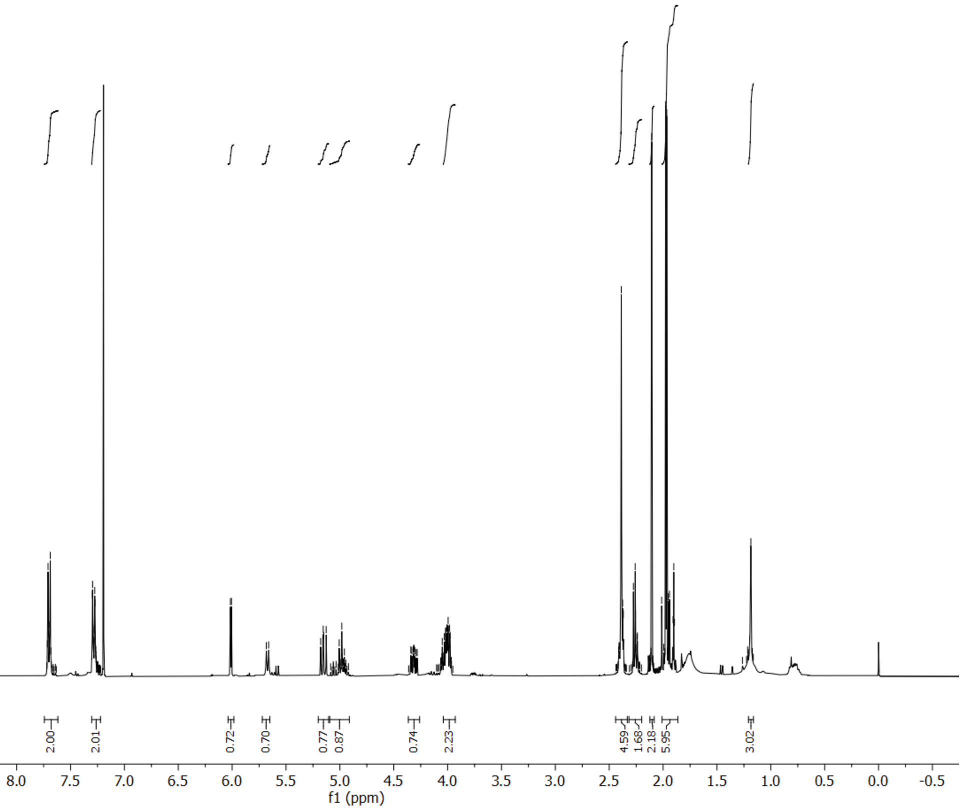


$^{13}$C NMR, $CDCl_3$, 100 MHz

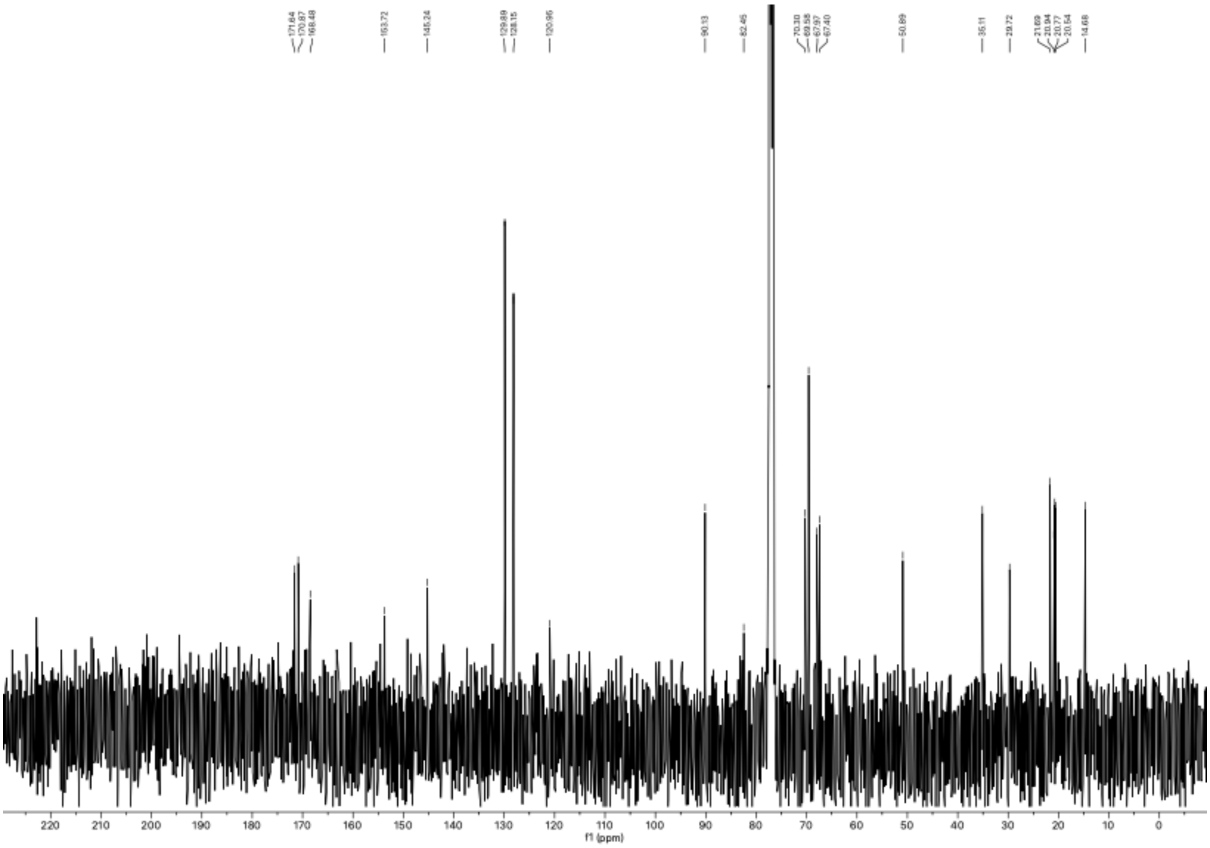
